# Hidden genetic diversity in 320 nearly-complete East Asian genome assemblies

**DOI:** 10.64898/2026.07.27.740875

**Authors:** Dongya Wu, Chentao Yang, Quanyu Chen, Mingyu Suo, Feifei Zhou, Anguo Liu, Dan Yu, Lei Nie, Ting Yang, Yanqing Sun, Junmin Han, Lili Yang, Qingyang Ni, Danyang Sun, Yanlin Lu, Lianting Fu, Yiqing Yang, Jie Yu, Jiajia Qi, Wei Dai, Xiangyu Yang, Lingxin Qiu, Dongxia Yang, Yonghui Jiao, Feng Zhou, Wenya Zhang, Fen Wang, Yihua Yang, Zhonghong Zeng, Zonghui Feng, Yan Chen, Yafeng Li, Yaheng Li, Shuyun Zhao, Aizhuan Long, Zhixue Wang, Qiye Li, Ruoping Zhao, Guo Ding, Qiong Wang, Ya Tuo, Jingjing Yu, Hui Li, Kai Liu, Yu Zhang, Xiaosong Yan, Dawa, Yanfei Zhang, Aoyue Bi, Guangji Chen, Sheng Hu Qian, Xin Li, Xupeng Bi, Jing Liu, Jiaqi Li, Kezhi Fu, Sirui Ye, Shichun Wang, Jialan Yang, Qi Zhou, Jingxin Jiang, Wenna Xu, Yifeng Liu, Aixia Liu, Jiajun Meng, Yunlong Zhao, Yu Zhou, Bo Hong, Chuanfeng Lin, Long Zhou, Feng Liu, Jiawei Yang, Yunqiu He, Xiaofei Yang, Xiaotao Wang, Chuanle Xiao, Longjiang Fan, Ammar Al-Baadani, Jian Chen, Aifu Lin, Wanlin Wang, Weijie Fang, Zihui Zhang, Yaoxi He, Yongyong Shi, Kai Ye, Qi Zhou, Bing Su, Jian Huang, Jian Xu, Kai Wang, Dan Zhang, Hefeng Huang, Songmin Ying, Tianhua Zhou, Yafei Mao, Guojie Zhang

## Abstract

East Asian populations, representing over 20% of the global population, remain critically underrepresented in human genomic studies, limiting our understanding of population-stratified genetic variation and its implications for health and disease. Here we present the first phase of the Asian Pan-Genome project (APG), comprising 320 nearly complete, fully phased haploid genome assemblies from 160 East Asian individuals. These assemblies achieve unprecedented quality, with an average contig N50 of 144.3 megabase pairs and an average quality value of 64.5. Leveraging these superior assemblies, we reveal previously uncharacterized diversity in human repeatome, including population-stratified patterns in centromere satellites and rDNA arrays. Compared to existing global human genome assemblies, the newly generated genomes supplement 152 million base pairs of novel sequences, 355 gene gains, 18,300 structural variation loci and 26 large euchromatic inversions missing from current human pangenomes. We perform population stratification analyses of structural variations, and further resolve the structural haplotypes of complex genomic regions such as Major Histocompatibility Complex and Survival Motor Neuron loci across global pangenomes, exemplifying tandem-duplicate and inversion-rich complex locus architectures in the human genome, respectively. This resource provides a critical foundation for human genetic studies, especially for East Asian populations, promoting more accurate variant discovery, reducing bias, and ultimately advancing the equity and efficacy of genomic medicine.

## Introduction

An important mission in current human genomics is to resolve the profound sampling and reference bias in genetic studies, which has created profound gaps in our understanding of the full spectrum of human genetic diversity and its implications for precision medicine (Fatumo *et al*., 2022; Corpas *et al*., 2025). The East Asian (EAS) superpopulation, comprising approximately 20% of humanity, harbors substantial genetic diversity shaped by complex demographic and adaptive history, yet remains underrepresented in comprehensive genomic resources. Large-scale population genomic projects like the 1000 Genomes Project (1KGP; The 1000 Genomes Project Consortium, 2015), the Human Genome Diversity project (HGDP; Bergström *et al*., 2020), and the GenomeAsia 100K Project (GenomeAsia100K Consortium, 2019), have provided foundational catalogs of genetic variation across global populations, including EAS groups. However, these studies were built primarily on short-read sequencing and linear mapping on non-Asian reference sequences, which inherently struggles to accurately resolve complex genomic regions like segmental duplications (SDs), tandem repeats and centromeres, which harbor abundant structural variations (SVs) and novel sequences. Consequently, functionally important variants, particularly those impacting gene regulation and complex disease risk in EAS superpopulation, may have remained “dark” or mischaracterized.

Recent advancements in long-read sequencing technologies and genome assembly algorithms have enabled the construction of complete telomere-to-telomere (T2T) genome assembly, revealing extensive genetic variations in previously inaccessible complex regions, exemplified by the first complete genome T2T-CHM13 and the first complete Han Chinese reference T2T-CN1 (Nurk *et al*., 2022; Yang *et al*., 2023). The linear mapping quality may benefit from using population-matched reference genomes to potentially improve variant calling (Yang *et al*., 2023). Furthermore, to overcome the limitations inherent to a single linear reference, long-read-based pangenome efforts, such as the Human Pangenome Reference Consortium (HPRC) and the Human Genome Structural Variation Consortium (HGSVC), are building more globally representative genomic resources (Ebert *et al*., 2021; Liao *et al*., 2023; Logsdon *et al*., 2025). The Chinese Pangenome Consortium (CPC) and the 1000 Chinese Pangenome project (1KCP) have further expanded EAS representation (Gao *et al*., 2023; Wang *et al*., 2026). Despite these remarkable breakthroughs, high-quality T2T-level assemblies, capable of fully resolving the most complex genomic regions such as peri/centromeres, ribosomal DNA (rDNA) arrays, long tandem repeats and recurrently rearranged regions, e.g., Survival Motor Neuron (SMN) locus, are still scarce across human populations, and EAS T2T assemblies remain severely insufficient in quantity.

To address these critical gaps, we launched the Asian Pan-Genome (APG) project, aiming to generate a comprehensive, T2T-quality pangenome resource across diverse Asian ethnic groups. Here, we present APG phase 1 (APGp1), comprising 320 nearly T2T and haplotype-phased genome assemblies from 160 EAS male individuals. This resource represents the largest collection of nearly complete genome assemblies for regional ethnic groups to date, providing an essential resource for understanding genome-wide population-stratified genetic diversity, improving the accuracy of complex variant calling, and advancing precision medicine for EAS communities. The high contiguity and completeness of our assemblies allow us to investigate previously inaccessible complex regions, revealing novel insights into diversity of repeatome, rDNA, centromere organization, and medically relevant genomic loci.

### Assembling 320 phased and nearly T2T EAS genomes

We sampled a total of 160 healthy male individuals of EAS ancestry from China and Korea in APGp1 through a coordinated effort across multiple regional centers (**Fig. 1a**). All participants provided fully informed consent for the collection of biological samples and the generation and academic sharing of genomic information, and declared being in good health. Principal component analysis (PCA) using single nucleotide polymorphisms (SNPs) from Next-Generation Sequencing (NGS) short reads suggested representativeness of these samples in the genetic diversity within EAS populations (**Supplementary Fig. 1**). We sequenced high-depth long reads per sample (**Fig. 1b**; **Supplementary Table 1**). On average, we generated 51.4-fold (×) Pacific Biosciences High-Fidelity (PacBio HiFi) sequencing reads, with individual coverage ranging from 39.0× to 65.7×, using either PacBio Revio or Sequel II platforms. Additionally, we obtained ∼130-fold Oxford Nanopore Technologies (ONT) sequencing reads, with a N50 length of 96.4 kilo-base pairs (kbp), including ∼62.0-fold ultra-long ONT reads (>100 kbp, read N50: 177.3 kbp). To facilitate chromosome-scale scaffolding, assembly polishing and quality assessment, we generated clean Hi-C data (∼188.6-fold) and complementary paired-end short reads (∼111.6-fold) for each individual (**Supplementary Fig. 2**). For 142 participants, we additionally sequenced parental genomes and yielded high-coverage MGI-seq short reads (∼115.4-fold) to enable high-accuracy assembly phasing (**Supplementary Fig. 2**).

**Figure 1.**
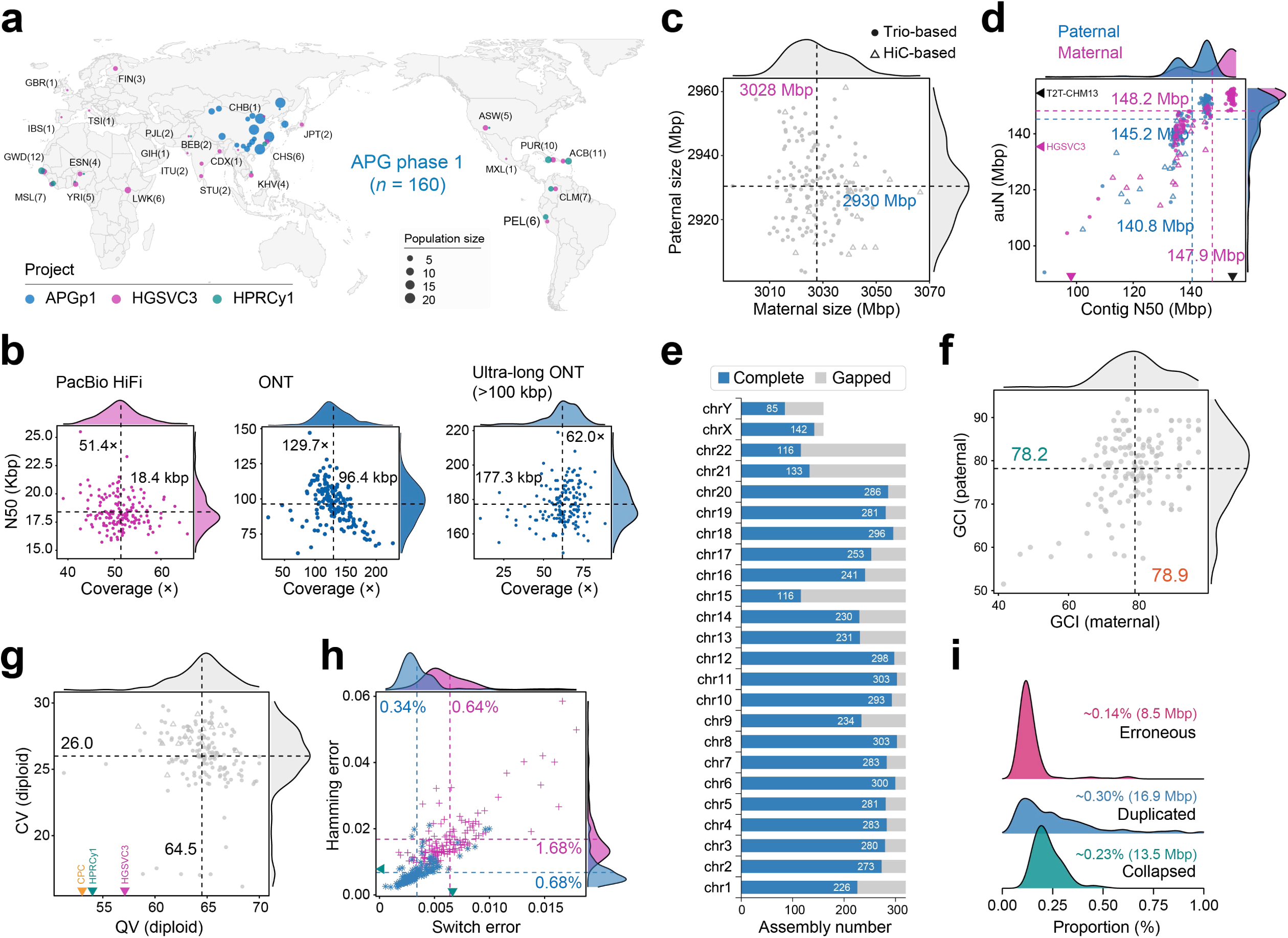
Sampling, sequencing and assembling APG phase 1 genomes. **a**, Genome sampling in APG phase 1 (APGp1), and HPRCy1 and HGSVC3 on a map of Earth. Samples from the 1000 Genomes Project (1KGP) are shown in subpopulations (three letter abbreviations) and samples from Genome in a Bottle (GIAB, HG002 and HG005), CEPH/UTAH pedigree panel (NA12329) and HapMap phase 3 (NA21309 and NA21487) are not included. ACB, African Caribbean in Barbados; ASW, African Ancestry in Southwest US; BEB, Bengali in Bangladesh; CDX, Chinese Dai in Xishuangbanna, China; CHB, Han Chinese in Beijing, China; CHS, Han Chinese South; CLM, Colombian in Medellin, Colombia; ESN, Esan in Nigeria; FIN, Finnish in Finland; GBR, British in England and Scotland; GIH, Gujarati Indians in Houston, Texas; GWD, Gambian in Western Division; IBS, Iberian populations in Spain; ITU, Indian Telugu in the UK; JPT, Japanese in Tokyo, Japan; KHV, Kinh in Ho Chi Minh City, Vietnam; MSL, Mende in Sierra Leone; MXL, Mexican Ancestry in Los Angeles, California; PEL, Peruvian in Lima, Peru; PJL, Punjabi in Lahore, Pakistan; PUR, Puerto Rican in Puerto Rico; STU, Sri Lankan Tamil in the UK; TSI, Toscani in Italy; YRI, Yoruba in Ibadan, Nigeria. **b**, Length and coverage statistics of long reads using PacBio HiFi and ONT sequencing platforms in APGp1. Mean values are shown in dashed lines. **c**, Genome sizes of phased maternal and paternal assemblies. Mean sizes are indicated by dashed lines. Assemblies phased by trio and Hi-C data are distinguished by dots and triangles, respectively. **d**, Assembly continuity index contig N50 and auN. The average values for assemblies from previous studies are displayed with solid triangles along the axis, where black, purple, green and yellow colors denote T2T-CHM13, HGSVC3, HPRCy1 and CPC, respectively. **e**, Assembly completeness per chromosome in APGp1 genomes. Blue bar represents the sample number with gap-free assembly for each chromosome. **f**, GCI scores of APGp1 assemblies. **g**, Base-accuracy quality values (QVs) and completeness values (CVs). **h**, Yak-reported switch and hamming error per assembly, indicating phasing quality. **i**, Summary of Flagger evaluation on APGp1 assemblies.

To generate complete and fully phased human genomes of individuals with parental NGS reads, we first constructed phased assemblies using hifiasm (Cheng *et al*., 2024) in trio-based phasing mode, integrating both PacBio HiFi and ultra-long ONT reads for individuals with available parental short-read sequences. We also generated phased assemblies using Verkko (Rautiainen *et al*., 2023) in trio-based phasing mode and hifiasm in Hi-C-binning mode to fill gaps in the initial hifiasm trio-based phased assemblies (**Supplementary Fig. 3**; **Methods**). For individuals lacking parental short reads (*n* = 18), the genome assemblies were phased with paired-end Hi-C reads using hifiasm. The Hi-C-phasing strategy exhibited comparable continuity and phasing accuracy with trio-based phasing strategy using high-coverage long reads (**Supplementary Fig. 4**). Local alignment of ONT reads against each draft assembly was conducted to fill gaps. Subsequently, we investigated the collinearity of each assembly against T2T-CN1 (v1.0) and T2T-CHM13 (v2.0) to identify and rule out large structural assembly errors by manual curation (**Supplementary Fig. 3**). Additionally, we performed a series of polishing steps to improve assembly base accuracy (**Methods**). Finally, we yielded a total of 320 phased EAS genome assemblies, and maternal haplotype assemblies averaged 3.028 giga-base pairs (Gbp; ranging from 2.996 to 3.066 Gbp) and paternal haplotype 2.930 Gbp (2.903 to 2.961 Gbp; **Fig. 1c**), with the difference attributed to sex chromosomes X and Y (**Supplementary Table 2**). Total assembly sizes across 160 diploid EAS individuals varied from 5.926 Gbp to 6.000 Gbp, with a mean size of 5.958 Gbp.

We conducted comprehensive quality assessments of our assemblies using multiple established tools (**Methods**). For contiguity, the average contig N50 value of the 320 phased assemblies reached 144.3 mega-base pairs (Mbp; 147.9 Mbp for maternal and 140.8 Mbp for paternal assemblies), substantially exceeding that of GRCh38 (57.9 Mbp) and recent long-read-based assemblies by HPRC year 1 (HPRCy1; ∼40 Mbp; Liao *et al*., 2023), CPC (∼35.6 Mbp; Gao *et al*., 2023) and HGSVC phase 3 (HGSVC3; ∼98.1 Mbp; Logsdon *et al*., 2025) (**Fig. 1d**). This superior contiguity was further confirmed by auN index (area under the Nx curve). On average, 18 chromosomes were completely assembled per haploid, though the acrocentric chromosomes 15, 21, 22 and Y remained challenging for gapless assembly (**Fig. 1e**; **Supplementary Table 3**). To evaluate completeness, we used Genome Continuity Inspector (GCI) workflow to detect potential assembly gaps lacking sufficient read support by PacBio HiFi and ONT reads (Chen *et al*., 2024). We detected an average of 15 assembly issues by using GCI, which were mainly distributed in rDNA (15.5%), and centromere regions (54.7%) of 8 chromosomes (excluding chromosome Y). Our assemblies achieved an average GCI score of 78.6, substantially higher than HPRCy1 (∼14.6) and HGSVC3 assemblies (∼39.6; **Fig. 1f**; **Supplementary Table 4**). Notably, 52 haploid assemblies exceeded the GCI scores of T2T-CHM13 assembly (87.04). For base-level accuracy, our diploid assemblies yielded an average quality value (QV) of 64.5, indicating <1 base error per Mbp, much higher than HPRCy1 (53.6; Liao *et al*., 2023) and HGSVC3 assemblies (56.1; **Fig. 1g**; Logsdon *et al*., 2025). VerityMap analysis of *k*-mer concordance between the assemblies and PacBio HiFi reads (Bzikadze *et al*., 2022) revealed minimal discrepancies (**Supplementary Fig. 5**). Regarding haplotype phasing accuracy, the average switch error was 0.64% and 0.34% for maternal and paternal assemblies, respectively, lower than the values of HPRCy1 assemblies (0.67%). Hamming error was estimated to be 1.68% and 0.68% for maternal and paternal, respectively (**Fig. 1h**). Multiple metrics also demonstrated the high completeness of our assemblies. The base-level completeness value (CV) averaged 26.0, indicating 99.75% of *k*-mers derived from raw reads were captured in the assemblies (**Fig. 1g**). Gene-level assessments using compleasm (Huang & Li, 2023) showed average completeness of 99.83% (maternal) and 96.74% (paternal) (**Supplementary Fig. 6a**). Aligning the annotated transcripts of T2T-CHM13 and T2T-CN1 onto each assembly demonstrated ∼99.16% of autosomal gene completeness (**Supplementary Fig. 6b**). Finally, we employed an integrative mapping-coverage-based pipeline Flagger (Liao *et al*., 2023) to identify unreliable assembled regions per assembly. The issued blocks detected in each diploid assembly accounted for 0.67% of the whole genome size on average, including 16.9 Mbp (0.30%) of duplicated, 13.5 Mbp (0.23%) of collapsed and 8.5 Mbp (0.14%) of erroneous segments per diploid genome (**Fig. 1i**; **Supplementary Fig. 7**). Taken together, these results indicate the high correctness, completeness, and contiguity of APGp1 assemblies.

### Repeatome diversity in EAS genomes

We leveraged these nearly T2T assemblies to characterize repetitive sequence diversity across EAS genomes, an analysis previously hindered by the limitations of draft genomes (**Methods**). Repetitive DNA sequences comprised an average of 1,496.9 Mbp or 52.27% of autosomal sequences in the 320 assemblies (**Fig. 2a**; **Supplementary Table 5**). Tandem satellites and simple repeats exhibited a relatively high coefficient of variation compared to other types of repeat elements (**Supplementary Fig. 8**). The distribution of simple repeats, particularly Variable Number Tandem Repeats (VNTRs) and Short Tandem Repeat (STRs), varied across chromosomes, but with 12 chromosomes showing relatively stable sizes with Coefficient of Variation (CV) < 0.05. In comparison, satellite repeats displayed a uniform variability pattern across all chromosomes. Notably, we observed substantial variation of SINE sequences in total cumulative length specifically on chromosome Y compared to other chromosomes, attributable to *Alu*Y, a member of SINE family, that constitutes basic repeat units within the heterochromatic region Yq12, one of the most dynamic regions across the human genome (Rhie *et al*., 2023).

**Figure 2.**
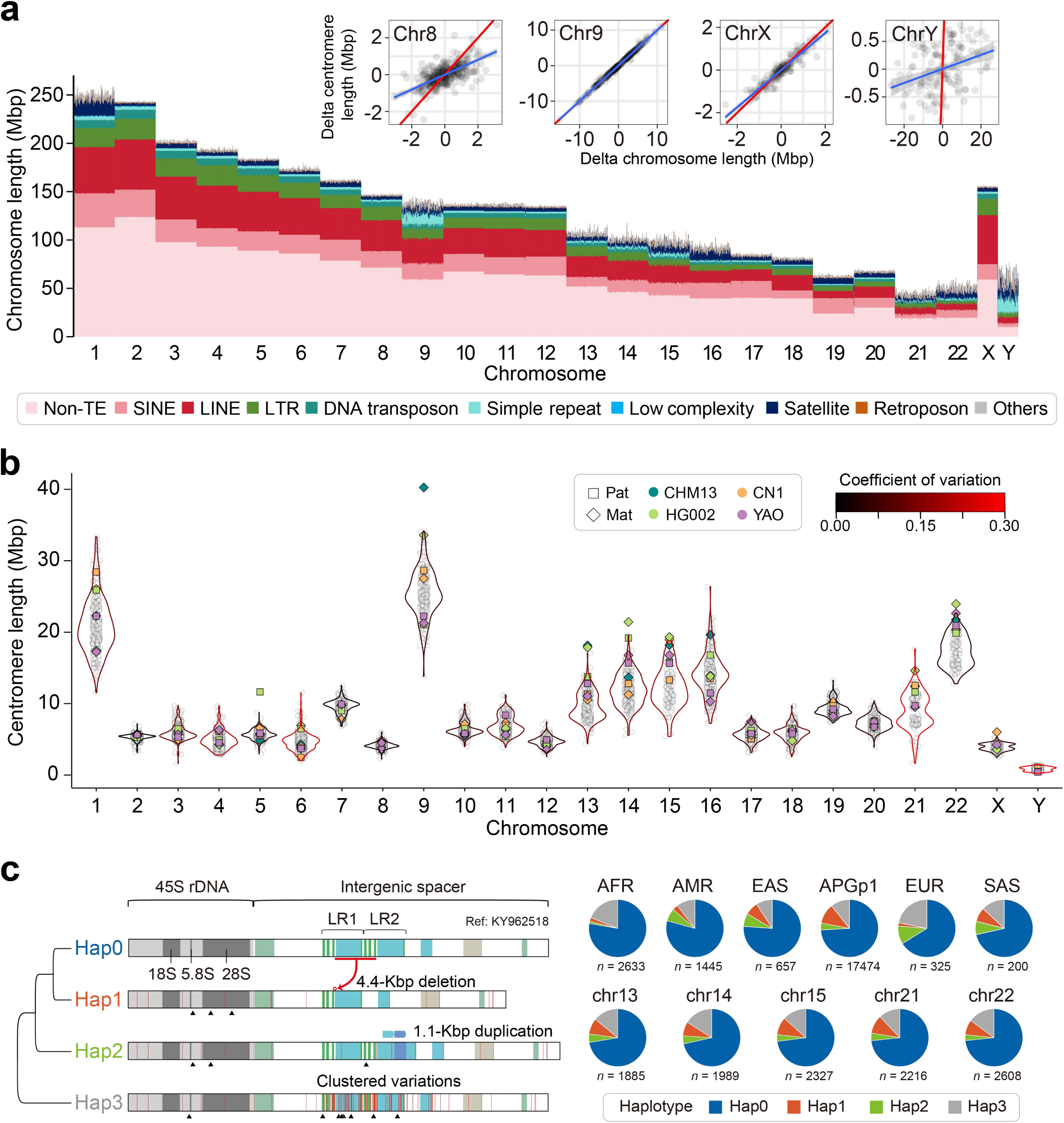
Repeatome diversity in EAS genomes. **a**, Repetitive element lengths across all human chromosomes. Top right, correlation plots of length variations between chromosomes and centromeres on chromosomes 8, 9, X and Y. Red lines represent the *y* = *x* reference and blue lines depict fitting linear regression curves. **b**, Length variation of complete centromeres in APGp1 assemblies. Previously released T2T assemblies are included (phased assemblies of T2T-CN1, YAO and HG002, and haploid assembly T2T-CHM13). Violin-plot curve colors represent the coefficient of variation (CV) in centromere lengths per chromosome. **c**, Genetic haplotypes of human rDNA elements. Left, Phylogenetic relationships and structural schematics of four rDNA haplotypes. rDNA is divided into the coding 45S rRNA genes (45S rDNA) and non-coding intergenic spacer. The long repeat (LR) comprises two copies (LR1 and LR2), each containing a long repeat, CT micro-satellites and Butterfly repeat. Variations are shown relative to the reference rDNA sequences (KY962518). Right, Haplotype frequency across human genomes from diverse populations and the five acrocentric chromosomes.

We next focused on the size variation of centromeres, which are mainly composed of satellites. Among all the assemblies, 6,312 out of 7,360 centromeres were fully gap-free, while the five acrocentric chromosomes and chromosome 9 required additional efforts to fill gaps (**Supplementary Fig. 9a**). On average, the resolved centromeric sequences occupied 7.33% of the whole genome (ranging from 6.31% to 8.20%). Size variations of complete centromeres differed among chromosomes, with chromosomes 4, 6, 21 and Y, displaying higher CVs than others (**Fig. 2b**). Within the centromeric regions, alpha satellites comprised the largest proportion (∼2.86% across the whole genome, from 2.47% to 3.60%) (**Supplementary Fig. 9b**). In contrast, the human satellites, Hsat1, Hsat2 and Hsat3, were predominantly found on specific chromosomes (**Supplementary Figs. 9c-9e**). Notably, we observed a pronounced shift in the distribution of the centromere sizes of chromosomes 9 in EAS assemblies toward shorter lengths (Mean±s.d.: 25.44±3.18 Mbp), compared to T2T-CHM13 (40.25 Mbp) and African ancestry genomes (AFR; 28.06±4.53 Mbp; *P* = 2.9×10^-5^, Wilcoxon test), primarily driven by the reduction of Hsat3 arrays (**Supplementary Fig. 9f**). At present, it remains difficult to distinguish whether such centromeric size divergence arises from neutral genetic drift or adaptive natural selection, owing to the shortage of well-established analytical frameworks to detect selective constraints acting on highly repetitive non-coding regions. The dynamics of centromeric sequence abundance largely determined the variations of sequence sizes across all autosomal chromosomes and chromosome X, similar to the model plant *Arabidopsis* (**Fig. 2a**; **Supplementary Fig. 10**; Lian *et al*., 2024). Exceptionally, non-centromeric repetitive sequences on chromosome 8 contributed significantly to chromosome size variation, attributed to extensive structural rearrangements of innate immunity-associated β-defensin gene clusters (8p23.1; Logsdon *et al*., 2021).

Located within the centromeric regions of five acrocentric chromosomes, rDNA arrays represent the most challenging regions to assemble completely thus far. Across the 320 assemblies, we assembled a total of 17,474 intact rDNA copies with intact 45S coding region and non-coding intergenic spacer (IGS; **Fig. 2c**). By combining 5,260 intact rDNA copies from T2T-CHM13, HPRCy1 and HGSVC3, we clustered the global rDNA sequences into four distinct haplotypes based on specific variations (**Fig 2c**; **Supplementary Fig. 11**; **Methods**). Compared to the reference sequence of human rDNA (KY962518), Hap0 displayed limited variations and accounted for 74.8% of all the intact rDNA elements. Hap2 (4.0%) was characterized by a ∼1.1-kbp duplication in the CT-rich sub-region of Butterfly/Long Repeat 2 (LR2; **Fig. 2c**). In contrast, Hap1 (9.5%) contained one ∼4.4-kbp deletion in the LR region, which has been reported in the T2T-CN1 genome (Yang *et al*., 2023). As the most ancestral haplotype of human rDNA repeats, Hap3 (11.8%) did not show visible consensus SVs but exhibited a series of highly linked SNPs in the LR and its flanking region compared to the reference sequence (**Supplementary Fig. 11**). These haplotypes were differentiated by PCA using SNPs, confirming the validity of our categorization (**Supplementary Fig. 12a**). Differentiated frequency of rDNA types were observed among superpopulations, i.e., AFR, European (EUR), South Asian (SAS), American (AMR) and EAS ancestry groups (**Fig. 2c**). Hap3 accounted for ∼20% in AFR and EUR rDNA repeats, but only ∼10% in EAS genomes from both APGp1 and previous EAS assemblies. Hap1 repeats were more prevalent in Asian genomes (11.3%, 7.5% and 8.0% for APGp1, EAS and SAS genomes, respectively), compared to other superpopulations (AFR: 2.1%; EUR: 1.8%; AMR: 3.0%). No composition bias among the five acrocentric chromosomes was observed, consistent with the frequent recombination among acrocentric chromosomes (Guarracino *et al*., 2023) (**Fig. 2c**).

It was previously known that the rDNA array is composed of highly homogenized tandem clusters (Nurk *et al*., 2022; Hori *et al*., 2021; Rothschild *et al*., 2025). Here we examined the local homogenization of rDNA arrays, where rDNA repeats from the same structural haplotype tend to locate successively. Firstly, we validated the reliability of haplotyping in the assembled rDNA arrays by annotating rDNA elements of ultra-long ONT reads (>250 kbp, encompassing at least five rDNA copies; **Supplementary Fig. 13**; **Methods**). Among the 1,569 long assembled rDNA arrays (≥5 intact rDNA elements), 685 arrays (43.7%, covering 38.9% of rDNA copies) were found to be homogenized, where Hap0 was the most predominant (89.8%). The rest of the arrays showed widespread mosaic haplotype compositions, suggesting the presence of local haplotype heterogeneity (**Supplementary Fig. 14**). All these findings emphasized the need for continued efforts to assemble and fully decipher the complete rDNA arrays.

### Copy number variation of human protein-coding genes

We annotated the protein-coding genes using a hybrid annotation pipeline for each APGp1 assembly, capturing an average of 98.66% and 98.98% of annotated protein-coding genes from the GRCh38 and T2T-CN1 references per assembly, respectively (**Supplementary Figs. 15 and 16a**). By applying the identical annotation pipeline, EAS assemblies (*n* = 30) from HPRCy1 and HGSVC3 showed high consistency in gene copy number (CN) and diversity estimation with APGp1 (**Supplementary Fig. 16b**), demonstrating the comparability among the three pangenome datasets. Within APGp1, we identified ∼90 genes per assembly (excluding sex chromosomes) showing CN gains relative to GRCh38. Of the total 1,938 copy-gain genes, 355 were uniquely identified in APGp1 and absent from HPRCy1 and HGSVC3 genomes. Among these newly discovered copy-gain genes, 307 (86.5%) overlapped assembly issue regions in at least one assembly from HPRCy1 and HGSVC3, highlighting the improved assembly quality of APGp1 for capturing more gene copy gains, along with 48 copy-gain genes primarily owing to expanded population sampling depth (**Supplementary Fig. 17a**). Notably, merely eight of these amplified loci showed a frequency exceeding 1%, suggesting that the landscape of common functional gene CN variations is largely conserved across human populations. However, the remaining 321 singleton (present in one haplotype), and 26 rare (<1%) provide an essential resource for investigating the rare-variant architecture in EAS.

Among the APGp1 assemblies, 2,442 genes exhibited copy number variations (CNVs), of which 55.6% (*n* = 1,358) were singleton, and 12.6% (*n* = 307) were rare with frequency < 0.01. Genes with high CNV diversity (>0.26; top 2%) were highly enriched in the pathways related to sensory perception, defense response, and immunity, consistent with previous findings in HPRCy1 (**Supplementary Fig. 18**; Liao *et al*., 2023). These variable genes mostly were organized in tandem gene clusters, including *TAF11L*, *USP17L*, *PRAMEF* and *RASA4* (**Supplementary Fig. 19a**; **Supplementary Fig. 20**). Substantial CNVs across immunity-related gene clusters were observed, including the Major Histocompatibility Complex (MHC) region (43 of 128 genes from *HLA-F* to *HLA-DPB1*), the killer-cell immunoglobulin-like receptor (KIR) locus (15 out of 24 genes from *LILRB3* to *KIR3DL2*) and β-defensin locus on chromosome 8 (34 of 38 genes from *DEFB1* to *DEFB4A*; **Supplementary Fig. 19a**).

Notably, 74 genes have higher intra-population CN heterogeneity in EAS, as revealed by CNV diversity (difference > 0.1), implying potential selection on CNVs in EAS though rigorous evolutionary testing is still required to disentangle the potential effects of selection (**Supplementary Figs. 19b and 19c**; **Supplementary Table 6**). We also observed 146 genes displaying population-level expansion (mean CN difference > 0.1) in EAS than other populations. A notable example is the late cornified envelope (LCE) gene cluster, which encodes stratum-corneum proteins essential for innate cutaneous host defense. Prior studies have established that the deletion of *LCE3B* and *LCE3C* genes is a susceptibility factor for psoriasis (de Cid *et al*., 2009). We found this deletion occurred at a substantially lower frequency in AFR compared to non-AFR superpopulations (39.0% *versus* 61.0%; **Supplementary Fig. 21a**). Moreover, we identified another gene loss at *LCE1D*, which shows a markedly lower frequency in EAS than non-EAS superpopulations (2.0% for EAS *versus* 25.8% for non-EAS and 44.4% for EUR; **Supplementary Figs. 21b and 21c**). This difference aligns well with the observed frequencies of psoriasis, which are lowest in EAS and highest in EUR (Global Psoriasis Atlas; https://www.globalpsoriasisatlas.org).

Several CNVs were also observed in blood-related genes, including *RHD*, *RCHE*, *HBG1*, *HP*, *HPR* and *HBA2* (**Fig. 3**). Of particular interest is the *RHD* gene, which plays a crucial role in determining the Rh blood group system, the second most clinically significant blood group after ABO. The *RHD* gene exhibited the lowest deletion frequency in EAS (5.7%) among superpopulations. Additionally, we observed distinct population-enriched patterns in two adjacent haptoglobin-related genes *HP* and *HPR*. Specifically, the *HP* gene displays recurrent gene loss restricted in EAS assemblies (8 out of 350; *P* = 0.037, Chi-square test), whereas the *HPR* gene exhibits CN expansion solely in AFR (18 out of 100; *P* = 2.03×10^-19^; **Fig. 3**; **Supplementary Fig. 22**).

**Figure 3.**
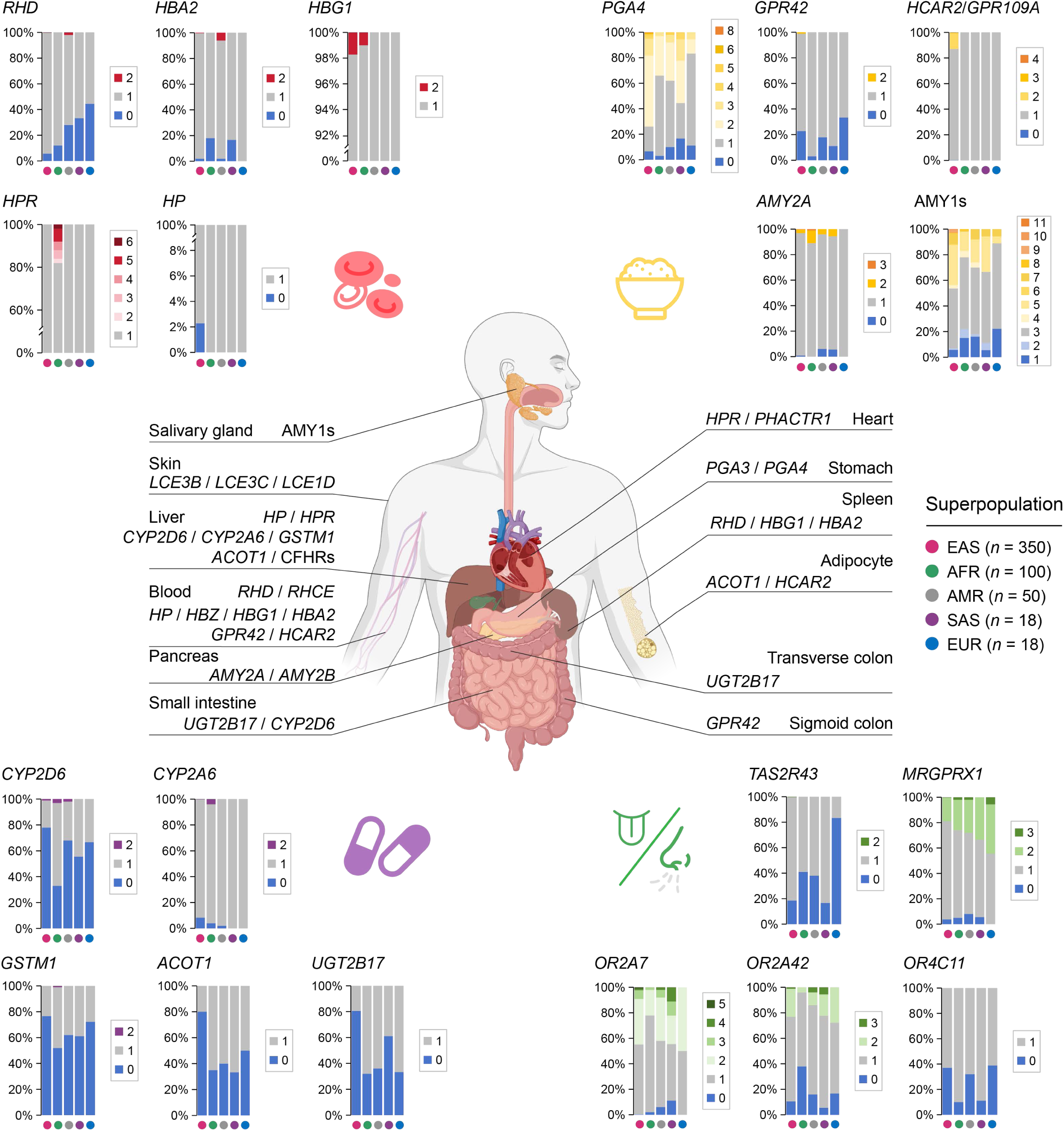
Copy number variations of human genes across super populations. Gene cases related to four categories are displayed, including blood, dietary, drug metabolism and sense receptor.

Population stratification in genes related to starch, protein or lipid digestion and metabolism was evident (**Fig. 3**). For the amylase-coding gene *AMY1*, while the most common haploid CN (3 copies) is largely shared across superpopulations, the overall distribution of *AMY1* CNs showed a significant shift toward higher copies in EAS (*P* = 3.22×10^-8^, one-side Wilcoxon rank-sum test), a pattern potentially linked to agricultural domestication of high-starch crops (Bolognini *et al*., 2024; Yilmaz *et al*., 2024). Similarly, gene *PGA4*, which encodes a protein precursor of the digestive enzyme pepsin and its serum levels has been used as a biomarker for atrophic gastritis and gastric cancer (Broutet *et al*., 2003), exhibited significant CN expansion in EAS with an average CN of 1.96 compared to 1.11 in EUR and 1.34 in AFR (*P* = 2.93×10^-10^ and 5.61×10^-5^, respectively, one-side Wilcoxon rank-sum test). This variation may reflect the long-term inter-population differences in dietary protein sources (animal-source or plant-derived; **Fig. 3**; **Supplementary Fig. 23a**). Gene *GPR42* encodes a G protein-coupled receptor activated by short chain fatty acids, playing a crucial role in whole-body energy homeostasis (Puhl *et al*., 2015). Approximately 23.4% of EAS haplotypes showed copy loss, contrasting sharply with AFR (3.0%; **Supplementary Fig. 23b**). Furthermore, *HCAR2*/*GPR109A*, which encodes a niacin receptor whose loss is associated with aging-related hepatic steatosis and obesity (Jadeja *et al*., 2019), showed CN expansion, with 12.9% of EAS assemblies (45 out of 350) containing at least two copies (**Supplementary Fig. 23c**).

Assessing the performance difference in drug metabolism and detoxification among individuals and populations is crucial for drug safety and efficacy (Margaret *et al*., 2002). We found several metabolic genes with reduced CNs in EAS, including *CYP2D6*, *CYP2A6*, *GSTM1*, *ACOT1* and *UGT2B17* (**Fig. 3**; **Supplementary Fig. 24**). Furthermore, population differentiation is particularly pronounced in sensory receptor genes (Nozawa *et al*., 2007), especially in olfactory receptor genes such as *OR2A7*, *OR2A42* and *OR4C11* (**Fig. 3**). In terms of taste perception, *TAS2R43* has been associated with coffee preference and caffeine bitter perception (Pirastu *et al*., 2014). Compared to EUR, both EAS and SAS exhibited lower deletion frequencies (*P* = 5.52×10^-10^ and 6.33×10^-5^, respectively; Chi-square test), supported by gnomAD frequencies (EUR: 0.514, *n* = 65,564; EAS: 0.209, *n* = 4,054; SAS: 0.230, *n* = 4,406).This stratification may associate with the regional differences in coffee and tea consumption habits. Furthermore, *MRGPRX1*, which encodes a G protein-coupled receptor that preferentially expresses in small-diameter primary sensory neurons mediating nociception and pruritus (Liu *et al*., 2009), displayed variable CNs, with EAS maintaining fewer copies (**Fig. 3**; **Supplementary Fig. 25**).

### Constructing and evaluating EAS pangenome graphs

To construct a comprehensive EAS pangenome graph, we integrated the 320 haplotype-resolved APGp1 assemblies, using Minigraph (MG; Li *et al*., 2020) and Minigraph-Cactus (MC; Hickey *et al*., 2024). The MC graphs demonstrated superior variation resolution, exhibiting node and edge sizes two orders of magnitude larger than MG, despite having smaller pangenome sizes (**Supplementary Table 7**). In the T2T-CN1-based APGp1 MC graph, the assemblies collectively supplemented 319.5 Mbp of sequences that are absent from the reference. Frequency distribution of novel sequences showed that only 0.54 Mbp were core sequences present in >95% of the assemblies, while 66.31 Mbp appeared commonly across the population (5% to 95%). The discovery of 60.68 Mbp of rare sequences (occurring in <1% but ≥ 2 assemblies) and 125.77 Mbp of singletons highlighted the depth of genetic diversity within the EAS superpopulation (**Fig. 4a**). Further analysis on the graph suggested that the pangenome growth curve closely followed Heaps’ Law under nonlinear least squares regression (**Fig. 4a**). This graph demonstrated unprecedented scale and complexity, with a total base length of 3.42 Gbp, comprising 87.9 million nodes and 120.9 million edges, exceeding the size of previous constructs, including the EAS-only MC graph derived from HPRCy1 and HGSVC3 (30 haplotypes, with a total length of 3.20 Gbp, including 41.7 million nodes and 57.4 million edges with T2T-CN1 as the backbone) and the publicly available CPC MC graph (124 haplotypes, sized 3.28 Gbp, with 64.1 million nodes and 89.2 million edges), highlighting the significance of incorporating more personal genomes in pangenome graph construction (**Supplementary Table 7**). Notably, we observed an openness value of 0.434, lower than that of the pangenome graph derived from HPRCy1 and HGSVC3 assemblies (0.523), in line with the relatively lower genetic diversity within EAS superpopulation compared to global populations (**Supplementary Fig. 1b**).

**Figure 4.**
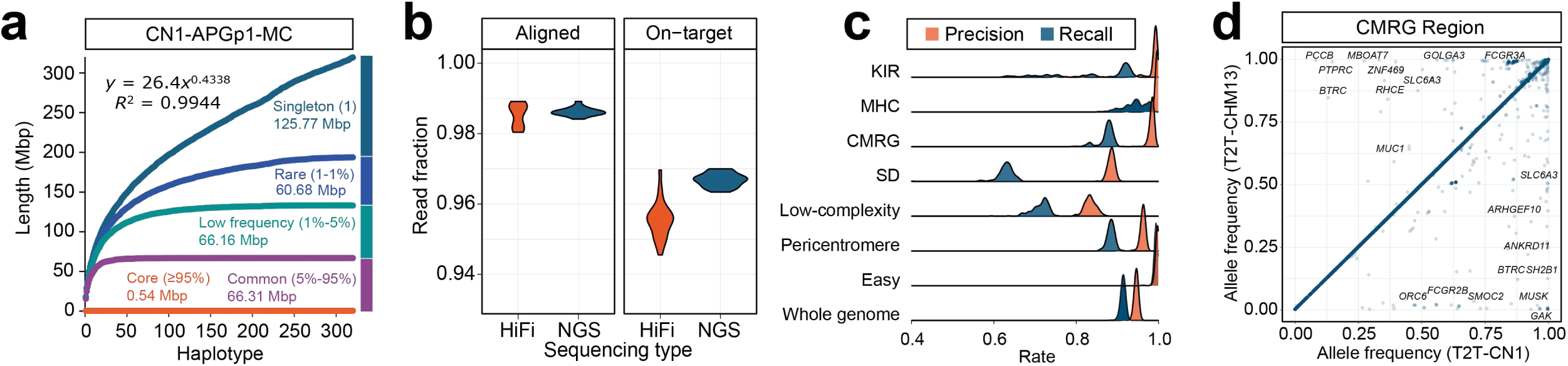
Human pangenome graph construction and evaluation. **a**, Growth curves of non-reference sequences in the Minigraph-Cactus (MC) pangenome graph from APGp1 assemblies (*n* = 320). Five sequence categories are defined according to the frequency. **b**, Alignment performance of PacBio HiFi and NGS reads from APGp1 samples against APGp1 MC graph. **c**, Precision and recall of autosomal variants by mapping NGS short reads against APGp1 MC pangenome graph and calling using DeepVariant, relative to graph-decomposed variant calls. **d**, Allele frequency of shared variant calls in APGp1 genomes using the T2T-CN1 and T2T-CHM13-based MC pangenome graphs, respectively.

We next assessed the sequence representativeness of the T2T-CN1-based APGp1 MC graph, by mapping NGS short reads and PacBio HiFi reads from each APGp1 individual onto the graph using Giraffe (Sirén *et al*., 2021) and GraphAligner (Rautiainen & Marschall, 2020). The results demonstrated exceptional performance, with alignment rates exceeding 98.5% for both NGS and PacBio HiFi reads (**Fig. 4b**). Furthermore, 96.68% of aligned NGS reads and 95.54% of aligned PacBio HiFi reads successfully targeted their haplotypic paths (on-target), despite that the unreliable coverage rates (off-target edges with >0.1 times the expected mapping depth) were higher in repetitive regions (**Fig. 4b**; **Supplementary Fig. 26**). We subsequently assessed the performance of variant calling based on the alignments of NGS reads on the APGp1 graph using DeepVariant (Poplin *et al*., 2018), by comparing to the variants decoded from the graph. An average recall rate of 91.4% and precision rate of 94.6% were yielded across the whole genome (**Fig. 4c**). ‘Easy’ regions, which cover 74.4% of T2T-CN1, exhibited superior performance with high precision (99.7%) and recall rate (99.4%; **Fig. 4c**). For the challenging medically relevant genes (CMRGs), we observed an average recall rate of 87.3% along with a precision rate of 98.2%. Of particular notes were the immunity-related loci, MHC, with a precision and recall of 99.6% and 94.3%, and KIR, showing precision and recall rate of 98.4% and 84.9%, respectively (**Fig. 4c**).

Considering potential reference bias, we compared the MC graphs of APGp1 assemblies constructed using T2T-CHM13 and T2T-CN1 as reference backbones, respectively. The T2T-CHM13-referenced graph exhibited a slightly larger size but fewer nodes and edges (**Supplementary Table 7**). Cross-graph coordinate liftoff of decomposed variant calls revealed that the discordance was highly localized. Structurally repetitive, divergent, or inverted regions constituted major hotspots for liftoff failures, including the two largest inversions at 16p13.11-p12.3 and 8p23.1 between the two references (**Supplementary Table 8**; **Supplementary Figs. 27 and 28**). These reference-specific discrepancies highlight the necessity of exercising caution when interpreting variant callsets in complex regions, as they are confounding factors stemming from the inherent limitations of graph heuristics in repetitive regions, real structural divergence between reference genomes, and coordinate-shifting artifacts during cross-reference lifting. Conversely, within genomic “easy” regions, the variant profiles from both graphs were nearly identical, with 99.1% of calls being pairwise lifted (**Supplementary Table 8**). Generally, the liftable variants across the genome displayed consistent allele frequencies (AFs; *R*^2^ = 0.991), with a small proportion (1.53%) showing AF differences greater than 0.1, which were primarily concentrated in complex regions, including 313 small variants situated within 55 CMRGs (**Fig. 4d**; **Supplementary Fig. 29a**). For instance, 131 out of all 178 shared variants between the two graphs showed AF difference >0.1 within the *FCGR3A/B* region (1q23.3), largely due to gene CNV between the two graph references (**Supplementary Fig. 29b**). This region is associated with the susceptibility of systemic autoimmune disease (Zhou *et al*., 2010) and the copy number of *FCGR3A/B* genes in T2T-CN1 represents the most common type in about 80% of human genomes. To investigate the impact of reference backbones on graph-based read mapping, we aligned short reads from EAS individuals, HG006 and HG007 from Genome in a Bottle (GIAB), and PacBio HiFi reads from EAS individuals YAO (He *et al*., 2023) and Chinese Quartet (Jia *et al*., 2023), against the T2T-CN1 and T2T-CHM13-based graphs. The T2T-CN1-based graph demonstrated slightly higher mapping rates with an increase of ∼0.05% and ∼0.24% for NGS and PacBio HiFi reads, respectively, and lower rates of unreliable mapping, compared to the T2T-CHM13-based graph (**Supplementary Fig. 30**).

By integrating previously released human genome assemblies in HPRCy1 and HGSVC3, we further constructed a MC pangenome graph for global populations (GLOBALp1), encompassing a total of 540 haplotype-resolved assemblies. The GLOBALp1 graph achieved a total length of 3.61 Gbp with 156.6 million nodes and 216.0 million edges (**Supplementary Table 7**). Notably, the APGp1 assemblies contributed 152.3 Mbp of novel non-reference sequences besides those present in the HPRCy1 and HGSVC3 assembly graphs (**Supplementary Fig. 31**). Given that pangenome graphs representing diverse human ancestries can potentially impact read mapping and variant calling, we sought to employ Mapping Rate (MR), High-Confidence Mapping Rate (HCMR), and Perfect Mapping Rate (PMR), to assess NGS alignment performance for diverse populations from HGDP (Bergström *et al*., 2020) across six MC pangenome graphs, including two previously released by HPRCy1 and CPC (**Supplementary Fig. 32**; **Methods**). The dynamic between MR and PMR highlights a structural trade-off in pangenome graph mapping. MR primarily reflects overall alignment scalability and general sequence accessibility; a more streamlined graph topology, achieved by filtering out low-frequency alleles or utilizing a highly curated set of representative discovery samples, maximizes MR by mitigating graph complexity. However, this architectural simplification inadvertently compromises downstream genotyping sensitivity for rare or lineage-specific variants. Conversely, PMR quantifies the proportion of reads aligning with zero mismatches or indels, making it highly sensitive to the presence of exact local haplotypes within the graph backbone. Graphs that retain substantial allelic diversity, such as GLOBALp1, achieve significantly elevated PMR by providing exact sequence matches for population-stratified alleles, albeit at the expense of increased topological complexity. Beyond the efficiency-diversity balance reflected by MR and PMR, HCMR acts as a more pragmatic indicator for practical graph applicability. This metric prioritizes high-quality reliable alignments without mandating full sequence identity, thereby better reflecting the contribution of ancestry-matched homologous sequences.

While T2T-CHM13-referenced HPRCy1 graph demonstrated superior overall MRs across populations, APGp1 graphs showed notably better performance for EAS and Oceania populations in terms of HCR and PMR (**Supplementary Fig. 32**). These findings demonstrate that while an optimized global graph layout facilitates pan-ethnic alignment, the strategic injection of ancestry-specific sequences is important for maximizing high-confidence read recovery within specific lineages.

### Loss-of-function variants in the human genome

Loss-of-function variants are a critical subset of genomic variants that can have severe effects on protein products and determine gene essentiality (Sun *et al*. 2024; MacArthur *et al*. 2012). Using the variant calls from the global MC pangenome graph, we annotated autosomal small putative loss-of-function variants (pLoF, ≤ 50 bp) per assembly and individual (**Methods**). Our initial validation against the well-characterized HG002 genome from GIAB (v4.2.1; Wagner *et al*. 2022) demonstrated the robustness of our pangenome graph decomposition approach. While we identified 195 shared pLoF variants with GIAB’s reported 225 variants, we also discovered 72 novel pLoF variants, of which 38 were independently confirmed through PacBio HiFi and NGS read alignments (**Supplementary Table 9**). The discrepancies with GIAB’s calls were primarily due to specific technical challenges, including MNPs (*n* = 7), miscalling with short reads (*n* = 9) and assembly errors of GRCh38 reference (e.g., *FCGBP*; *n* = 9; **Supplementary Table 9**). Given that the genomes of HPRCy1 and HGSVC3 were sequenced from cell lines with potential long-term mutation accumulation, they displayed an abnormal pLoF variant spectrum and were thus excluded from further analysis. In APGp1 individuals, we observed a consistent pattern of pLoF variants comparable to that in HG002. Each assembly harbored a median of 195 pLoF variants, while diploid individuals carried approximately 293 variants. These variants predominately manifested as frameshift (61.7%), followed by stop-gained (24.7%), and essential splice variants (13.6%), which disrupt splicing-donor/acceptor sites (**Fig. 5a**). Notably, our pangenome-based approach identified an average of 75 additional pLoF variants per individual that were missed by traditional NGS-based methods (Sun *et al*. 2024; Karczewski *et al*. 2020; MacArthur *et al*. 2012), with ∼30% of these variants actually present in the gnomAD database but previously unclassified as pLoF due to quality control limitations (**Fig. 5b**; **Supplementary** Table 10).

**Figure 5.**
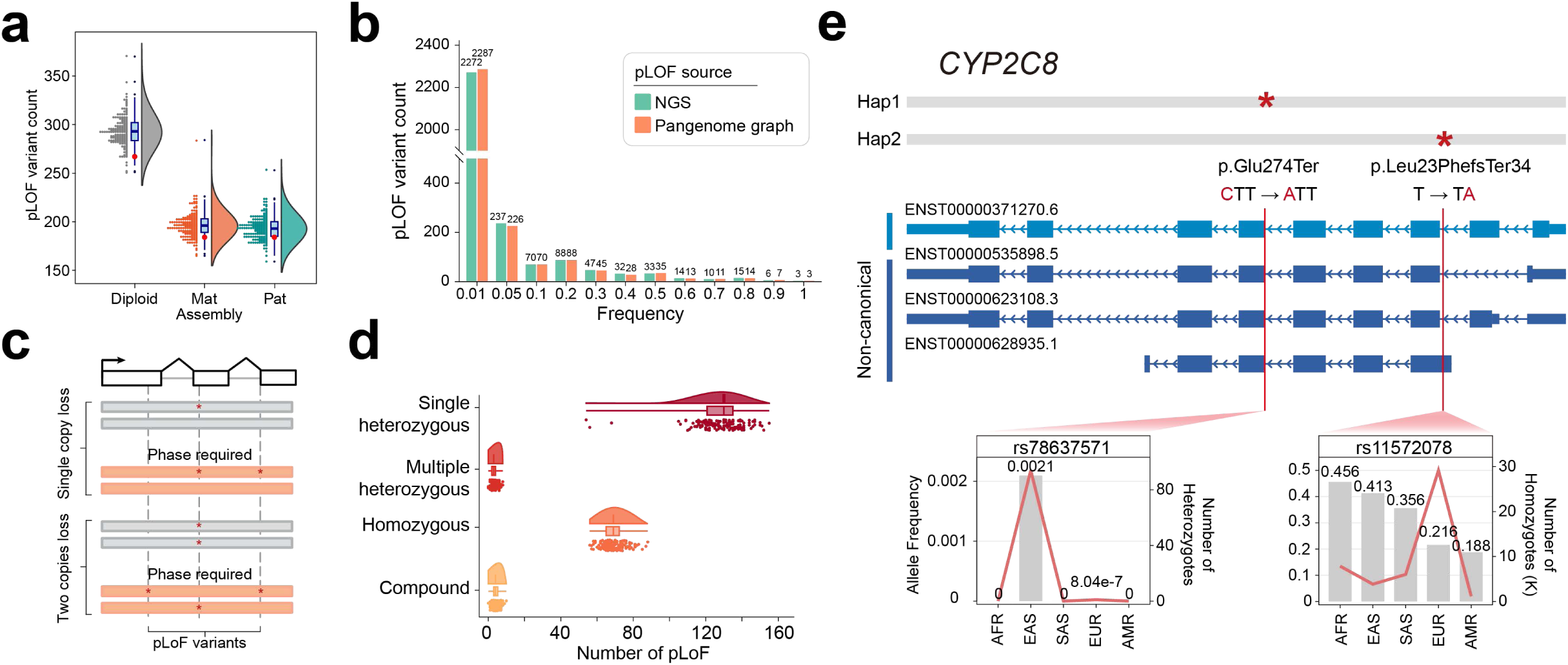
Loss-of-function variations in human genomes. **a**, Count distribution of small pLoF variants in APGp1 haplotype-resolved assemblies and individuals. The red dots represent the pLOF calls in HG002. **b**, Allele frequency of consistent pLoF variants across pangenome-based and NGS-based variant calls. **c**, Gene-loss types with phased pLoF variants. **d**, Number of pLoF variants classified as four categories in APGp1 samples. **e**, Example of a compound gene loss event in *CYP2C8* featuring a EAS-specific LoF variant. AF and the number of heterozygotes / homozygotes were retrieved from GnomAD (v4.1). pLoF variants are denoted as red asterisks.

Complete and haplotype-resolved genome assembly allows us to further investigate the phasing patterns of pLoF variants (**Fig. 5c**). By examining 2,515 affected genes across APGp1 genomes, we found that each individual had a median of four and three genes with multiple heterozygous (all in *cis*) and compound (in *trans*) pLoF phasing states, respectively, significantly fewer than the counts of genes with a single heterozygous or homozygous pLoF (127 and 69; *P* < 2.2×10^-16^, Wilcoxon rank-sum test; **Fig. 5d**). A total of 68 genes in APGp1 showed instances with compound pLoF variants, in which 34 genes were present in at least two individuals (**Supplementary Table 11**; **Supplementary Fig. 33a**). These genes displayed significant enrichment for high tissue-specificity in expression (*P* = 1.24×10^-4^, Fisher’s exact test) and weak evolutionary constraint (*P* = 4.14×10^-5^, Fisher’s exact test; **Supplementary Figs. 33b and 33c**).

Besides the combination of two common pLoF variants in the compound genes, 57.4% (39/68) of the observed compound genes had one common pLoF variant along with a rare or singleton pLoF variant (**Supplementary Table 11**). One notable example is gene *CYP2C8*, which encodes a liver-specific drug metabolizing enzyme (Backman *et al*., 2016). We identified a compound gene loss event involving a common frameshift variant (rs11572078, global AF = 0.24) and a rare EAS-specific stop-gained variant (rs78637571, AF_EAS_ = 0.002 in GnomAD) with no observed homozygotes (**Fig. 5e**; **Supplementary Table 12**). The high frequency and prevalent homozygotes for the common frameshift variant across populations suggest that this particular compound loss has limited negative consequences, consistent with our broader observation that compound pLoF variants in healthy individuals typically affect genes with tissue-specific expression patterns and dispensable functions, distinguishing them from pathogenic compound losses (MacArthur *et al*. 2012).

### Genomic diversity and divergence of SVs

Through a graph decomposition approach with a novel SV-pruning algorithm PanSVMerger (https://github.com/Asian-Pan-Genome/PanSVMerger), we have uncovered an unprecedented view of SV (>50 bp) diversity across human populations (**Supplementary Table 13**; **Methods**). Using T2T-CN1 as the reference, APGp1 MC graph yielded 149,808 SVs across 71,434 sites when excluding centromeric and telomeric regions (**Supplementary Fig. 34a**). The pangenome graph identified over 79,000 additional SVs compared to assembly-based methods and more than 86,000 SVs beyond what PacBio HiFi read mapping could detect (**Fig. 6a**; **Supplementary Fig. 34b**). In the ‘easy’ regions, 91.5% (22,222) of our graph-based SVs were independently validated by at least one reference-based method, demonstrating the reliability and sensitivity of our approach.

**Figure 6.**
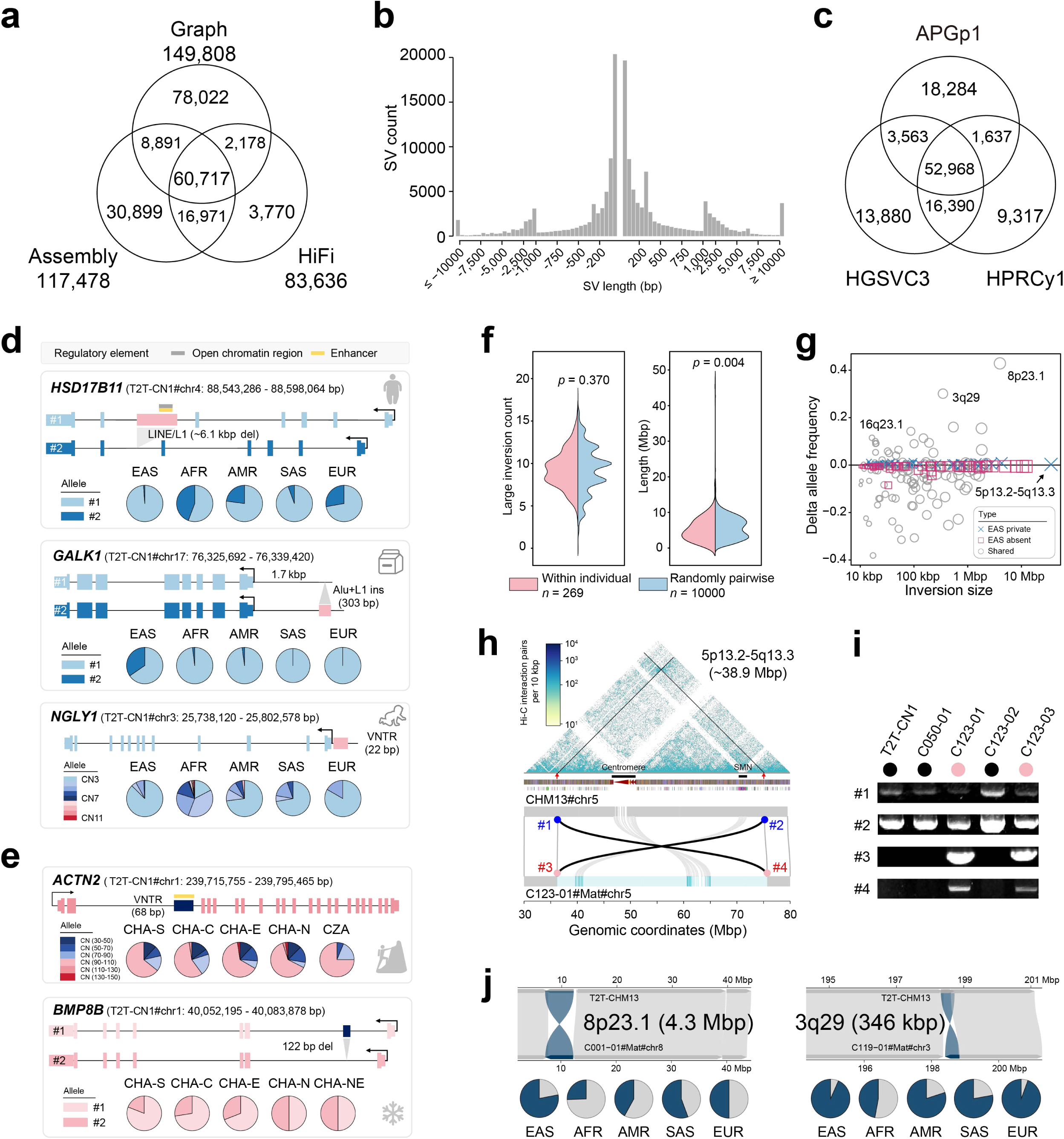
Genetic diversity and divergence of structural variations in human genomes. **a**, Shared SV alleles among pangenome-graph, assembly, and PacBio HiFi read mapping-based SV call sets. **b**, Length distribution of SVs identified from the pangenome graph. **c**, Overlap of SV loci across three human pangenome panels (APGp1, HPRCy1 and HGSVC3). **d**, SV allele frequencies in five super populations for genes *HSD17B11*, *GALK1* and *NGLY1*. **e**, SV allele frequencies among EAS populations for genes *ACTN2* and *BMP8B*. **f**, Inversion counts and sizes between pairwise haplotypes: random sampling *versus* within individuals. **g**, Frequency differences and sizes of large inversions (≥10 kbp), highlighting population stratification between EAS and non-EAS genomes (Delta allele frequency). **h**, Visualization of genomic synteny between the C123-01#Mat haplotype assembly and T2T-CHM13 around a 38.9-Mbp ultra-large inversion at 5p13.2-5q13.3. A Hi-C interaction heatmap aligning against T2T-CHM13 is overlaid to show the inversion signal. The centromere and SMN locus are specially indicated by black bars. Breakpoints of reference and inversion alleles are colored in blue and red, respectively. **i**, PCR products at 5p13.2-5q13.3 inversion breakpoints, obtained from the genomic DNA of T2T-CN1 (ref/ref), C050-01 (ref/ref) and C123 family (C123-01, child, ref/alt; C123-02, father, ref/ref; C123-03, mother, ref/alt). Two reference (Primer 1 and 2) and two inversion primers (Primer 3 and 4) are designed crossing the breakpoints (blue and red, respectively), with expected sizes of about 2 kbp. **j**, Synteny plots and frequency distribution in five super populations for inversions 8p23.1 and 3q29.

Size distribution of SV followed an expected exponential pattern (Liao *et al*., 2023; Sudmant *et al*., 2015), with distinct peaks at ∼300 bp and ∼6.5 kbp, corresponding to Alu and LINE/L1 elements, respectively (**Fig. 6b**). In total, these variations encompassed approximately 208 Mbp of euchromatic autosomal non-reference sequences. VNTRs were the most abundant SV class, while segmental duplications (SDs) dominated among large SVs (≥ 1 kbp; **Supplementary Fig. 35**). Deletions and insertions were evenly distributed across most repeat classes, a slight bias towards insertions was observed specifically in VNTRs, Alu elements and simple repeats (**Supplementary Fig. 35**). Consistent with previous studies (Gong *et al*., 2025; O’Donnell *et al*., 2023; Audano *et al*., 2019; Collins *et al*., 2020), we found that SVs were predominantly enriched in proximal telomeric and peri/centromeric regions (**Supplementary Fig. 36a**). Intriguingly, ‘easy’ regions harbored only 14.8% of SVs, despite these regions accounting for about three-quarters of the whole genome. Furthermore, we identified a negative relationship between SV frequency and recombination rates in the subtelomeric SV-rich regions, which contrasted with the positive correlation observed on chromosomal arms. These opposing patterns suggested a complex, spatially dynamic interplay between SV formation and homologous recombination in the human genome (**Supplementary Fig. 36b**).

When expanding our SV analysis by complementing T2T-CHM13 and assemblies from HPRCy1 and HGSVC3, we yielded 116,097 SV loci containing over 483,000 SVs relative to T2T-CN1. A total of 18,284 SV sites were unique to the APGp1 assemblies, though a large proportion (72.5%) were singleton (**Fig. 6c**). Additionally, APGp1 contributed 134,290 new SV alleles across 13,836 known SV sites, shared with HPRCy1 and HGSVC3 assemblies, expanding the allelic spectrum of previously characterized SVs. These SVs overlapped within the gene bodies and 5-kbp flanking regions, potentially affecting 5,020 protein-coding genes. Functional enrichment analysis revealed significant associations with nervous system development (GO:0007399, BH-corrected *p* = 4.62×10^-13^) and calcium signaling pathway (KEGG: hsa04020, BH-corrected *p* = 3.33×10^-4^; **Supplementary Table 14**). Significant enrichment was also observed with obesity and diabetes, as demonstrated by the NHGRI-EBI GWAS catalog (BH-corrected *p* = 1.46×10[^8^) and the Online Mendelian Inheritance in Man (OMIM) database (OMIM:125853, BH-corrected *p* = 0.0363; **Supplementary Table 15**). These functional categories are universally hyper-variable for SVs across human populations. The substantial number of novel alleles discovered here highlights the underestimation of allelic diversity in existing pangenome resources, underscoring the need for larger-scale pangenomic efforts.

We identified 2,935 SVs with high differentiation (Hudson *Fst* > 0.288, top 5%) between EAS and non-EAS populations. Among these, we discovered several functionally essential variants (**Supplementary Table 16**), including a ∼6-kbp retrotransposon long interspersed element-1 (LINE-1) insertion within the fifth intron of gene *HSD17B11* that was nearly fixed in EAS (98.9%) but polymorphic in other superpopulations (64.4%; **Fig. 6d**). This gene encodes short-chain alcohol dehydrogenases, associated with the metabolism of secondary alcohols and ketones (Brereton *et al*., 2001). Notably, this insertion was absent in great apes and Neanderthal genomes, suggesting a recent origin in modern humans and potential selection in EAS (**Supplementary Fig. 37**). Another example involves a 303-bp duplication in the 1.7-kbp upstream of *GALK1* (17q25.1), encoding galactokinase, a key enzyme in galactose metabolism (**Fig. 6d**). Deficiency of this enzyme causes congenital cataracts during infancy and presenile cataracts in adults (Rubio-Gozalbo *et al*., 2021). This duplicate consists of a SINE/Alu and a LINE-1 fragment, and showed the highest frequency in the EAS population (34.8%). Additionally, a 22-bp repeat VNTR upstream of *NGLY1* (3p24.2) showed distinct copy number distributions among superpopulations (**Fig. 6d**). Specifically, the 3-copy allele represents the most frequent type in all superpopulations (e.g., 87.4% for EAS) except AFR (17.3%). *NGLY1* deficiency causes congenital disorder of N-linked deglycosylation, characterized by developmental delay and intellectual disability during early childhood (Lam *et al*., 2017). Given its positional proximity to the promoter, the VNTR’s diverse copy number variation may act as a *cis*-eQTL to modulate the expression of *NGLY1*, potentially contributing to normal phenotypic diversity.

Within the EAS superpopulation, we identified 424 SVs displaying frequency differences between Han and Tibetan individuals, encompassing 170 protein-coding genes (**Supplementary Table 17**; **Supplementary Fig. 38a**). For instance, we observed a 104-bp intronic insertion within gene *ATP13A3*, with a much higher prevalence in Tibetans (75.0%). This gene is involved in cation transport and cellular ion balance, and has been identified under strong selection in Tibetans, potentially contributing to optimized cardiopulmonary function (Zheng *et al*., 2023). Furthermore the gene *ACTN2*, encoding alpha-actinin-2, a crucial cytoskeletal protein in cardiac and skeletal muscle cells, had higher copy numbers of a 68-bp repeat in its enhancer region among Tibetans (∼96.4 copies) compared to low-altitude Han populations (89.5; *P* = 0.0028, Wilcoxon rank-sum test; **Fig. 6e**). *ACTN2* displayed strong selection signals in Tibetan genomes, particularly in its upstream regulatory region (**Supplementary Fig. 39a**). The *ACTN2* enhancer is associated with heart failure in a GWAS (Arvanitis *et al*., 2020) and its expression levels may regulate heart rates (D’Antonio *et al*., 2023). Globally, the 68-bp VNTR exhibits extensive polymorphism across superpopulations. AFR displays the highest allelic diversity, whereas non-AFR superpopulations show divergent frequency spectrum, likely tracing back to out-of-Africa bottlenecks and genetic drift (**Supplementary Fig. 39b**). Similarly, despite overall weak genetic differentiation between North Han and South Han populations, we identified 33 highly stratified SVs by comparing AFs (**Supplementary Fig. 38b**; **Supplementary Table 17**). One example is a 121-bp deletion in *BMP8B* (**Fig. 6e**), a gene crucial for thermogenic function of brown adipose tissue and induces browning in white adipose tissue (Whittle *et al*., 2012), whose deletion frequency was higher in North Han than that in South Han (*P* = 0.001; Fisher’s exact test). Notably, EAS genomes exhibited elevated deletion frequency compared to other superpopulations (*P* = 3.49×10^-6^; Fisher’s exact test; **Supplementary Fig. 39c**).

### Large inversions

Genomic inversions represent a crucial yet challenging class of structural variations to detect, even with modern pangenome graph approaches. We characterized large genomic inversions (≥10 kbp) across the 536 assemblies from APGp1, HGSVC3 and HPRCy1 using three reference-based approaches (PAV, SVIM-asm and LGvar; **Supplementary Fig. 40**; **Methods**). A total of 159 balanced inversions were detected on the autosomes and chromosome X, excluding peri/centromeric and telomeric regions, with pronounced distribution on chromosomes 1, 7, 16, and X (**Supplementary Fig. 41**; **Supplementary Table 18**). Notably, large inversion is absent on chromosome 18 among all surveyed global assemblies of healthy individuals, suggesting a potentially constrained structural landscape. This scarcity may be attributed to purifying selection against inversion polymorphism, or to a paucity of local structural motifs (e.g., SDs or inverted repeats), that typically facilitate inversion formation. The genomic positions of inversions showed significant association with SDs at the breakpoints (*P* = 0.0001, permutation test), highlighting the mechanistic basis of their formation.The genes flanking these inversions showed significant enrichment in biological pathways related to immunoglobulin mediated immune response (GO:0016064), detection of chemical stimulus involved in sensory perception of smell (GO:0050911), and DNA cytosine deamination (GO:0070383; **Supplementary Fig. 42**).

Our population-level analysis revealed that each haploid genome typically carries approximately nine inversions relative to other genomes, spanning about 6.1 Mbp (**Fig. 6f**). The distribution pattern of these inversions followed an exponential relationship between size and frequency, with larger inversions exhibiting lower population AFs (**Fig. 6g**), suggesting selective pressure against large genomic rearrangements. However, it is notable that 19 inversions exceeded 1 Mbp in length, including one exceptionally large pericentric inversion in a single assembly from APGp1 (C123-01#Mat). This remarkable genomic rearrangement spans approximately 39 Mbp, extending from 5p13.2 to 5q13.3 (36.3 to 75.2 Mbp in T2T-CHM13) and traversing the centromere region (**Fig. 6h**). The authenticity of this exceptional inversion was rigorously validated through multiple independent approaches, including Hi-C interaction mapping, long-read alignments, and PCR verification (**Figs. 6h** and **6i**; **Supplementary Fig. 43**; **Supplementary Table 19**). Unlike typical pericentric inversions among healthy individuals, which encompass limited genes (e.g., 9p11-9q12/13; Hsu *et al*., 1987), this inversion encompasses more than 150 protein-coding genes, representing a scale of genomic rearrangement rarely observed in viable human genomics. Of critical importance, the right-side breakpoint intersects the *HEXB*, (**Supplementary Fig. 44**), which encodes an enzyme crucial for preventing ganglioside accumulation in neurons. Disruption of this gene has been previously linked to neurodegenerative disorders (Yoshizawa *et al*., 2002). This inversion likely originated through the interaction of inverted SINE-VNTR-Alu (SVA) elements at the breakpoints (**Supplementary Figs. 44 and 45**). While we confirmed the maternal inheritance of this inversion through both NGS mapping and PCR verification, ruling out a *de novo* mutation event (**Fig. 6i**; **Supplementary Figs. 43 and 44**), its presence in apparently healthy heterozygous carriers raises intriguing questions about genetic resilience and compensatory mechanisms. Although current clinical assessment of the carriers shows no obvious disorders through questioning and routine physical examination, the potential long-term impacts of such a massive rearrangement on human health warrant careful monitoring and further investigation.

We identified 18.2% (*n* = 29) of all inversions were exclusively observed in EAS assemblies, with the majority (*n* = 27) are rare variants showing AFs <0.01 (**Supplementary Table 18**). We verified their authenticity by long-read alignments and Bionano optical mapping (**Supplementary Figs. 46 and 47**). Among them, a 14.5-kbp inversion at 17q21.2 was only found in five EAS assemblies (*n* = 5), and this AF stratification among superpopulations was confirmed by its record in gnomeAD (**Supplementary Table 20**). Moreover, 58 inversions displayed stratification between EAS and non-EAS populations (*P* < 0.05, Fisher’s exact test), of which 15 exhibited elevated frequencies in EAS, including several occurring in genomic instability regions associated with neuronal development delay risks (e.g., 8p23.1 and 16p11; Wang *et al*., 2023). The largest among them, a 4.6-Mbp inversion at 8p23.1, showed much higher prevalence in EAS genomes (Yang *et al*., 2023; Porubsky *et al*., 2022; Salm *et al*., 2012) (**Fig. 6j**). Additionally, we found a ∼19-kbp inversion at 16q23.1 (from 81.251 to 81.271 Mbp in T2T-CHM13) nearly fixed in EAS (100%) and AFR (97%) genomes but absent in nearly one third of SAS and EUR assemblies (**Supplementary Table 18**). This inversion is located within the *CTRB1*-*CTRB2* locus and has been associated with chronic pancreatitis risk in EUR populations but no effect in Chinese population due to allele fixation (Pang *et al*., 2013; Rosendahl *et al*., 2018; Tang *et al*., 2018). Furthermore, a ∼346-kbp inversion at 3q29 exhibited higher frequencies in EAS (93.4%) and EUR (94.4%), compared to AFR (47.1%; **Fig. 6j**). This SV appeared to alter local chromatin architecture and modified loop interactions surrounding *MUC4*, a gene involved in epithelial renewal and differentiation (Chaturvedi *et al*., 2007; **Supplementary Fig. 48**).

### Structural diversity of complex loci

Our MG pangenome graph reveals several genomic islands with structural hypervariability along the chromosomes, representing SV hotspots in human genomes (**Supplementary Fig. 49**). These high-complexity regions primarily emerge from recurrent segmental duplications and rearrangements. Among these, several well-characterized complex loci stand out, including the immune-related β-defensin cluster on chromosome 8, *CLEC* cluster on chromosome 12, and neurodevelopment-related regions such as *TBC1D3* cluster on chromosome 17 and *NPIP* cluster on chromosome 16. Some high-complexity spots arose from tandem repeat variants (VNTRs or STRs), including the macrosatellite repeat DXZ4 cluster within gene *DANT2* on chromosome X and the hAT-Charlie repeat cluster located between *COPS7B* and *MIR1471*. To illuminate the profound structural diversity hidden within these complex regions, we conducted analyses of two critical genomic loci, MHC and SMN regions.

### MHC region

The MHC region represents one of the most intensively studied human genomic regions, with over seven decades of research driven by its critical role in immunity and disease. Despite this extensive investigation, the structural variations within this region have remained largely enigmatic, obscured by its extraordinary genetic diversity and sequence complexity across human populations. The MHC region shows a prominent peak of structural variations (**Fig. 7a**). The completeness of our dataset is remarkable with all APGp1 and HGSVC3 assemblies achieving gapless coverage of the MHC region, while three assemblies from HPRCy1 are incomplete in this region (**Supplementary Table 21**). Through multiple annotation approaches, we found that each haplotype harbors an average of 37.0 HLA (Human Leukocyte Antigen) genes, ranging from 31 to 40, alongside an average of 172.4 non-HLA genes/pseudogenes, ranging from 167 to 182 (**Fig. 7b**; **Supplementary Table 21**). These numbers substantially exceed those previously reported in HGSVC3 (Logsdon *et al*., 2025). The phylogenetic tree constructed for all annotated multiple-copy HLA genes exhibited well-defined monophyletic clustering to each HLA gene category, strongly validating the accuracy and reliability of our HLA gene annotation (**Supplementary Fig. 50**). Intriguingly, it seems that EUR assemblies contained more HLA genes compared to other superpopulations (*P* = 0.018, two-sided Wilcoxon rank-sum test), which requires larger sample sizes to validate (**Fig. 7b**).

**Figure 7.**
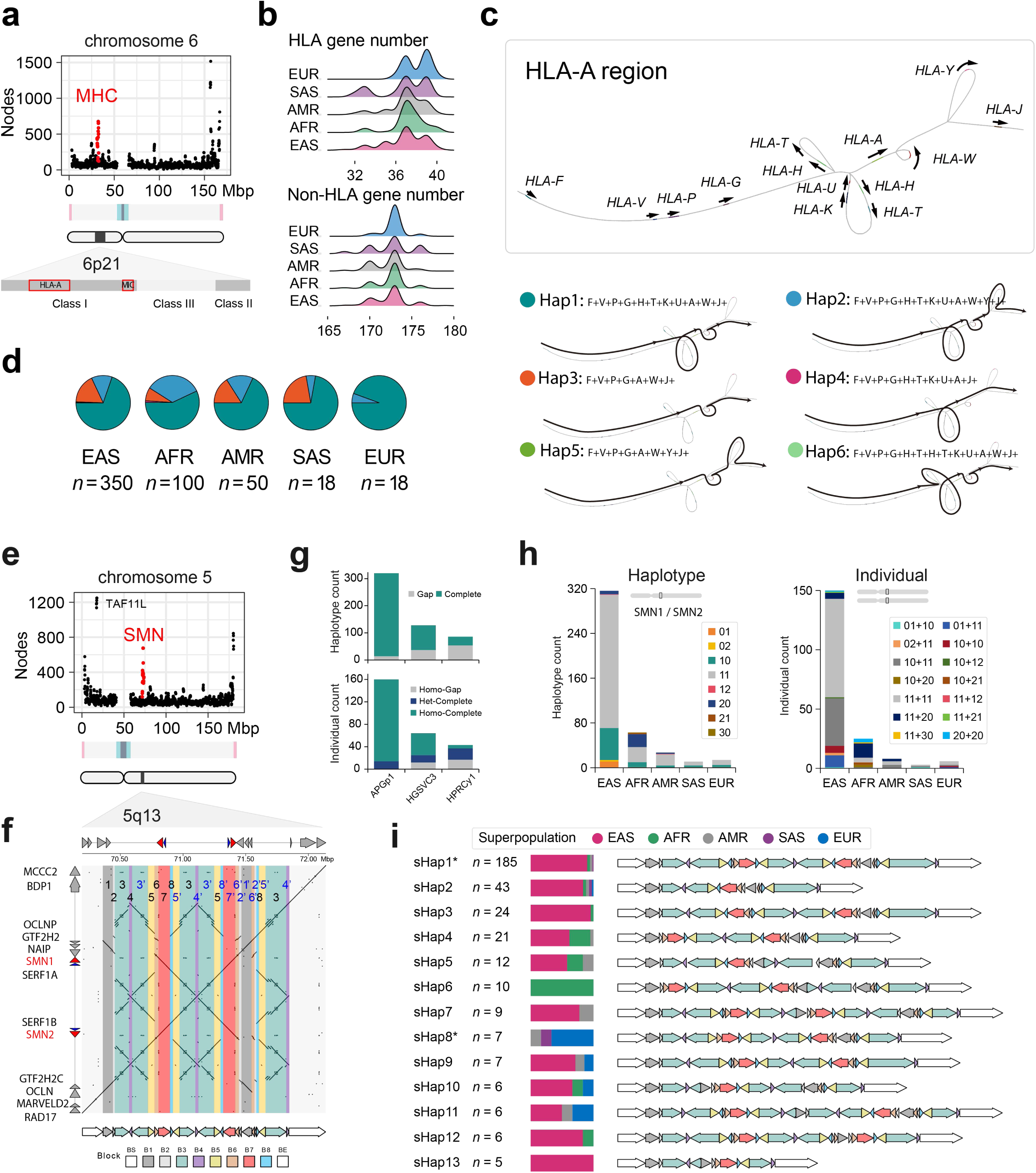
Structural diversity of MHC and SMN loci in the human genome. **a**, Complexity for chromosome 6 in the Minigraph (MG) pangenome graph (APGp1+HPRCy1+HGSVC3), as measured by node counts in sliding windows. The location of the MHC locus is highlighted in red. **b**, Counts of HLA and non-HLA genes across five super populations. **c**, Structural haplotypes of the HLA-A region from the Minigraph-Cactus (MC) graph. Different structural haplotypes take different graph paths, visually represented by bold lines with arrows. **d**, Haplotype frequency across five super populations. **e**, Graph complexity for human chromosome 5 in the global MG graph. The genomic position of the SMN locus is marked in red. **f**, Gene annotation and sequence dot plot within the SMN region of T2T-CHM13 genome. Eight principle blocks (from B1 to B8) are defined to dissect the structures of SMN sequences. The prime symbol represents a reverse strand, compared to the respective defined block. **g**, Haplotype and individual counts with gapless and incomplete assemblies over the SMN region. **h**, Copy number and combination of *SMN1* and *SMN2* genes at the haploid (haplotype) and diploid (individual) levels across five super populations. To be specific, “01” denotes zero copies of *SMN1* and one copy of *SMN2*, and “11+20” represents that one allele contains one copy of *SMN1* and one *SMN2*, while the other has two copies of *SMN1* and no *SMN2*. **i**, Representation of 13 common structural haplotypes (sHaps) in the SMN region, and their respective proportions in the five super populations. Common haplotypes are defined as those with an allele frequency greater than 1% (i.e., *n* ≥5 here).

Previous studies have linked *MICA* to autoimmune diseases such as rheumatoid arthritis and coeliac disease, and its deletion may increase the pathogenicity risk (Choy & Phipps, 2010). Within the MIC region of MHC Class I cluster, we identified two distinct structural haplotypes, with one represented by the three reference assemblies, and the other containing a ∼94-kbp deletion. This deletion, present in ten haplotype assemblies from EAS, AFR and AMR superpopulations, results in the complete loss of *MICA*, *HLA-X* and several other genes/pseudogenes (**Supplementary Fig. 51**).

Within the HLA-A region spanning from *HLA-F* to *HLA-J*, we discovered six structural haplotypes from the MC graph, comprising three common (Hap1 to Hap3) and three singleton haplotypes (Hap4 to Hap6, ordered by haplotype frequency; **Fig. 7c**). Among them, Hap5 (C001-CHA-E01-01#Pat) and Hap6 (C002-CHA-E02-01#Pat) represent novel structural architectures that were unreported previously (Liao *et al*., 2023), both being exclusively captured in the APGp1 assemblies (**Supplementary Fig. 52**; **Supplementary Table 21**). Compared to the predominant haplotype Hap1 (represented by GRCh38 and T2T-CHM13), Hap5 displayed a deletion of *HLA-H*, *HLA-T*, *HLA-K*, and *HLA-U*, and an insertion of *HLA-Y*. Structural comparison and phylogenetic analysis revealed that Hap5 was derived from a recent recombination between Hap2 and Hap3 (**Supplementary Fig. 52**). Hap6 featured a ∼50-kbp recent duplication encompassing the *HLA-H* and *HLA-T* loci (**Supplementary Fig. 53**). Furthermore, Hap2, carrying a *HLA-Y* insertion represented by T2T-CN1 reference genome, is present across all populations, while it displays much elevated frequency in AFR (34.7%) relative to other superpopulations (12.1%; **Fig. 7d**).

### SMN region

The *SMN1* and *SMN2* genes are essential determinants for human spinal muscular atrophy (SMA), an autosomal recessive disorder characterized by progressive degeneration of lower motor neurons. Most SMA cases are caused by homozygous absence of *SMN1* (0-copy), due to deletion or *SMN1*-to-*SMN2* gene conversion (Keinath *et al*., 2021). These two SMN genes reside within an extremely complex region at 5q13.2 (**Fig. 7e**), which harbors extensive multi-layered inverted repeats (**Fig. 7f**). We successfully resolved the complete sequences of SMN loci in 95.6% of the 320 APGp1 assemblies, exceeding the completeness achieved in HPRCy1 (37.2%) and HGSVC3 (71.1%; **Fig. 7g**). Moreover, both maternal and paternal copies of SMN loci were completely resolved in 146 individuals of APGp1 (91.3%; **Fig. 7g**), providing an opportunity to investigate the diploid architecture of this critical region. The 434 fully resolved SMN loci across global populations exhibited remarkable length variations, demonstrating remarkably evolutionary dynamics that correlate with the CNV of SMN genes (**Supplementary Fig. 54**). Phylogenetic analysis of 937 SMN genes across all assemblies showed clear separation between *SMN1* and *SMN2*, with multiple fixed divergent sites, including the causative variant C840T responsible for functional differences (**Supplementary Fig. 55**).

The majority of assemblies (70.0%, 304/434) contained the canonical configuration of one copy each of *SMN1* and *SMN2* (**Fig. 7h**). However, AFR assemblies showed notably higher *SMN1* CNs, with 41.3% (26 out of 63 complete SMN sequences) containing two (*n* = 25) or three copies (*n* = 1), contrasting sharply with EAS where multiple *SMN1* copies appear in only 1.9% of cases (6/316; **Supplementary Table 22**). Phylogeny and sequence alignments suggested four haplotypes for *SMN1* sequences and the majority of AFR multi-copy *SMN1* alleles were duplicates of an AFR-enriched *SMN1* haplotype (20/26), whereas the two-copy *SMN1* alleles in non-AFR assemblies have no such duplicate (**Supplementary Fig. 55**). This finding highlights how similar genotypes can emerge through distinct evolutionary trajectories leading to the convergent CNVs in the two populations.

In AFR, 52.4% of assemblies (33 of 63) exhibited *SMN2* loss, much more frequent than other superpopulations (EAS: 20.0%, 63/316; AMR: 22.2%, 6/27). Despite the overall constraints on *SMN1*, 17 gap-free assemblies (3.9%, of 434) deleted this essential gene, similar to the proportion observed in the recent HGSVC3 analysis (Logsdon *et al*., 2025). Among the 192 individuals with both parental SMN loci fully resolved, all retained at least one functional *SMN1* copy (**Fig. 7h**). Non-AFR individuals predominantly carried one copy of *SMN1* in both maternal and paternal haplotype (88.0%, 132/150), while AFR individuals commonly displayed a combination of a two-*SMN1*-copy and a one-*SMN1*-copy haplotype (64.0%, 16/25; **Fig. 7h**).

We next sought to characterize the structural diversity of SMN loci in human genomes from sequences, which has not been fully characterized due to extreme dynamics of nested multi-layered palindromes (**Fig. 7f**). Even in the pangenome MC graph, this region shows chaotic connections, indicating the failure in building accurate local graphs. To represent and quantify the large-scale structures of SMN loci, we defined eight principal genomic blocks as the minimizer repeat units, where *SMN* and *SERF1* genes were embedded within Block 7 (B7), based on comprehensive pairwise sequence alignments across SMN loci (**Fig. 7f**). We found that only one single copy of each block and no palindromic structures in the non-human primate genomes, suggesting human-specific structural rearrangements (**Supplementary Fig. 56**).

These eight blocks were used to decompose SMN sequences and assign structural haplotypes (sHaps). Individual blocks displayed limited structural variants, primarily including large deletions in B1 and in B6, alongside hyper-variable nested VNTRs in B3 (**Supplementary Fig. 57**). Averagely 98.4% of SMN sequences per assembly were covered, yielding 74 block-based structural types across global genomes, including 13 common (AF > 0.01), 17 rare (AF ≤ 0.01) and 44 singleton sHaps (**Supplementary Figs. 58-60**). Notably, the most frequent configuration sHap1, represented by T2T-CN1, was shared across 185 assemblies, while the T2T-CHM13 and GRCh38 represents two rare configurations classified into sHap8 and sHapR4, capturing only seven and four assemblies, respectively (**Fig. 7i**; **Supplementary Table 22**). Several haplotypes exhibited population stratification (**Fig. 7i**; **Supplementary Fig. 59**). The most common haplotype (sHap1) reaches a frequency of 52.5% in EAS (166/366) but dropped to 15.7% in non-EAS assemblies (18/115). Conversely, some haplotypes show potentially exclusive population restriction, with sHap6 being endemic to AFR assemblies, and sHap8 being absent from both EAS and AFR while reaching 29.6% frequency in EUR. We note that given the undersampling and technical challenges associated with resolving complete SMN sequences, these population frequency dynamics should be treated as preliminary estimates awaiting validation in larger pangenome cohorts.

## Discussion

Recent advancements in long-read sequencing technologies have unlocked unprecedented access to complex and structurally divergent genomic sequences in the human genome. At this critical juncture, conducting regional long-read human genome studies offers a unique opportunity to not only address the under-representation of certain populations in genetic research but also simultaneously investigate complex genomic regions, thereby advancing our understanding of human evolution and health across diverse ethnic populations. Here, we present 320 nearly T2T and phased human genome assemblies from 160 EAS individuals in APGp1 with superior quality, compared to previously reported human pangenome resources (Liao *et al*., 2023; Gao *et al*., 2023; Littlefield *et al*., 2024; Logsdon *et al*., 2025; Nassir *et al*., 2025). By comprehensively annotating repeatome and gene elements across these assemblies, we characterized fine-scale variation in peri/centromeres, rDNA arrays, and medically relevant complex regions (e.g., MHC and SMN loci), revealing genetic diversity that has been largely inaccessible to previous short-read-based studies.

Important lessons emerge from our study that can inform future genomic research directions. First, achieving sequence diversity saturation requires continued recruitment from under-represented populations. Our analysis demonstrates that with >500 global assemblies, novel sequences continue to be discovered, emphasizing the need for broader sampling. Importantly, severe sampling bias hinders the inference of evolutionary and biomedical implications of complex haplotypes and alleles showing population stratification in frequency. Although global efforts have been to made to build global or regional pangenomes, a unified comparison and integration framework among pangenome graphs are also urgently needed. Second, our studies and others show that while long-read sequencing enables highly continuous assemblies, persistent challenges remain in resolving complex regions such as rDNA arrays, telomeres, peri/centromeres, due to their highly repetitive and intrinsic sequence patterns (Nurk *et al*., 2022; Rhie *et al*., 2023). These technical limitations underscore the urgent need for specialized quality control frameworks and analytical approaches are needed to define standards for T2T-era assemblies. Third, ensuring the representativeness and interpretability of complex regions remains unaddressed. Current tools struggle with the MC graph construction for inversion-rich loci like SMN, requiring manual curation and context-specific strategies. The minimizer-decomposition strategy represent one of solutions, such as PGGB and PGR-TK, as exemplified by the of amylase locus (Chin *et al*., 2023; Garrison *et al*., 2024; Bolognini *et al*., 2024). Fourthly, evolutionary framework and statistical models for complex variants are lacking, which hinders effective inference of selection. Whether SNVs are sufficient for population genetics analysis is an open question. Finally, while pangenome graph-based genotyping tools enhance SV detection for large cohorts with NGS data (Ebler *et al*., 2022; Sirén *et al*., 2021), they can not resolve the novel variants absent from the pangenome reference graph, highlighting the need for iterative graph augmentation approaches. As the field transitions toward large-scale long-read sequencing, emerging graph-based mapping methods (e.g., SVarp and SVPG) that align new long reads directly to existing pangenomes represent a vital future direction for comprehensive SV analysis (Schloissnig *et al*., 2025; Jiang *et al*., 2025). Nevertheless, to fully capture hidden genomic diversity, these advanced mapping paradigms must be coupled with highlighting the need for iterative graph augmentation approaches.

As a flagship work of the APG project, phase 1 provides a genome-wide full-spectrum sequence landscape and a genetic foundation for understanding human adaptation and disease susceptibility in EAS superpopulation, even as under-represented regions (e.g., Japan and Mongolia) require deeper sampling in the near future. Details of the genetic diversity and evolution of human centromeres, sex chromosomes, and other repeatomes are explored in accompanying companion papers (He et al., 2025; Fu *et al*., 2026; Liu *et al*., 2026, under review; Sun *et al*., 2026, under review; Suo *et al*., 2026, under review; Yang *et al*., 2026, under review). The APG team is actively expanding recruitment for phase 2, aiming to generate 600 complete, phased T2T assemblies spanning diverse, multi-regional Asian populations. This expansion will directly address the geographical and ethnic gaps in 1KGP, the primary sampling source of HPRC and HGSVC, to some extent. Through global collaborative efforts, as championed by the Human Pangenome Project (https://humanpangenomeproject.org/), we anticipate the imminent realization of a new “1000 Genomes Project” comprising thousands of entirely near-T2T *de novo* assemblies from global ancestries. This comprehensive human pangenome will unlock unprecedented resolution into human genomic architecture, benefiting basic research in human evolution and disease biology while accelerating precision medicine applications.

## Acknowledgments

We thank all the participants in this project. We sincerely thank Tibetan Fukang Hospital for their valuable assistance with sample collection in this study. Computational infrastructure and support are provided in part by the Super-computing platform of Innovation Center of Yangtze River Delta Zhejiang University, High Performance Computing cluster of Zhejiang lab, the Super-computing Center of Hangzhou City University, and HPC of Center for Bioinformatics and Big Data Technology, Zhejiang University. We also acknowledge the Information Technology Center of Zhejiang University and China Mobile Zhejiang Co., Ltd (Hangzhou Branch) for providing the computation resource. This work was funded, in part, by Fundamental and Interdisciplinary Disciplines Breakthrough Plan of the Ministry of Education of China (No. JYB2025XDXM508), the funding from International Institutes of Medicine of Zhejiang University (KY2022-098), National Key R&D Program of China (2024YFA1802500), Basic Research Center Program (32388102), the New Cornerstone Science Foundation through the XPLORER PRIZE and the K.C.Wong Education Foundation to G.Z., National Key R&D Program of China (2025YFC3410300) to D.W., C.Y., Y.M., Xiangyu Yang, XiaofeiYang, Yaoxi He and B.S., National Natural Science Foundation of China (32370658 and W2623001), the Scientific Research Innovation Capability Support Project for Young Faculty (grant no. SRICSPYF-ZY2025101), “Shuguang Program” supported by Shanghai Education Development Foundation and Shanghai Municipal Education Commission (25SG17), Shanghai Jiao Tong University 2030 Initiative (WH510363003/016), Natural Science Foundation of Chongqing, China (CSTB2024NSCQ-JQX0004), the Computational Biology Program of Science and Technology Commission of Shanghai Municipality (24JS2840300), and the New Cornerstone Science Foundation through the XPLORER PRIZE to Y.M., National Key R&D Program of China (2022YFC3400300), the National Natural Science Foundation of China (32422019, 62172325), the Natural Science Foundation of Shaanxi Province (2024JC-JCQN-28), the Fundamental Research Funds for the Central Universities (xzy012024088), and the Scientific Research Program of Shaanxi Provincial Department of Education (23JK0290) to Xiaofei Yang.

## Author contributions

Project conception and design: G.Z., T.Z., S.Ying, D.Z. and Y.M.; Project supervision: G.Z., T.Z., S. Ying and H.H.; Ethics management: D. Yu, D.W., L.Y., Y. Liu, Yunqiu He and J.C.; Sample collection: D. Yu, D.W., L.Y., D.S., Y. Lu, Xiangyu Yang,Yiqing Yang, Jie Yu, J.Q., Xiangyu Yang, D. Yang, Y.J., Feng Zhou, W.Z., F.W., Yihua Yang, Z. Zeng, Z.F., Y.C., Yafeng Li, Yaheng Li, S.Z., A. Long, Z.W., Q.L., R.Z., G.D., Q.W., Y.T., Jingjing Yu, H.L., K.L., Yu Zhang, X. Yan, Dawa, Yanfei Zhang, A.B., G.C., S.H.Q., X.L., X.B., J. Liu, J. Li, S.W., J. Yu, Qi Zhou (Yiwu), J.J., W.X., Aixia Liu, J.M., Y. Zhao, Yunqiu He, F.L., J.Yang, A.A.-B., Y.Z., A. Lin, Yaoxi He, J. Huang, K.W., J.X. and D.Z.; Sequencing: W.D., D. Yu, Yiqing Yang, D.S., Y. Lu, Y. Zhao and J.M.; Ancestry analysis: M.S. and D.W.; Genome assembly, gap-filling and polishing: D.W., C.Y., L.N. and Q.C.; Assembly quality control and evaluation: D.W., C.Y., Q.C., J. Han, Y. Sun and Q.N.; Repeat and gene annotation: Q.C., Y. Sun and Q.N.; Centromere analysis: Y. Sun, K.F. and S.Ye.; rDNA element analysis: L.N. and D.W. CNV analysis: D.W. and Q.C.; Pangenome graph construction and comparison: M.S. and D.W.; Analysis of pLoF variants: Anguo Liu and D.W.; SV identification and comparison: C.Y., Q.C., T.Y., L.Q. and L. Fu.; Inversion: Feifei Zhou, D.W., D.Y., Q.N. and Y.M.; Complex loci analysis (MHC and SMN): D.W., Q.C. and L.N.; Data coordination and management: D.W., D.Y., L.N. and L.Z. Computing resource support and database development: L. Fan, Z. Zhang, W.F., W.W., Y. Zhou, B.H. and C.L.; Principal investigators and laboratory organizers in APG consortium: G.Z., S.Ying, D.Z., Y.M., B.S., K.Y., Y. Shi, Qi Zhou (Hangzhou), C.X., Xiaofei Yang, C.Y., X.W., Yaoxi He, L.Y., D.W. and D. Yu.; Paper writing: D.W., C.Y., Y.M., G.Z., Q.C., M.S., Anguo Liu, D. Yu, Feifei Zhou, L.N., Y. Sun and J. Han.

## Methods

### Ethics declarations

This study was conducted in accordance with the Helsinki Declaration and approved by multiple ethics committees, including the Biomedical Research Involving Humans Ethics Committee in the Fourth Affiliated Hospital of Zhejiang University School of Medicine (Approval No. K2022126), the Human Research Ethics Committee of the Second Affiliated Hospital of Zhejiang University School of Medicine (Approval No. 2022-0699), Ethics Committee of People’s Hospital of Xinjiang Uygur Autonomous Region (Approval No. KY2023020305), Shanxi Provincial People’s Hospital Ethics Committee (Approval No. 2023-08), Medical Ethics Committee of the First Affiliated Hospital of Guangxi Medical University (Approval No. 2022-K037-01), Ethics Committee of Tibet Autonomous Region The Third People’s Hospital (Approval No. 202204), Ethics Committee of Huaihua City Maternal and Child Health Care Hospital (Approval No. 202204), Ethics Committee of Guizhou Medical University Affiliated Hospital (Approval No. 2023-022), Ethics Committee of the Second Affiliated Hospital, Heilongjiang University of Chinese Medicine (Approval No. 2022-K244), and The Ethics Committee for Human-Related Scientific Research, Kunming Institute of Zoology, Chinese Academy of Sciences (Approval No. KIZRKX-2025-002). All participants provided written informed consent after receiving a comprehensive explanation of the study’s objectives, procedures, potential risks, and benefits. For volunteers under 18 years, informed consent was obtained from their legal guardians, with additional assent acquired from the minors themselves. To safeguard participant privacy, all collected data were anonymized, and individual-level data was strictly limited to authorized personnel.

### Sampling

In APG phase 1, we recruited 160 East Asian individuals, including 152 participants from China and eight from South Korea. All participants provided fully informed consent and declared being in good health status. To capture representative genetic diversity across East Asia, volunteers were enrolled from geographically diverse regions through multiple collaborative sampling centers. Each individual was required to have consistent ethnic and regional ancestry across at least three generations (parents and grandparents), as verified by self-report. (Sub)population labels correspond to self-reported ethnic groups. For Han Chinese participants, regional subgroups were further classified based on geographic administrative divisions and declared ancestral origins. For each defined population, individuals were recruited from multiple local regions to reduce sampling bias. In addition, the kinship among sequenced individuals was calculated by using NGS genotyping data to confirm their genetic independence.

For each participant, approximately 20-30 mL of peripheral blood was collected in EDTA-coated tubes and gently inverted 10 times post-collection to ensure uniform anticoagulation. Samples were cryopreserved at -80°C immediately after sampling in a constant-temperature ultra-low freezer (Thermo Fisher Scientific) using cryogenic vials. Additionally, for trio-based haplotype phasing in genome assembly, 5-10 mL of peripheral blood was obtained from the parents of 142 participants, providing separate written informed consent for their participation. Blood samples with visible hemolysis or clotting were excluded, and replicate aliquots were prepared for each sample to minimize freeze-thaw cycles.

### DNA extraction and sequencing

#### PacBio HiFi sequencing

Peripheral blood samples were thawed at 37 °C, washed with pre-cooled buffer, and centrifuged to isolate cells. Cell pellets were lysed with buffers containing RNase A and Proteinase K, followed by phenol and chloroform–isoamyl alcohol extraction to remove proteins and other contaminants. DNA was precipitated with anhydrous ethanol, washed twice with 75% ethanol, and resuspended in Buffer EB. Mechanical shearing was minimized to preserve DNA integrity throughout the process. DNA quality and fragment size were evaluated by spectrophotometry, fluorometry, and pulsed-field gel electrophoresis to ensure suitability for PacBio HiFi sequencing (Sequel II and Revio platforms). PacBio HiFi circular consensus sequence (CCS) reads were generated from subreads by ccs (https://github.com/PacificBiosciences/ccs).

#### Ultra-long ONT sequencing

Genomic DNA was extracted from 5□mL whole blood using an SDS lysis buffer (1% SDS, 100□mM Tris-HCl pH8.0, 50□mM EDTA, 100□mM NaCl) at 65□°C for 60- 90□min. Proteins were removed with phenol:chloroform:isoamyl alcohol (25:24:1), and DNA was precipitated with isopropanol, washed with 75% ethanol, air-dried, and dissolved in EB buffer. DNA quality and yield were evaluated by spectrophotometry and gel electrophoresis. Libraries were prepared using the Oxford Nanopore SQK-LSK114 kit following a modified hexammine cobalt-based NEMO bead protocol. Briefly, high-molecular-weight DNA was precipitated onto sterilized glass beads, washed with PEGW buffer, and eluted in EB at 37□°C. The DNA library (75□μL DNA + 75□μL SQB) was incubated at room temperature for 30 min and loaded onto pre-primed flow cells. Sequencing was performed on the Oxford Nanopore PromethION platform, with each flow cell (R9.4.1 or R10.4.1) reloaded every 24□hours for up to three runs.

#### Hi-C sequencing

Frozen peripheral blood samples were thawed at 37 °C for 3 min, diluted with pre-cooled 1× PBS, and centrifuged at 2,500× g for 5 min at 4 °C to collect white cell pellets. Pellets were resuspended in PBS and cross-linked with 1% formaldehyde for 1 hour at room temperature on a vertical mixer, followed by quenching with 2.5 M glycine. Cross-linked cells were harvested by centrifugation at 4,000× g for 5 min at 4 °C, flash-frozen in liquid nitrogen and stored at –80 °C. Hi-C libraries were generated through proximity ligation and biotin-based enrichment, followed by fragmentation and pull-down of ligation junctions. Libraries were prepared using the MGIEasy Universal DNA Library Prep Set, enriched with Agilent SureSelect XT HS probes, and sequenced on the MGI DNB-SEQ platform using a 150-bp paired-end sequencing strategy.

#### PCR-free NGS sequencing

Genomic DNA was extracted from peripheral blood using an automated magnetic bead-based protocol (CWbio-CW2361S). Blood samples were mixed with Proteinase K, incubated, and then an isopropanol–magnetic bead mixture was added to capture DNA. Eluted DNA was stored at –20 °C until library preparation. For library construction, 50-1000 ng of high-quality genomic DNA was enzymatically fragmented with FS Buffer II and FS Enzyme Mix II at 30 °C, followed by enzyme inactivation at 65 °C. Fragmented DNA was size-selected using magnetic beads to enrich for 450-600 bp (for high input) or 600-750 bp (for low input) fragments. The DNA was end-repaired, A-tailed, and ligated with MGIEasy PF adapters. Ligated products were purified, denatured at 95 °C, and circularized at 37 °C, with non-circular DNA digested by exonuclease treatment. Final single-stranded circular (ssCir) libraries were quantified using Qubit ssDNA assay and assessed by Agilent Bioanalyzer. Libraries with yields ≥75 fmol were selected for sequencing on MGI platforms, including T7 and T20.

#### Bionano optical map

Five samples were selected for generating Bionano optical maps to validate the accuracy of EAS-specific large inversions. High-molecular-weight genomic DNA was isolated from 3 mL of frozen whole blood using the Nanobind CBB Big DNA Kit, following the Bionano Prep SP Frozen Human Blood DNA Isolation Protocol v2 (Bionano Genomics, https://bionano.com/wp-content/uploads/2023/01/30395-Bionano-Prep-SP-Frozen-Human-Blood-DNA-Isolation-Protocol-v2.pdf). DNA was labeled and counterstained via the Direct Label and Stain (DLS) protocol, then loaded onto a Saphyr chip and imaged on the Saphyr instrument (Bionano Genomics).

#### Ancestry inference

To characterize ancestry and genetic diversity in APGp1, we processed paired-end NGS short reads using fastp (v0.23.4) with the parameters ‘*-u 30 -q 20*’ (Chen *et al*., 2018) for quality trimming and adapter removal. Clean reads were uniformly down-sampled to 30× and aligned to T2T-CHM13 using *bwa mem* (Li, 2013). Small variants were called using GATK HaplotypeCaller (v4.1.8.1; Van der Auwera, *et al*., 2013), and merged with bcftools (v1.16; Danecek *et al*., 2021). For trios, identity-by-descent (IBD) was calculated to confirm parent-child relationship with plink (v1.90; Chang *et al*., 2015). Furthermore, we integrated APGp1 variants with 1KGP T2T-CHM13-referenced variant calls (https://github.com/marbl/CHM13), including five East Asian (EAS) populations: KHV (Kinh in Ho Chi Minh City, Vietnam), JPT (Japanese in Tokyo, Japan), CDX (Chinese Dai in Xishuangbanna, China), CHB (Han Chinese in Beijing, China), and CHS (Southern Han Chinese, China). Ancestry characterization was primarily conducted via principal component analysis (PCA) using plink (v1.90).

### Genome assembly

#### Draft assembly

PacBio HiFi reads were filtered using hifiadapterfilt with the parameter ‘*-l 44 -m 97*’ (Sim *et al*., 2022). Ultra-long ONT reads (>100 kbp) were selected for genome assembly to enhance contiguity. Genomic *k*-mer library per individual was constructed with yak (*k* = 31) (https://github.com/lh3/yak) and meryl (*k* = 30; Rhie *et al*., 2020). The paired-end Hi-C short reads were filtered using fastp. For trio samples (*n* = 142), we generated initial phased maternal and paternal assemblies (hifiasm-trio) for each offspring individual by integrating both PacBio HiFi and ultra-long ONT reads, using hifiasm (versions 0.19.2-r560 to 0.19.8-r603) in the trio-based phasing mode (**Supplementary Table 2**). To address gaps in the hifiasm-trio assemblies, we incorporated complementary assemblies generated by Verkko in trio mode (Verkko-trio) and by hifiasm in Hi-C mode (hifiasm-Hi-C) for each individual (**Supplementary Fig. 3**). The hifiasm-Hi-C phased assemblies were produced by leveraging the Hi-C information to guide phasing. These Hi-C phased contigs were subsequently evaluated and merged by parental *k*-mers, enabling re-phasing the contigs into biologically paternal and maternal haplotype assemblies. Verkko was employed to generate phased assemblies utilizing trio information (versions ranging from v1.0 to v1.4.1; Rautiainen *et al*., 2023). To avoid introducing new gaps, the scaffolded Verkko assemblies were deliberately split into contigs before subsequent gap-filling step. For individuals with no availability of parental NGS data (*n* = 18; **Supplementary Table 1**), Hi-C information was utilized to generate phased assemblies from PacBio HiFi and ultra-long ONT reads using hifiasm. The maternal and paternal haplotypes were temporarily assigned according to the presence of sex chromosomes X and Y.

#### Gap-filling by assemblies

Using hifiasm-trio phased assemblies as backbones, we iteratively integrated phased hifiasm-Hi-C and Verkko-trio assemblies to fill gaps using a custom Perl script gfasm.pl (https://github.com/Asian-Pan-Genome/APGp1/blob/main/Genome_assembly/assembly). Specifically, to fill the gaps in the reference backbone, query contigs were mapped against the reference contigs using minimap2 (v2.26-r1175; Li, 2018), retaining alignments with >20 kbp aligned length and mapping quality > 55. To avoid overfilling artifacts, we required that the flanking unaligned proportion be less than 0.05 of the total length of reference contig, and the aligned proportion of the reference contig aligned within the query contig exceed 0.80.

#### Gap-filling by ONT reads

Gaps were further filled using yagcloser (https://github.com/merlyescalona/yagcloser) with ONT long reads. ONT reads were firstly phased into paternal and maternal haplotypes by utilizing parental genomic information using the command ‘*canu -p asm -d binning genomeSize=3g useGrid=false maxThreads=${threads} - haplotypePat $pat_ngs -haplotypeMat $mat_ngs -nanopore-raw ${ont} - stopAfter=haplotype -corMhapOptions=”--threshold 0.8 --ordered-sketch-size 1000 --ordered-kmer-size 14” -correctedErrorRate=0.105*’ (Nurk *et al*., 2020). Subsequently, the phased ONT reads were mapped against corresponding draft haplotype assembly using minimap2 (v2.26-r1175), with ‘*--secondary=yes*’ to tag secondary alignments, then converted to sorted and indexed BAM format using samtools (v1.7; Li *et al*., 2009). Primary alignments with MAPQ > 20 were filtered to identify reliably spannable gaps. For each gap, a consensus sequence was generated from spanning ONT reads (v0.4 in **Supplementary Fig. 3**). To avoid over-filling and introducing large structural errors, alignments between assemblies and T2T-CHM13 and T2T-CN1 were visually inspected using LinkView (https://github.com/YangJianshun/LINKVIEW2). Potential large structural inversions and translocations were manually curated by examining ONT read alignments around the breakpoints and contigs were split at the breakpoints with no sufficient support of read alignments.

#### Assembly polishing

The assemblies underwent multiple polishing procedures to enhance base accuracy (**Supplementary Fig. 3**). For each phased assembly, phased ONT reads and all PacBio HiFi reads were aligned to the draft using minimap2. Potential structural variants (SVs) were identified with Sniffles (v2.3.2; Smolka *et al*., 2024), and filtered to generate *SV_filtered.VCF* callset. Refinement of these SVs was performed using Iris (v1,0.4) and Jasmine (v1.1.5; Kirsche *et al*., 2023), producing *SV_refined.VCF* file, which was subsequently incorporated into the draft assembly using BCFtools (v1.19) to generate assembly v0.6. The updated assembly was then subjected to two rounds of polishing with NextPolish2 (v0.2.0; Hu *et al*., 2024) using PacBio HiFi read alignments and *k*-mer datasets (*k* = 21, 31) from short NGS reads processed by yak.

### Assembly quality assessment

#### Switch and hamming error

For a trio, the genomic 21-mer libraries were constructed from NGS short reads for each individual (child, father, mother) using *yak count* (https://github.com/lh3/yak) . Switch errors and hamming errors were evaluated for maternal and paternal haploid assembly by using *yak trioeval*. The phasing errors for HGSVC3 assemblies are not available, considering its phasing strategy using Strand-seq data, without parental NGS reads.

#### QV and CV

To estimate the base accuracy and completeness of APGp1 genome assemblies, Merqury (Rhie *et al*., 2020) was employed to calculate the quality value (QV) and completeness value (CV) using NGS and Pacbio HiFi reads. First, *k*-mer databases (*k* = 21) were generated from NGS and PacBio HiFi reads separately using meryl, with singleton *k*-mers (occurring once) filtered out. The two *k*-mer libraries were merged into a hybrid one. Subsequently, QVs were calculated using Merqury with default settings and the CVs were derived from the completeness values calculated by Merqury, following the formulas: QV = -10log_10_(error *k*-mer counts/total *k*-mers) and CV = -10log_10_(1-completeness/100). The QV values for HGSVC3 and HPRCy1 assemblies were previously calculated in the HPRCy1 (Liao *et al*., 2023) and HGSVC3 papers (Logsdon *et al*., 2025), respectively.

#### GCI

To evaluate assembly continuity at the near-T2T level, we applied a newly developed method, GCI, to summarize general continuity levels and report potential assembly issues (Chen *et al*., 2024). We utilized both minimap2 (v2.26-r1175; Li, 2018) and Winnowmap2 (v2.03; Jain *et al*., 2022) to map phased PacBio HiFi (processed via secphase) and phased ONT reads generated with Canu to corresponding haplotype assembly. Subsequently, the Pacbio HiFi and ONT alignments were processed by GCI with the command ‘*python GCI.py -r $mat_asm --hifi ${mat.winnowmap.hifi.bam} ${mat.minimap2.hifi.paf} --nano ${mat.winnowmap.ont.bam} ${mat.minimap2.ont.paf} -t $threads -p -it pdf*’. GCI reported whole-genome and per-chromosome continuity scores, along with candidate assembly issues lacking sufficient support from sequencing reads. Additionally, GCI scores of HGSVC3 and HPRCy1 samples were evaluated for ten randomly selected samples from each panel, representing diverse superpopulation ancestries.

#### VerityMap

We performed diploid-aware error detection using PacBio HiFi reads with VerityMap (v2.1.2; Bzikadze *et al*., 2022), specifying the ‘-d hifi-diploid’ mode for parental haplotype analysis. The raw output (<out_dir>/*_kmers_dist_diff.bed) included all candidate regions with ≥20% discordant read support (encompassing both potential heterozygous sites and errors), we refined error detection by parsing quantitative statistics from <out_dir>/*_errors.tsv files to isolate regions with ≥80% discordant reads.

#### Flagger

Flagger utilizes the alignments of long reads to the diploid assembly, fits the mapping coverage distribution by constructing mixture models, and classify genomic blocks into different categories reflecting assembly accuracy at those locations, including correct regions (haploid) and regions that may contain errors (erroneous, duplicated, collapsed; Liao *et al*., 2023). The Flagger (v0.3.2) pipeline was adopted in evaluating the assembly quality of APGp1 genomes. First, the mapping depths were calculated using *samtools depth* (‘*-aa -Q 0*’) and *depth2cov* from the alignments of phased Pacbio HiFi reads using Winnowmap2 per assembly. To make coverage thresholds more sensitive to local patterns, the genomes were segmented into 10-Mbp windows for evaluation. Due to the biases in sample preparation and sequencing leading to abnormal PacBio HiFi coverage in some satellite regions (Nurk *et al*., 2022), we incorporated and corrected such coverage bias. We calculated the read coverages in Hsat1, Hsat2, and Hsat3 regions (annotated by RepeatMasker) for the APGp1 samples sequenced on both PacBio Sequel II and Revio platforms (**Supplementary Table 1**). These coverages were compared to the whole-genome average and the coverage correction parameters were determined. Ultimately, we applied coverage thresholds of 0.55 for Revio and 1.2 for Sequel II to Hsat2/Hsat3 regions, and no obvious platform-specific bias was observed for HSat1. We updated the assessment using the adjusted coverage parameters for these Hsat2/3 regions and integrated the results. We also refined duplicated component calls using high-confidence mapped regions, as suggested by Flagger, to minimize false positives.

#### Asmgene and Compleasm

To evaluate gene completeness in APGp1 assemblies, we aligned Ensembl gene cDNA sequences from the human reference genome GRCh38 (release 111) to T2T-CHM13, T2T-CN1, and APGp1 assemblies using minimap2 with the command ’*minimap2 -cxsplice:hq $asm Homo_sapiens.GRCh38.cdna.all.fa.gz > $asm.cdna.paf*’. Gene completeness per APGp1 assembly was subsequently evaluated using *paftools asmgene* in minimap2 (v2.26-r1175), specifically for autosomal regions (via the *-a* option). Additionally, gene-level completeness for APGp1 assemblies were further assessed by leveraging compleasm (v0.2.5; Huang *et al*., 2023) with the command ’*compleasm run -a $asm -o compleasm_$asm -l primates -L $LIBRARY_PATH*’.

### Genome annotation

#### Repeat annotation

Repeat elements were initially identified using RepeatMasker (v4.1.2) with the Dfam (v3.3) database under the following settings: sensitive mode (*-s*), species tag for human (*-species human*), and the NCBI BLAST-derived search engine RMBlast (*-e ncbi*). Satellites located in peri/centromeric regions, including αSat, HSat1, HSat2, HSat3, βSat, and γSat, were annotated separately. For αSat, a consensus database was generated for each monomeric class from the T2T-CHM13 genome and identified using megablastn with parameters ‘*-evalue 1e-10 -task megablast*’. The resulting BED file was processed using a custom bash script (https://github.com/Asian-Pan-Genome/Centromere/tree/main/SatelliteAnnotationWorkflow), with filtering for identity < 85% and alignment length > 100 bp. Other types of satellites were annotated following the T2T-CHM13 pipeline (https://github.com/altemose/chm13_hsat) to produce the centromere satellite annotation track.

#### Centromere

Centromeric satellite annotation tracks, including rDNA, were merged using specific distance thresholds: 10 kbp for αSat and 5 kbp for HSat1, HSat2, HSat3, βSat, and γSat. To precisely define pericentromeric and centromeric regions, an iterative extension approach was implemented using a custom script (https://github.com/Asian-Pan-Genome/Centromere). Specifically, the initial start and end positions were defined based on the αSat region, with 5 Mbp flanking spans on both sides. In each iteration, continuous satellite blocks were compared with the current peri-centromeric region. If a block exceeded 10 kbp or resided within 2 Mbp of the current peri-centromeric regions, the boundary was extended to encompass it. Acrocentric short arms were fully incorporated into the final cenSat annotation track. To minimize errors from excessive extension during centromere lengths comparisons across populations, bar-plots of each centromere were generated for manual validation.

#### rDNA sequence annotation

The reference human rDNA unit sequence (Genbank: KY962518.1) was aligned to each assembly using blastn (v2.15.0+) with the parameter ’*-gapopen 3 -gapextend 1*’. The alignments with length > 1 kbp, identity > 85% and *e*-value = 0, were extracted and merged using bedtools (v2.31.1, Quinlan & Hall, 2010) with ’*-d 2000*’. Finally, genomic regions exceeding 40 kbp were designated as rDNA regions. To decompose these regions into individual rDNA copies, we used blastn (*-gapopen 3 -gapextend 1*) to map the ribosomal part (first 13.332-kbp sequence) of the reference KY962518.1 to the rDNA regions. Region aligned to the reference rDNA ribosomal part was identified and the intervals between adjacent regions were extracted as primary rDNA copies. Additionally, the rDNA arrays with assembly gaps were split and the copies >52 kbp were manually curated to determine further splitting.

#### rDNA haplotyping

To analyze rDNA haplotypes, we used nucmer in the Mummer package (v4.0.0rc1; Marçais *et al*., 2018) to align the reference rDNA (KY962518.1) against individual rDNA copies with the parameter ‘*--maxmatch -l 100 -c 1000*’ and the resulting delta files were filtered via delta-filter with ‘*-m -i 90 -l 100*’. SNPs were called using SyRI (v1.7.0) from filtered delta files (Goel *et al*., 2019). For intact rDNA copies, PCA was performed using SNPs by Plink (v1.90b7.2; Chang *et al*., 2015) to determine haplotypes. Additionally, we randomly selected 150 copies per haplotype, aligned them to the reference KY962518.1 sequence with minimap2 (v2.26-r1175) using ‘-axasm10’ and visualized the alignments in IGV (v2.19.1; James *et al*, 2011) to verify haplotype clusters. Furthermore, we constructed a maximum-likelihood phylogenetic tree of the selected rDNA copies with iqtree (v1.6.12; Minh *et al*., 2020) under the TVM+F+R10 model with 1000 bootstrap replicates. For rDNA array comparing between assemblies and raw ONT reads, two samples (C050-CHA-N10-01 and C051-CHA-N11-01) were selected and long reads >250 kbp mapping to the rDNA arrays with at least 5 rDNA copies were extracted, excluding reads with clipping >20% of total length. Focusing on hap1 (a 4.4-kbp deletion) and hap2 (a 1.1-kbp insertion) relative to the reference sequence, dot plots were generated with Gepard (v2.1.0; Krumsiek *et al*., 2007) to identify haplotype positions and validate order consistency between reads and assemblies.

### Gene annotation

#### A hybrid annotation pipeline

To systematically annotate gene models in APGp1 genome assemblies, we integrated three independent methods, including liftover, homology search, and *de novo* prediction (**Supplementary Fig. 15**). Firstly, we employed liftoff (v1.6.3; Shumate *et al*., 2021) to map gene annotations from GRCh38.p14 RefSeq Release 110 to individual APGp1 haplotype assemblies with the command:’*liftoff $asm.chrom.fa $hg38.chrom.fa -sc 0.95 -copies -g $hg38.chrom.gff -polish -o $asm.chrom.liftoff.gff - exclude_partial*’. Secondly, we masked the repeat regions annotated by using RepeatMask, and recruited Exonerate (v2.2.0; Slater *et al*., 2005) to align the UniProtKB/Swiss-Prot protein sequences (release 2024_01) to the masked genome assembly to generate the protein-alignment set, with the command: ’*exonerate --model protein2genome --showvulgar no --showalignment no --showquerygff no --showtargetgff yes --softmasktarget yes --percent 80 --targetchunkid 1 --targetchunktotal 100 -q uniprot_sprot.fasta -t $asm*’. Thirdly, we used Augustus (v3.5.0; Stanke *et al*., 2008) to perform *de novo* gene prediction for the repeat-masked assembly, capturing novel gene structures not covered by above two methods. Finally, only gene annotations with ≥95% overlap in gene, exon, and CDS features between the Exonerate protein-alignment set and Augustus prediction set were retained, forming a complementary annotation dataset to the liftoff annotations. Translated coding sequences of these complementary gene elements were aligned to protein sequences from GRCh38.p14 RefSeq release 110, with the best alignment hit (*e*-value < 1e-5) assigned as functional annotation. Ultimately, the liftoff, homology search and de novo prediction sets were merged, and the longest isoform of each gene was preserved for downstream analysis.

#### Integrating and annotating HPRCy1 and HGSVC3 assemblies

To recruit previous high-quality genomes, we collected the assemblies of HPRCy1 (Liao *et al*., 2023) and HGSVC3 (Logsdon *et al*., 2025), generated by hifiasm and Verkko, respectively. Three samples (HG00733, HG02818 and NA19240) from HPRCy1 were excluded, due to superior assembly quality for the same three samples in HGSVC3. Furthermore, the HG002 assemblies in HPRCy1 were updated to the Q100 version (https://github.com/marbl/HG002). Contigs of each assembly were firstly scaffolded onto chromosomes using RagTag (Alonge *et al*., 2022), followed by uniform application of the APGp1 assembly annotation pipeline (including repeat elements, rDNA arrays and protein-coding gene) to a final of 44 HPRCy1 and 65 HGSVC3 human genomes. During annotation, the Verkko-version assemblies of sample NA18939 showed an anomalously low level of completeness, and thus we switched to its hifiasm assembly. Additionally, NA19240 was omitted from the final gene-associated analyses due to unresolved potential assembly issues on chromosome 6 in both hifiasm and Verkko assemblies.

#### Gene CNV analysis

To assess potential copy number bias introduced by uneven assembly quality, we compared protein-coding gene annotations across 320 APGp1 haplotype-resolved assemblies and 30 HPRCy1 and HGSVC3 assemblies. The CNV diversity was calculated per gene within each of the five major super populations (EAS, AFR, SAS, EUR and AMR) referring to the calculation of traditional nucleotide diversity index (θπ), treating distinct copy numbers as multiple alleles. CNV diversity quantifies the probability of observing two different CN alleles in random two haploids within a given population. A higher CNV diversity value suggests reduced constraint on CNVs to a certain extent. To characterize population stratification of CNVs within EAS, we compared the average copy numbers and CNV diversity between EAS and other superpopulations, prioritizing genes exhibiting extreme values for further investigation.

To disentangle whether APGp1-specific copy gains arise from enhanced assembly contiguity or expanded population sampling, Flagger issue annotations of problematic regions (Collapsed, Duplicated, and Erroneous) from HGSVC3 and HPRCy1 assemblies were lifted to the T2T-CN1 reference. Flagger BED intervals were first projected from contig coordinates to RagTag chromosome-scale scaffolds using sample-matched AGP files for each haplotype. All APGp1-specific gene-gain loci were then evaluated against these coordinate-lifted Flagger annotations. For a given locus where the target gene was annotated in a non-APGp1 haplotype, we examined the full gene body extended by 10 kbp of flanking sequence on both sides. Where the gene was absent in a non-APGp1 assembly, we identified the nearest upstream and downstream protein-coding genes on the same GRCh38-ordered chromosome and assessed the intervening syntenic intergenic interval. Loci were excluded from further evaluation if they lacked both upstream and downstream flanking genes on the same chromosome, had flanking genes assigned to different chromosomes, or produced overlapping flanking intervals. For each retained evaluable interval, we quantified overlap with lifted Flagger issue annotations. A gene-gain locus was designated as harbouring a Flagger issue if the queried gene body or syntenic intergenic interval overlapped any issue interval in at least one non-APGp1 haplotype.

### Pangenome graph construction and evaluation

#### Constructing pangenome graphs

To construct pangenome graphs, we employed minigraph (MG, v0.21-r606; Li *et al*., 2020) and Mimigraph-Cactus (MC, v2.8.2; Hickey *et al*., 2023) with reference assemblies T2T-CN1 v1.0 and T2T-CHM13 v2 serving as backbones, respectively (**Supplementary Table 7**). We did not compare our graphs to those referenced to GRCh38, as such comparisons were already thoroughly investigated by HPRCy1 (Liao *et al*., 2023). In addition to graphs from APGp1 genome assemblies, we constructed integrated graphs combining APGp1, HPRCy1 and HGSVC3 assemblies, as well as specific graphs for EAS genomes from HPRCy1 and HGSVC3. Unanchored contigs and mitochondria assemblies were not included in graph construction. To characterize the genomic structural diversity, we quantified graph complexity for MG graphs by counting the number of graph nodes per 500 kbp.

### Pangenome graph quality assessment

#### Read alignment to the APGp1 MC graphs

We mapped NGS short reads from each APGp1 sample to the APGp1 MC T2T-CN1-referenced pangenome graph using *vg giraffe* (v.1.56.0; Sirén *et al*., 2021), with the command *vg giraffe -R $RG -Z $pref.gbz $pref.min -d $pref.dist -p -f $h1 -f $h2 -t $threads*. The reads with no alignment or with an aligned fraction <99% were filtered out. The output was set as BAM format for downstream variant calling and GAF format for evaluating alignment rates and mapping accuracy, respectively. PacBio HiFi reads were aligned to the MC graph using GraphAligner (v1.0.17; Rautiainen *et al*., 2020) with the option ‘*-x vg*’. Low-quality HiFi alignments were filtered using: (1) keeping only the highest-scoring alignment per read (based on AS value), (2) discarding reads with <80% length aligned to the graph, (3) removing alignments with MAPQ < 1, and (4) excluding alignments with identity <90%. Subsequently, per-node read depths were calculated using *vg pack* (v1.56.0). Mapped nodes/edges per sample were classified as on-target or off-target based on their presence within the sample paths (encoded in P-lines of the GFA files). Additionally, we stratified the mapping on off-target nodes from different genomic regions defined based on the sequence complexity and biomedical significance in the T2T-CN1 reference genome. ‘Easy’ regions represent the most unique genomic sequences in T2T-CN1, annotated by lifting over coordinates from T2T-CHM13 ‘easy’ regions by LiftOver. Pericentromere regions denote centromeres plus 5-Mbp flanking sequences. Segmental duplications (SDs) of T2T-CN1 assembly version v1.0 were annotated using the same methodology used by Yang *et al*. (2023). Low-complexity regions were defined by the simple repeat annotation of RepeatMasker (v4.1.2). Challenging medically relevant genes (CMRGs), KIR and MHC gene loci were extracted from the T2T-CN1 gene annotation file.

#### Graph-based small variant calling

Variant sites decomposed from the MC graph were identified using *vg deconstruct*. Multi-sample VCF files were converted to per-sample VCF files using *bcftools view - a -l -s sample* and multi-allelic sites were split into bi-allelic records using *bcftools norm -m -any*. For each site, the maternal and paternal alleles were merged into diploid genotypes. Leveraging NGS alignments to the graph, variants were called using DeepVariant (v1.6.0; Poplin *et al*., 2018), focusing exclusively on small variants (SNPs and InDels) due to short read limitations. During the DeepVariant procedure, we used the flags *--keep_legacy_allele_counter_behavior* and *--normalize_reads*, and set a minimum mapping quality of 1 in the *make_example* step before variant calling. For these graph mapping-based variant calls, we retained only loci with non-reference genotype (excluding 0/0) and those with ‘PASS’ marks annotated by DeepVariant. Genomic coordinates of variants in both VCF files per sample were then converted to BED format, with the original starting position and ending position extended by the maximal allele length. Highly-repetitive complex regions (including centromere, rDNA, and telomere regions) were excluded from both BED files using *bedtools intersect -v -a variant.bed -b complex_region.bed*. Finally, NGS-DeepVariant graph-based call sets were compared with the pangenome-decoded variants using *bcftools intersect*.

### Pangenome reference bias evaluation

#### Comparing graph-decomposed variants

We used ‘*vg stats -zNEIL*’ to count the numbers and lengths of nodes and edges in the graphs with T2T-CN1 and T2T-CHM13 as backbones, respectively. Variants decomposed from the two APGp1 MC graphs were classified into four categories, including single nucleotide polymorphism/variants (SNPs), structural variants (SVs, ≥50 bp), multiple nucleotide polymorphisms (MNPs) and small insertion-deletions (Indels, <50 bp and ≥1bp), with conflicting annotations resolved by priority: SNP > SV > MNP > InDel. We used LiftOver to compare the variant positions between T2T-CHM13 and T2T-CN1-based calls.Variants were designated as “lift-call” if they satisfied two criteria: successful coordinate conversion via LiftOver under default parameters, and a genomic overlap with the variant callset derived from the alternative reference backbone. To further assess reference bias in allele frequency, we extracted shared SNPs from both decomposed VCFs and compared reference allele frequencies. The structural differences between T2T-CHM13 and T2T-CN1 were analyzed using nucmer (v4.0.0; Marçais *et al*., 2018) with the parameter ‘*—maxmatch -t 5 -l 50 -c 100 -D 1*’.

#### Aligning short and long reads to the graphs

Genomic reads from four EAS individuals were used to investigate reference bias in graph-based read alignment. For NGS data, we utilized 50× short Illumina reads of Chinese individuals HG006 and HG007 from Genome in a Bottle (GIAB). For Pacbio HiFi reads, we selected 20× coverage from YAO (He *et al*., 2023) and 10× coverage from the Chinese Quartet (CQ; Jia *et al*., 2023). Sample reads were aligned against the T2T-CN1 and T2T-CHM13-referenced APGp1 MC pangenome graphs, using *vg giraffe* for NGS reads and GraphAligner for PacBio HiFi reads, respectively. The resulting GAF alignment files were filtered as previously described. Two mapping metrics were assessed: mapping ratio (proportion of reads aligned to the graph), and unreliable mapping ratio (proportion of genomic bases with abnormal depths, defined as <0.1 times or >5 times the expected mapping depth across all nodes).

#### Evaluating multiple pangenome graphs for diverse populations

To comprehensively evaluate NGS short-read alignment performance across diverse human populations, we analyzed the whole-genome sequences from the Human Genome Diversity Project (HGDP; Bergström *et al*., 2020) by aligning them to six pangenome graphs using *vg giraffe*.The HGDP samples spanned 51 globally diverse populations, including seven populations from Africa (Yoruba, San, Mbuti, Mandenka, Biaka, BantuSouthAfrica, and BantuKenya), eight from Europe, three from Middle East, seven from South Asia, 18 from East Asia, three from Oceania, and five from America (**Supplementary Fig. 32**). The six MC pangenome graphs were included T2T-CHM13-referenced HPRCy1 graph with allele filtering parameter d9 (identified as ‘HPRCy1-CHM13-d9’), T2T-CHM13-referenced CPC graph with d12 filtering (CPC-CHM13-d12), T2T-CN1-referenced APGp1 graphs with filtering options of d2 and d32, respectively (APGp1-CN1-d2 and APGp1-CN1-d32), and T2T-CN1-referenced graph for global assemblies with filtering options of d2 and d54 (GLOBALp1-CN1-d2 and GLOBALp1-CN1-d54). The *dN* filtering parameter (e.g., d9) denotes removing the nodes traversed by fewer than N haplotype paths, implemented via ‘*vg clip -d N -m 10000*’ and ‘*vg clip -s*’. For APGp1 (320 assemblies) and GLOBALp1 (540 assemblies), d32 and d54 were chosen (10% threshold) to align with the recommendations of HRPCy1 and CPC graphs.

Besides Mapping Ratio (MR), we further employed another two metrics to quantify high-quality alignments using stringent filtering criteria: the Perfect Mapping Ratio (PMR) and the High-Confidence Mapping Ratio (HCMR). Perfect Mapping Reads are defined as sequencing reads that map entirely to a specific path within the variation graph with zero mismatches and zero gaps (indels). The PMR, representing the proportion of such flawless alignments, was calculated directly via ‘*vg stats -a sample.gam*’. Meanwhile, the HCMR was defined as the proportion of reads satisfying a composite threshold of high-stringency alignment parameters: an aligned fraction exceeding 95% of the total read length, a match fraction exceeding 95% of the aligned length, and a mapping quality (MAPQ) ≥ 30.

To systematically evaluate pangenome graph utility, these three metrics were designed to capture distinct dimensions of alignment performance and structural trade-offs: MR serves as an indicator of overall alignment efficiency, reflecting how effectively a graph backbone accommodates the general sequencing dataset. Broadly, MR is sensitive to graph complexity; a more streamlined graph (e.g., one filtered for rare alleles) typically optimizes MR by reducing ambiguous routing and topological complexity. PMR measures exact sequence homology and is highly sensitive to the preservation of local haplotypes. PMR quantifies a graph’s capacity to retain complete allelic diversity, as the inclusion of fine-scale, population-specific variations directly dictates the proportion of reads that can achieve full-length, error-free alignment. HCMR bridges the gap between overall efficiency and absolute identity, serving as a pragmatic metric for functional graph utility. By allowing minor sequence divergence while strictly filtering out low-quality or structurally ambiguous alignments, HCMR effectively emphasizes the impact of ancestry-specific homology and robust mappability for downstream variants calling.

### Loss-of-function variants

#### Graph-based pLoF variant calling

Given the availability of evolutionary constraint annotation for GRCh38 reference, we constructed pangenome graphs for global assemblies using GRCh38 as graph backbone per chromosome. We generated private variant calls per APGp1 sample. For pangenome-derived variants, the original multi-sample variant sites were firstly converted to per-sample variant format with ‘*bcftools view -a -I -s $sample -c 1:nref*’. The VCF files were further processed using vcfwave and multiallelic sites were split into bi-allelic records using ‘*bcftools norm -c s -f reference.fasta -m -any*’. Multi-nucleotide polymorphisms (MNPs) and complex indels were further decomposed into SNPs and simple InDels by vcfdecompose with parameters ‘*--break-mnps --break-indels*’ from RTG tools (v.3.12.1). Putative Loss-of-function (pLoF) variants were annotated with the LOFTEE plugin in Variant Effect Predictor (VEP, Ensembl Release 112; Karczewski *et al*. 2020). Downstream analyses focused on high-confidence pLoF variants (≤50 bp, tagged LoF=HC), which were further annotated against GnomAD v4.1 database records (Chen *et al*., 2024).

#### HG002 benchmark

To assess the performance of pangenome graph-based pLoF variant calling, we compared HG002 calls against reference variants from GIAB. We firstly annotated high-confidence autosomal pLoF variants for the HG002 v4.2.1 benchmark GIAB callset (Wagner *et al*. 2022) and compared with pangenome graph-derived calls using vcfeval (https://github.com/RealTimeGenomics/rtg-tools). Private pLoF variants between the two sets were manually validated using two publicly released alignments of NGS short reads and PacBio HiFi long reads. Using samtools, 1-kbp local alignments surrounding private pLoF variants were extracted from BAM files and visualized in IGV. Curated flags were applied to all private pLoF calls to mark potential artifacts. Based on local read alignments and original annotations, the flags included (1) pLoF status (stop-gained / frameshift / splicing-donor / splicing-acceptor), (2) curated status (MNP, LCR, assembly error in GRCh38 reference, and ambiguous call) representing false positives.

#### NGS-based pLoF variant calling and comparison

For comparison, we called NGS-based pLoF variants per APGp1 individual. 30× NGS data was aligned to GRCh38 using *bwa mem* and small variants were jointly called using GATK. Variant call accuracy was estimated via Variant Quality Score Recalibration (VQSR) and further filtered using VariantFiltration in GATK for downstream comparison. Shared pLoF variants between the pangenome graph-based and the NGS-based calls were identified per sample using vcfeval and bcftools insect (Danecek *et al*. 2021). Allele frequency of these shared pLoF variants were compared at the population level between the two call sets. To minimize false positives from statistical noise, additional filtering criteria were applied: genotype missing rate < 5%, allele frequency ≤ 0.95 in both call sets, no overlap with low complexity regions, and perfect allelic match at the same position between callsets.

#### Gene category by pLoF

We classified genes containing pLoF variants into four categories, including single heterozygous (with a single pLoF variant), multiple heterozygous (harboring multiple pLoF variants within one of the two parental gene copies), homozygous (carrying one or more homozygous pLoF variants affecting both gene copies, no any heterozygous pLoF variants) and compound (affected by multiple distinct pLoF variants on both gene copies). The compound category was further subdivided into: strict compound with more than one heterozygous pLoF variant on each copy, and loose compound with both heterozygous and homozygous pLoF variants that collectively affect both gene copies.

#### Gene constraint and expression tissue-specificity

To assess gene essentiality across four pLoF variant classes, we utilized two constraint metrics, pLI (the probability of being LoF intolerant; Lek *et al*., 2016) and *s*_het_ (selective effects for heterozygous protein-truncating variants; Zeng *et al*., 2024). Constraint value thresholds for LoF-tolerent genes and genes under strong selection were set at pLI = 0.1 and *s*_het_ = 0.01, following original definitions (Lek *et al*., 2016; Zeng *et al*., 2024), respectively. Additionally, tissue-specificity for pLoF-affected genes across four categories was evaluated using the tau (_τ_) index, a commonly-used indicator of tissue-specific expression. The value of _τ_ ranges from zero (ubiquitous expression) to one (tissue-specific expression). Tau values for all genes were retrieved from GTEx v8 data (Palmer *et al*., 2021), with τ ≥ 0.6 designating high tissue-specificity.

### Structural variations

#### Pangenome graph-based SV calls

Variant sites in the MC graphs were identified using *gfatools bubble* (v0.5) and *vg deconstruct*, respectively. Large spurious deletions were filtered using vcfbub (v.0.1.0) with the option ‘*-l 0 -r 10000000*’. To generate non-redundant SV calls, we adopted PanSVMerger (https://github.com/Asian-Pan-Genome/PanSVMerger) to collapse highly similar paths. For each SV site, decomposed allele sequences were extracted from the graph and subsequently clustered by VSEARCH (v2.30.0; Rognes *et al*., 2016). A new clustering category was defined if any path differed by >50 bp from the longest path. The cluster containing the reference allele was labeled as the reference cluster. Finally, according to the clustering category, the AC (allele count), AF (allele frequency), and AN (allele number) information in the VCF file were updated to reflect non-redundant SVs. This pipeline was applied to prune SVs decomposed from the MC pangenome graphs constructed by HPRC (HPRC.Phase1.CHM13v2) and CPC (CPC.HPRC.Phase1.CHM13v2). These two pangenome graphs revealed a consistent pattern in size distribution after pruning with the results in original studies, and the APGp1 pangenome graph.

#### SV calling based on PacBio HiFi read alignments

Phased PacBio HiFi reads were aligned to the T2T-CN1 reference using Winnowmap2 (v2.03) with the parameters ‘*-x map-pb -a -Y -L --eqx --cs*’ (Jain *et al*., 2022). MD tags required for Sniffles were generated using ‘*samtools calmd*’ and the resulting BAM files were sorted and indexed (Li *et al*., 2009). Three tools were applied to call SVs from the PacBio HiFi read alignments. For PBSV (v2.6.2; https://github.com/PacificBiosciences/pbsv), SV signatures were identified using ‘*pbsv discover*’ with default settings. SVs were then detected using ‘*pbsv call*’ with the parameters ‘*--ccs --preserve-non-acgt -t DEL,INS,INV,DUP,BND -m 40*’. For SVIM (v2.0.0; Heller & Vingron, 2019), SVs were called using ‘*svim alignment*’ with parameters ‘*--read_names --zmws --interspersed_duplications_as_insertions --cluster_max_distance 0.5 --minimum_depth 4 --min_sv_size 40*’, excluding calls with quality <10. For Sniffles (v2.3.2; Smolka *et al*., 2024), SVs were discovered with the parameters ‘*-s 4 -l 40 -n -1 --cluster --ccs_reads*’. Unlike PBSV and SVIM, Sniffles does not generate consensus sequences for insertions from aggregating multiple supporting reads.

#### SV calling based on assemblies

Three methods were recruited to detect SVs from haplotype-resolved assemblies, including SVIM-asm (Heller & Vingron, 2020), PAV (Ebert *et al*., 2021) and LGvar (https://github.com/YafeiMaoLab/LGvar). For SVIM-asm (v1.0.2), assemblies were aligned to T2T-CN1 v1.0 reference using minimap2 (v2.21) with ‘*-x asm5 -a --eqx --cs*’ and then sorted and indexed using samtools. SVs were called using ‘*svim-asm diploid*’ with parameters ‘*--query_names --interspersed_duplications_as_insertions --min_sv_size 40*’. The resulting VCF files were sorted and indexed using BCFtools. For PAV (v0.9.1), assemblies were aligned to the T2T-CN1 v1.0 reference using minimap2 with options ‘*-x asm20 -m 10000 -z 10000,50 -r 50000 --end-bonus=100 --secondary=no -a --eqx -Y -O 5,56 -E 4,1 -B 5*’. SVs were called by using cigar string in the alignments. The LGvar (Large Scale Genetic Variation Identification) pipeline has been documented in Github. Briefly, the assemblies were aligned to the T2T-CN1 v1.0 reference using minimap2 with the options ‘*-cx asm10 --secondary=no --eqx -Y - K 1G -s 1000*’, followed by large indels (>50 bp) detection using the parameters ‘*CLUSTER_FILTER=200000, DESIRED_DEL_LENGTH_ALIGNMENT=300000,CENT_AND_TELO=cts, THREADS_CIGAR_CAL=20*’.

#### Integration of HiFi-based and assembly-based SVs

To integrate SV sets from three HiFi-based and three assembly-based callers to align with pangenome graph-based SV sets, we implemented a two-tier merging strategy: caller-level integration followed by individual-level aggregation. To determine the merging method and threshold used for multiple callers, nine pipelines for merging assembly-based T2T-CHM13-referenced SVs of HG002 were evaluated by comparing with the GIAB SV benchmark set, including jasmine (https://github.com/mkirsche/Jasmine; Kirsche *et al*., 2023), truvari (https://github.com/ACEnglish/truvari; English *et al*., 2022), survivor (https://github.com/fritzsedlazeck/SURVIVOR; Jeffares *et al*., 2017), panpop (https://github.com/starskyzheng/panpop; Zheng *et al*., 2024), bcftools and their combinations (**Supplementary Table 13**). Performance metrics (precision, recall, F1 and genotype congruence) revealed that a mixed strategy of bcftools merge plus SURVIVOR outperformed other pipelines. At the individual level, SURVIVOR was employed to merge SVs across samples. Subsequently, we further clustered adjacent SVs and removed redundancy by using ‘*truvari collapse*’. Final cross-comparison of SV sets was performed using ‘*truvari bench*’.

#### Annotating pangenome graph-derived SVs

The pangenome graph-based SV sites were annotated into diverse sequence categories. Allele sequences from each SV site were extracted as FASTA files, followed by repeat element identification, including transposon and interspersed repeats by RepeatMasker with the NCBI/RMBLAST search engine and Dfam database (v3.3), exact tandem repeats by ETRF (commit fc059d5; https://github.com/lh3/etrf), and centromeric satellites by dna-brnn (v0.1; Li, 2019). SDs were assigned if the total node length of a SV site exceeded 1,000 bp and ≥20% of the bases showed ≥90% similarity to SD annotations in the reference. The sequences were further aligned to T2T-CN1 using minimap2 (v2.24) with options ‘-cxasm20 -r2k --cs’. Adjacent features of the same type within 20 bp were merged. SV sites were classified into diverse repeat classes using a custom pipeline (https://github.com/comery/SvAnnotProcessor/), including VNTR (variable number tandem repeat, unit motif length ≥7 bp), STR (short tandem repeat, unit motif length ≤6 bp), Simple repeat (tandem repeats excluding VNTRs/STRs), SINE/Alu (short interspersed nuclear element), LINE/L1 (long interspersed nuclear element), SVA (SINE-VNTR-Alu), ERV (endogenous retrovirus), other TEs (TEs excluding Alu, L1, SVA, and ERV), and other repeat (mixture of multiple repeat classes).

#### Population-stratified SVs

Population-divergent SVs were identified using pruned graph-derived SV calls with custom scripts. To evaluate whether the identification of APGp1-specific SVs originates improved assembly quality or more sampling, Flagger issue annotations (restricted to regions labeled as Collapsed, Duplicated, or Erroneous) from HGSVC3 and HPRCy1 were lifted onto the T2T-CN1 reference coordinate system using halLiftover (v2.2; Hickey *et al*., 2013) and the HAL file produced by MC. For each APGp1-specific SV, we assessed overlap with lifted Flagger intervals across all non[APGp1 haplotypes. An SV was categorized as affected by a Flagger issue if it overlapped at least one Flagger issue interval in any non[APGp1 haplotype. For functional impact analysis, protein-coding genes within 5 kbp of APGp1-specific SVs were identified using bedtools (v2.31.1) with the following command ‘*bedtools slop -i CN1v1.protein_coding.gff -g CN1v1.fa -b 5000 | bedtools intersect -a - -b APG.SVs.bed*’. Genes overlapping with APGp1-specific or homozygous SVs were analyzed via Gene Ontology (GO) and Kyoto Encyclopedia of Genes and Genomes (KEGG) pathway enrichment with clusterProfiler (v4.8.3; Xu *et al*., 2024) and enrichment for GWAS-associated loci and OMIM diseases with KOBAS (v3.0; Bu *et al*., 2021).

To quantify SVs exhibiting population stratification, we calculated the Hudson Fixation Index (Hudson *F_st_*) among populations using allele frequency per SV site. Compared to *F_st_* metrics of Wright and Nei, Hudson *F_st_* is robust to sample size variation (de Jong *et al*., 2024). Following the definition of Hudson *et al*. (1992), genetic distance between two populations X and Y could be defined as the mean dissimilarity within two populations (nucleotide diversity, _π_*_XY_*) compared to that between the two populations (absolute genetic distance, *D_XY_*). We modified the calculation of nucleotide diversity to be compatible with both bi-allelic and multi-allelic SV sites.

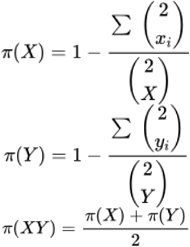

where *x_i_*, *y_i_* are the counts of allele *i* in the two populations, respectively, for a given SV locus, and *X* and *Y* denote the sample sizes of the two populations. The distance estimate between populations is given by the following formula:

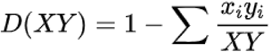

Finally, *HF_st_* is estimated by:

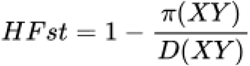

For the SVs with high allele frequency differences between populations, the *HF_st_* values between APGp1 and other EAS samples from HPRCy1 and HGSVC3 (*n* = 30) were used as a control. Only the SVs with top 5% *HF_st_* values (0.288) between EAS and non-EAS superpopulations were retained for further analysis. Within the EAS superpopulation, allele frequencies between populations were additionally compared by calculating the absolute frequency difference per SV site. In the South Chinese Han (CHA-S) versus North Chinese Han (CHA-N) comparison, we classified a SV locus as population-stratified when the frequency difference of reference allele exceeded 0.2, while in the Chinese Han (CHA) *versus* Tibetan (CZA) comparison, an additional requirement of *HF_st_* ≥0. 2 alongside the allele frequency threshold was imposed.

### Large inversions

#### Inversion calling

We identified large inversions (≥10 kbp) across the 538 haplotype assemblies from APGp1, HGSVC3 and HPRCy1, by leveraging three complementary callers, PAV (long-read-based phased assemblies; Ebert *et al*., 2021), SVIM-asm (Structural Variant Identification Method using Genome Assemblies; Heller & Vingron, 2020) and LGvar (https://github.com/YafeiMaoLab/LGvar), to generate inversion callsets against T2T-CHM13 (v2.0) for each assembly. To focus on the high-confidence, balanced inversions on chromosomes arms,we excluded the calls within peri/centromeric regions (defined as the centromere and its 5-Mbp flanking sequences per chromosome in T2T-CHM13), given the challenges in resolving breakpoints accurately due to high-density SDs and recurrent inversion polymorphisms in these regions. Non-redundant inversion calls were derived by integrating results from the three callers using bedtools (Quinlan & Hall, 2010) with the parameters: ‘*bedtools intersect -a callset1.bed -b callset2.bed -f 0.5 -F 0.5 -wa -wb*’. Overlapping inversion calls were merged by a priority hierarchy (PAV, SVIM-asm, LGvar). Regions containing overlapping or nested inversions that could not be resolved into a single event due to sample-specific genomic complexity were classified as “complex regions,” and we prioritized analysis of balanced inversions with clear breakpoints. All inversion breakpoints were manually curated using nucleotide sequence dot plots. For each candidate inversion, we extracted the inversion region plus its 50-kbp flanking sequences from the T2T-CHM13 and identified corresponding regions in assembled haplotypes using minimap2 (2.26-r1175; Li, 2018). Pairwise alignments between extracted regions and their T2T-CHM13 counterparts were generated with Nucmer and visualized via mummerplot (v3.5; Kurtz *et al*, 2004). Breakpoint refinement relied on alignment coordinates, dot plot visualizations, and SafFire (https://github.com/mrvollger/SafFire) for structural validation, ensuring precise breakpoint localization.

#### Benchmarking inversion calls

To assess the performance of our pipeline in calling large inversions, we selected 41 unrelated samples and identified large inversions, comparing our results to a previous study (Porubsky *et al*., 2023) that used multiple sequencing technologies to call inversions in the same cohort (**Supplementary Fig. 40**). The prior study reported 135 large inversions (≥10 kbp) outside centromeric regions. An inversion site is considered orthogonally supported if it shares 50% reciprocal overlap with a site reported in the public dataset. Our pipeline successfully identified 71.9% (97/135) of the previously reported inversions. Of the 38 undetected inversions, majority were primarily attributed to technological limitations, such as those identified via Strand-seq (e.g., chr11:1,751,546-1768,606) or Bionano optical map (e.g., chr17:19,045,523-19,102,543), low-confidence calls in the prior study.

#### Population-stratified inversions

We investigated the EAS-specific large inversions, which are absent from other super populations (AFR, AMR, SAS and EUR). Additionally, we performed Fisher’s exact test to detect population-stratified inversions between EAS and non-EAS super populations, defining statistical significance at a threshold of *P* value < 0.05).

#### Inversion validation

To validate EAS-specific inversion calls, we firstly performed alignments of long reads. Taking the ∼4.3-Mbp inversion at 16q22 as an example, high-quality alignments of phased ONT reads from maternal and paternal haplotypes on T2T-CHM13 were visualized in the IGV (v2.19.1), displaying collapse signals around the breakpoints, as an indicative of inversion. Additionally, we further utilized Bionano optical maps to confirm EAS-specific large inversions larger than 100 kbp. Among the eight such inversions, three at 5p13.2-5q13.3, 7q11.23, 12p13.31 were not verified by Bionano mapping due to insufficient blood DNA for Bionano experiments from their corresponding carriers (C123-CKZ05-01, C118-CMG05-01 and C143-CHU02-01, respectively). Bionano modules were assembled into contigs using Bionano Solve pipeline (v3.8), and the generated contigs were then aligned to the T2T-CHM13 reference. The resulting XMAP alignment files were filtered to retain alignments with a confidence score exceeding 90 and visualized in Bionano Access (https://bionano.com/access-software/).

#### Ultra-large pericentric inversion 5p13.3-5q13.2

For the inversion at 5p13.3-5q13.2, we validated its correctness using a multi-method approach, including sequencing read alignments, Hi-C chromosome interaction analysis, and PCR with Sanger sequencing. Phased PacBio HiFi and ONT reads were aligned against T2T-CHM13, revealing clipping signals in maternal alignments at both inversion breakpoints. For Hi-C data analysis, the short sequencing reads were aligned to T2T-CHM13 and the primary alignments with mapping quality ≥ 30 were kept. Chromatin interactions were visualized as a contact matrix heatmap to assess structural rearrangements. Four pairs of PCR primers were designed based on the breakpoints of the common allele (T2T-CHM13) and inverted allele (C123-CKZ01-01#Mat). PCR cycling conditions included: 95 °C for 5 min; denaturation at 98 °C for 10 s; 35 cycles of amplification with annealing temperatures optimized for each primer pair; elongation at 72 °C for 60 s; and a final elongation step at 72 °C for 5 min. Sanger sequencing of PCR products was performed, and sequences were aligned to T2T-CHM13 to define precise breakpoints. The primer details and Sanger sequencing results were provided in **Supplementary Table 19**. To trace the emergence of this inversion, we mapped the paired-end NGS reads from the mother and father against T2T-CHM13 and highlighted the clipping read pairs at the two inversion breakpoints. PCR validation in parental samples (father C123-CKZ05-02 and mother C123-CKZ05-03) confirmed maternal inheritance of this pericentric inversion.

#### Chromatin topology analysis at 3q29

To characterize the 3q29 inversion’s impact on chromatin architecture, we performed Hi-C analyses on T2T-CHM13 and two homozygous inversion samples (C119-CKZ01-01 and C137-CMH04-01). We utilized HiC-Pro to process and compare Hi-C data based on their corresponding genome assemblies (Servant *et al*., 2015).

#### Analysis of MHC locus

To comprehensively annotate the MHC region, we integrated different approaches. *HLA* and *C4* genes were characterized using Immuannot (commit b50f235; Zhou *et al*., 2024) leveraging the IPD-IMGT/HLA v3.55.0 reference database (Barker *et al*., 2023), which is specially optimized for the annotation of HLA and C4 genes. For other MHC genes, systematic annotation was performed using Liftoff (v1.6.3) with parameters ‘*-sc 0.95 -copies -exclude_partial -polish*’, referencing GRCh38.p14. Notably, three genes, *HLA-DRB2*, *DRB7* and *DRB8*, unannotated on chromosome 6 in the GRCh38 primary assembly, were resolved by lifting from alternative scaffolds using Liftoff. Manual curation was conducted to resolve overlapping gene annotations.

To map complex loci in the MHC region, we first identified SVs larger than 5 kbp, present in at least five assemblies, decomposed from the MC graph using bcftools (v1.21; Danecek *et al*., 2021). For each candidate locus, graph substructures were extracted using odgi (v0.8.6-0-ge647844f; Guarracino *et al*., 2022) and visualized with Bandage (v0.9.0; Wick *et al*., 2015). Gene-graph positional relationships were mapped using the following command ‘*odgi position -i HLA-A.og -E CN1v1.MHC.gff > HLA-A.og.CN1v1.MHC.gff.csv*’. Gene orientation and paths, representing diverse structural haplotypes, were manually annotated and drawn.

#### Structural haplotypes of SMN locus

Recurrent inverted repeats make SMN locus one of the most complicated genomic regions. We resolved the gene and structural haplotypes for genome assemblies from APGp1, HPRCy1 and HGSVC3, with complete SMN sequences. All SMN genes were aligned against the *SMN1* sequence from GRCh38, followed by construction of a maximum-likelihood phylogeny tree using iqtree (1.6.12) with the HKY+F+G4 substitution model and 1000 bootstrap replicates (Minh *et al*., 2020). The diagnostic variant C840T and its linked flanking variants were further examined to validate the distinction between *SMN1* and *SMN2* genes. We analyzed and compared SMN gene copy numbers and strand orientations across individuals and populations. We sought to investigate the structural diversity of SMN locus by employing the pangenome graph approach, however, MC graphs showed poor performance in this inverted region. Thus, we adopted a minimizer-block decomposition strategy to decode the structural haplotypes (sHaps) of SMN locus. We first defined eight principal genomic blocks as minimizer repeat units, by pairwise alignment among human SMN loci. Block coordinates and sequences are available at https://github.com/Asian-Pan-Genome/APGp1/tree/main/Complex_loci/SMN. The eight blocks were present as single copies in chimpanzee and bonobo genomes, suggesting their roles as ancestral building blocks. We decomposed SMN sequences in each assembly by mapping the principal blocks. Given the extensive recurrence of inversions in Palindrome 3-4-3’, we collapsed its structural polymorphism, and assigned sHaps based on block order and orientation. Synteny dotplots between each representative sHap and the T2T-CHM13 allele were visualized using Gepard with a word length of 25 (Krumsiek *et al*., 2007; v2.1).

## Code Availability

All newly developed scripts and pipelines in this study have been deposited at Github (https://github.com/Asian-Pan-Genome/APGp1).

## Data Availability

This study is compliant with the Guidance of the Ministry of Science and Technology of China for the Review and Approval of Human Genetic Resources. Datasets generated in this study have been deposited in the National Genomics Data Center (https://ngdc.cncb.ac.cn) under BioProject accession PRJCA030428, with the approval by the Human Genetic Resources Administration of China (registration numbers 2026BAT00614, 2026BAT00479). Whole-genome short reads and long reads (PacBio HiFi and ONT) are archived at the Genome Sequence Archive under the accession HRA010014. To protect participant confidentiality, the raw sequencing data are only available to the scientific community for general research through a controlled access process. Access can be requested by submitting an application to the Data Access Committee of APG in NGDC. The DAC is committed to reviewing and approving academic research requests within a four-week timeframe. The genome assembly data are publicly available at NGDC (BioProject accession PRJCA030428). Other related resources (including statistic metadata, genomic annotations, pangenome graphs and other supportive files) are available at the APG portal (https://genome.zju.edu.cn/APG) or github repository (https://github.com/Asian-Pan-Genome/APGp1). Genome assemblies of HPRCy1 were obtained from NCBI (https://www.ncbi.nlm.nih.gov/datasets/), and the Verkko-version and hifiasm-version assemblies in HGSVC3 are from https://ftp.1000genomes.ebi.ac.uk/vol1/ftp/data_collections/HGSVC3/working/. The Flagger issue regions in HPRCy1 ans HGSVC3 assemblies were adopted from https://s3-us-west-2.amazonaws.com/human-pangenomics/working/HPRC/$sample/assemblies/year1_f1_assembly_v2_genbank/annotation/flagger/$sample.hifi.flagger_final.simplified.unreliable_only.bed, https://s3-us-west-2.amazonaws.com/human-pangenomics/working/HPRC_PLUS/$sample/assemblies/year1_f1_assembly_v2_genbank/annotation/flagger/$sample.hifi.flagger_final.simplified.unreliable_only.bed and https://ftp.1000genomes.ebi.ac.uk/vol1/ftp/data_collections/HGSVC3/working/20241218_phase3-main-pub_data/uwash/flagger/verkko/final_beds_alt_removed/. The T2T-CN1 assembly (v1.0) is from https://genome.zju.edu.cn/Downloads. GRCh38.p14 primary assembly was downloaded from https://ftp.ebi.ac.uk/pub/databases/gencode/Gencode_human/release_46/. T2T-CHM13 v2 is from NCBI (GCF_009914755.1). PacBio HiFi and ONT data of HGSVC3 were downloaded for quality evaluation (QV and GCI) from https://ftp.1000genomes.ebi.ac.uk/vol1/ftp/data_collections/HGSVC3/working/. WGS short reads of HGDP individuals are from https://ftp.1000genomes.ebi.ac.uk/vol1/ftp/data_collections/HGDP/data/. NGS reads of EAS individuals HG006 and HG007 are accessible at GIAB (https://www.nist.gov/programs-projects/genome-bottle). The Hi-C data of T2T-CHM13 was downloaded from https://s3-us-west-2.amazonaws.com/human-pangenomics/T2T/CHM13/arima/. The previously released human pangenome graph CPC.HPRC.Phase1.CHM13v2 was downloaded from https://pog.fudan.edu.cn/cpc/#/data. The HPRCy1 MC graph is from https://github.com/human-pangenomics/hpp_pangenome_resources. Also gnomAD data was downloaded from https://storage.googleapis.com/gcp-public-data--gnomad/release/4.1/vcf/joint. The BAM files of HG002 aligned to reference GRCh38.p14 were obtained from https://ftp.ncbi.nlm.nih.gov/ReferenceSamples/giab/data/AshkenazimTrio/HG002_NA24385_son/NIST_BGIseq_2x150bp_100x/GRCh38/ and https://ftp-trace.ncbi.nlm.nih.gov/ReferenceSamples/giab/data/AshkenazimTrio/HG002_NA24385_son/PacBio_MtSinai_NIST/PacBio_minimap2_bam/.

## References

Arvanitis, M., et al. (2020) Genome-wide association and multi-omic analyses reveal *ACTN2* as a gene linked to heart failure, Nature Communications, 11(1), p. 1122.

Audano, P.A., et al. (2019) Characterizing the major structural variant alleles of the human genome, Cell, 176(3), pp. 663–675.e19.

Auton, A., et al. (2015) A global reference for human genetic variation, Nature, 526(7571), pp. 68–74.

Backman, J.T. (2016) Role of Cytochrome P450 2C8 in drug metabolism and interactions, Pharmacological Reviews, 68(1), pp. 168–241.

Bergström, A., et al. (2020) Insights into human genetic variation and population history from 929 diverse genomes, Science, 367(6484), p. eaay5012.

Bolognini, D., et al. (2024) Recurrent evolution and selection shape structural diversity at the amylase locus, Nature, pp. 1–9.

Brereton, P. (2001) Pan1b (17βHSD11)-enzymatic activity and distribution in the lung, Molecular and Cellular Endocrinology, 171(1–2), pp. 111–117.

Broutet, N., et al. (2003) Pepsinogen A, pepsinogen C, and gastrin as markers of atrophic chronic gastritis in European dyspeptics, British Journal of Cancer, 88(8), pp. 1239–1247.

Bzikadze, A.V., Mikheenko, A. and Pevzner, P.A. (2022) Fast and accurate mapping of long reads to complete genome assemblies with VerityMap, Genome Research, 32(11–12), pp. 2107–2118.

Chaturvedi, P., Singh, A.P. and Batra, S.K. (2008) Structure, evolution, and biology of the *MUC4* mucin, The FASEB Journal, 22(4), pp. 966–981.

Chen, Q. et al. (2024) GCI: a continuity inspector for complete genome assembly, Bioinformatics, 40(11), p. btae633.

Chin, C.-S., et al. (2023) Multiscale analysis of pangenomes enables improved representation of genomic diversity for repetitive and clinically relevant genes, Nature Methods, 20(8), pp. 1213–1221.

Choy, M.-K. (2010) MICA polymorphism: biology and importance in immunity and disease, Trends in Molecular Medicine, 16(3), pp. 97–106.

de Cid, R., et al. (2009) Deletion of the late cornified envelope *LCE3B* and *LCE3C* genes as a susceptibility factor for psoriasis, Nature Genetics, 41(2), pp. 211–215.

Collins, R.L., et al. (2020) A structural variation reference for medical and population genetics, Nature, 581(7809), pp. 444–451.

Corpas, M., et al. (2025) Bridging genomics’ greatest challenge: The diversity gap, Cell Genomics, 5(1).

D’Antonio, M., et al. (2023) Fine mapping spatiotemporal mechanisms of genetic variants underlying cardiac traits and disease, Nature Communications, 14(1), p. 1132.

Ebert, P., et al. (2021) Haplotype-resolved diverse human genomes and integrated analysis of structural variation, Science, 372(6537), p. eabf7117.

Ebler, J., et al. (2022) Pangenome-based genome inference allows efficient and accurate genotyping across a wide spectrum of variant classes, Nature Genetics, 54(4), pp. 518–525.

Fatumo, S., et al. (2022) A roadmap to increase diversity in genomic studies, Nature Medicine, 28(2), pp. 243–250.

Fu, L., et al. (2026) A long-read human pangenome initiative for comprehensive interpretation of nuclear-embedded mitochondrial DNA, Nature Communications, 17, p. 4371.

Gao, Y., et al. (2023) A pangenome reference of 36 Chinese populations, Nature, 619(7968), pp. 112–121.

Garrison, E., et al. (2024) Building pangenome graphs, Nature Methods, 21(11), pp. 2008–2012.

Ghorbani, M., et al. (2025) Near-complete Middle Eastern genomes refine autozygosity and enhance disease-causing and population-specific variant discovery, Nature Genetics, 57(5), pp. 1119–1131.

Gong, J., et al. (2025) Long-read sequencing of 945 Han individuals identifies structural variants associated with phenotypic diversity and disease susceptibility, Nature Communications, 16(1), p. 1494.

Guarracino, A., et al. (2023) Recombination between heterologous human acrocentric chromosomes, Nature, 617(7960), pp. 335–343.

He, Y., et al. (2023) T2T-YAO: A Telomere-to-Telomere assembled diploid reference genome for Han Chinese, Genomics, Proteomics & Bioinformatics, 21(6), pp. 1085–1100.

He, Y., et al., (2025) Tibetan near-complete pangenome reveals complex variants underlying high-altitude adaptation, bioRxiv, 10.64898/2025.12.16.694547.

Hickey, G., et al. (2024) Pangenome graph construction from genome alignments with Minigraph-Cactus, Nature Biotechnology, 42(4), pp. 663–673.

Hori, Y., Shimamoto, A. and Kobayashi, T. (2021) The human ribosomal DNA array is composed of highly homogenized tandem clusters, Genome Research, 31(11), pp. 1971–1982.

Hsu, L.Y.F., et al. (1987) Chromosomal polymorphisms of 1, 9, 16, and Y in 4 major ethnic groups: A large prenatal study, American Journal of Medical Genetics, 26(1), pp. 95–101.

Huang, N. and Li, H. (2023) Compleasm: a faster and more accurate reimplementation of BUSCO, Bioinformatics, 39(10), p. btad595.

Jadeja, R.N., et al. (2019) Loss of *GPR109A*/*HCAR2* induces aging-associated hepatic steatosis, Aging, 11(2), pp. 386–400.

Jia, P., et al. (2023) Haplotype-resolved assemblies and variant benchmark of a Chinese Quartet, Genome Biology, 24(1), p. 277.

Jiang, T., et al. (2025) SVPG: A pangenome-based structural variant detection approach and rapid augmentation of pangenome graphs with new samples, bioRxiv, 10.1101/2025.07.11.664486.

Karczewski, K.J., et al. (2020) The mutational constraint spectrum quantified from variation in 141,456 humans, Nature, 581(7809), pp. 434–443.

Keinath, M.C., Prior, D.E. and Prior, T.W. (2021) Spinal muscular atrophy: mutations, testing, and clinical relevance, The Application of Clinical Genetics, 14, pp. 11–25.

Lam, C., et al. (2017) Prospective phenotyping of NGLY1-CDDG, the first congenital disorder of deglycosylation, Genetics in Medicine, 19(2), pp. 160–168.

Li, H., Feng, X. and Chu, C. (2020) The design and construction of reference pangenome graphs with minigraph, Genome Biology, 21(1), p. 265.

Lian, Q., et al. (2024) A pan-genome of 69 *Arabidopsis thaliana* accessions reveals a conserved genome structure throughout the global species range, Nature Genetics, 56(5), pp. 982–991.

Liao, W.-W., et al. (2023) A draft human pangenome reference, Nature, 617(7960), pp. 312–324.

Littlefield, C. et al. (2024) A Draft Pacific Ancestry Pangenome Reference. bioRxiv, p. 2024.08.07.606392.

Liu, Q., et al. (2009) Sensory neuron-specific GPCR Mrgprs are itch receptors mediating chloroquine-induced pruritus, Cell, 139(7), pp. 1353–1365.

Liu, J. et al. (2026) Origin and structural evolution of the complex genomic regions of human Y chromosome, under review.

Logsdon, G.A., et al. (2021) The structure, function and evolution of a complete human chromosome 8, Nature, 593(7857), pp. 101–107.

Logsdon, G.A., et al. (2025) Complex genetic variation in nearly complete human genomes. Nature, 644(8076), pp. 430–441.

Ma, M.K., Woo, M.H. and McLeod, H.L. (2002) Genetic basis of drug metabolism. American Journal of Health-System Pharmacy, 59

MacArthur, D.G., et al. (2012) A systematic survey of Loss-of-Function variants in human protein-coding genes, Science, 335(6070), pp. 823–828.

Nassir, N., et al. (2025) A draft UAE-based Arab pangenome reference, Nature Communications, 16, p.6747.

Nozawa, M., Kawahara, Y. and Nei, M. (2007) Genomic drift and copy number variation of sensory receptor genes in humans, Proceedings of the National Academy of Sciences, 104(51), pp. 20421–20426.

Nurk, S., et al. (2022) The complete sequence of a human genome, Science, 376(6588), pp. 44–53.

O’Donnell, S., et al. (2023) Telomere-to-telomere assemblies of 142 strains characterize the genome structural landscape in *Saccharomyces cerevisiae*, Nature Genetics, 55(8), pp. 1390–1399.

Pang, A.W.C., et al. (2013) Mechanisms of formation of structural variation in a fully sequenced human genome, Human Mutation, 34(2), pp. 345–354.

Pirastu, N., et al. (2014) Association analysis of bitter receptor genes in five isolated populations identifies a significant correlation between *TAS2R43* variants and coffee liking, PLOS ONE, 9(3), p. e92065.

Poplin, R., et al. (2018) A universal SNP and small-indel variant caller using deep neural networks, Nature Biotechnology, 36(10), pp. 983–987.

Porubsky, D. (2022) Recurrent inversion polymorphisms in humans associate with genetic instability and genomic disorders, Cell, 185(11), pp. 1986–2005.e26.

Puhl III, H.L., et al. (2015) Human *GPR42* is a transcribed multisite variant that exhibits copy number polymorphism and is functional when heterologously expressed, Scientific Reports, 5(1), p. 12880.

Rautiainen, M. and Marschall, T. (2020) GraphAligner: rapid and versatile sequence-to-graph alignment, Genome Biology, 21(1), p. 253.

Rhie, A., et al. (2023) The complete sequence of a human Y chromosome, Nature, 621(7978), pp. 344–354.

Rosendahl J., et al. (2018) Genome-wide association study identifies inversion in the *CTRB1-CTRB2* locus to modify risk for alcoholic and non-alcoholic chronic pancreatitis, Gut, 67(10), pp. 1855–1863.

Rothschild D. et al. (2024) Diversity of ribosomes at the level of rRNA variation associated with human health and disease, Cell Genomics, 4(9), p. 100629.

Rubio-Gozalbo, M.E., et al. (2021) Galactokinase deficiency: lessons from the GalNet registry, Genetics in Medicine, 23(1), pp. 202–210.

Salm, M.P.A., et al. (2012) The origin, global distribution, and functional impact of the human 8p23 inversion polymorphism, Genome Research, 22(6), pp. 1144–1153.

Schloissnig, S., et al. Structural variation in 1,019 diverse humans based on long-read sequencing, Nature, 644(8076), pp. 442–452.

Sirén, J., et al. (2021) Pangenomics enables genotyping of known structural variants in 5202 diverse genomes, Science, 374(6574), p. abg8871.

Sudmant, P.H., et al. (2015) An integrated map of structural variation in 2,504 human genomes, Nature, 526(7571), pp. 75–81.

Sun, K.Y., et al. (2024) A deep catalogue of protein-coding variation in 983,578 individuals, Nature, 631(8021), pp. 583–592.

Sun, Y. et al. (2026) Multidimensional variation and population stratification across 8000 complete human centromeres, under review.

Suo, M. et al. (2026) Deciphering complete archaic introgression sequences in modern human genomes, under review.

Tang X., et al. (2018) The *CTRB1-CTRB2* risk allele for chronic pancreatitis discovered in European populations does not contribute to disease risk variation in the Chinese population due to near allele fixation, Gut, 67(7), pp. 1368–1369.

The GTEx Consortium (2020) The GTEx consortium atlas of genetic regulatory effects across human tissues, Science, 369(6509), pp. 1318–1330.

Wagner, J., et al. (2022) Benchmarking challenging small variants with linked and long reads, Cell Genomics, 2(5).

Wall, J.D., et al. (2019) The GenomeAsia 100K Project enables genetic discoveries across Asia, Nature, 576(7785), pp. 106–111.

Wang, Q. (2023) 16p11.2 CNV gene Doc2α functions in neurodevelopment and social behaviors through interaction with secretagogin, Cell Reports, 42(7), p. 112691.

Whittle, A.J. (2012) *BMP8B* increases brown adipose tissue thermogenesis through both central and peripheral actions, Cell, 149(4), pp. 871–885.

Xie, L., et al. (2025) Genetic diversity and evolution of rice centromeres. Nature Genetics, 57(11), pp. 2808–2818.

Xue, Y., et al. (2008) Adaptive evolution of *UGT2B17* copy-number variation, The American Journal of Human Genetics, 83(3), pp. 337–346.

Yang, C., et al. (2023) The complete and fully-phased diploid genome of a male Han Chinese, Cell Research, pp. 1–17.

Yang, Q. et al. (2026) Long-read genomics identifies novel risk variants for missing heritability of early-onset schizophrenia, under review.

Yoshizawa, T. (2002) Compound heterozygosity with two novel mutations in the *HEXB* gene produces adult sandhoff disease presenting as a motor neuron disease phenotype, Journal of the Neurological Sciences, 195(2), pp. 129–138.

Zheng, W. et al. (2023) Large-scale genome sequencing redefines the genetic footprints of high-altitude adaptation in Tibetans, Genome Biology, 24(1), p. 73.

Zhou, X. et al. (2010) Copy number variation of *FCGR3A* rather than *FCGR3B* and *FCGR2B* is associated with susceptibility to anti-GBM disease, International Immunology, 22(1), pp. 45–51.

## References for Methods

Alonge, M., et al. (2022) Automated assembly scaffolding using RagTag elevates a new tomato system for high-throughput genome editing, Genome Biology, 23(1), p. 258.

Barker, D.J., et al. (2023) The IPD-IMGT/HLA Database, Nucleic Acids Research, 51(D1), pp. D1053–D1060.

Bu, D., et al. (2021) KOBAS-i: intelligent prioritization and exploratory visualization of biological functions for gene enrichment analysis, Nucleic Acids Research, 49(W1), pp. W317–W325.

Chang, C.C., et al. (2015) Second-generation PLINK: rising to the challenge of larger and richer datasets, GigaScience, 4(1), pp. s13742-015-0047–8.

Chen, S., et al. (2018) fastp: an ultra-fast all-in-one FASTQ preprocessor, Bioinformatics, 34(17), pp. i884–i890.

Chen, S., et al. (2024) A genomic mutational constraint map using variation in 76,156 human genomes, Nature, 625(7993), pp. 92–100.

Danecek, P., et al. (2021) Twelve years of SAMtools and BCFtools, GigaScience, 10(2), p. giab008.

English, A.C., et al. (2022) Truvari: refined structural variant comparison preserves allelic diversity, Genome Biology, 23(1), p. 271.

Goel, M., et al. (2019) SyRI: finding genomic rearrangements and local sequence differences from whole-genome assemblies, Genome Biology, 20(1), p. 277.

Guarracino, A., et al. (2022) ODGI: understanding pangenome graphs, Bioinformatics, 38(13), pp. 3319–3326.

Heller, D. and Vingron, M. (2019) SVIM: structural variant identification using mapped long reads, Bioinformatics, 35(17), pp. 2907–2915.

Heller, D. and Vingron, M. (2020) SVIM-asm: structural variant detection from haploid and diploid genome assemblies, Bioinformatics, 36(22-23), pp. 5519–5521.

Hickey, G. et al. (2013) HAL: a hierarchical format for storing and analyzing multiple genome alignments, Bioinformatics, 29(10), pp. 1341–1342.

Hinrichs, A.S., et al. (2006) The UCSC Genome Browser Database: update 2006, Nucleic Acids Research, 34(suppl_1), pp. D590–D598.

Hu, J., et al. (2024) NextPolish2: A Repeat-aware Polishing Tool for Genomes Assembled Using HiFi Long Reads, Genomics, Proteomics & Bioinformatics, 22(i1), p. qzad009.

Huang, N. and Li, H. (2023) compleasm: a faster and more accurate reimplementation of BUSCO, Bioinformatics, 39(10), p. btad595.

Hudson, R.R., Slatkin, M. and Maddison, W.P. (1992) Estimation of levels of gene flow from DNA sequence data, Genetics, 132(2), pp. 583–589

Jain, C., et al. (2022) Long-read mapping to repetitive reference sequences using Winnowmap2, Nature Methods, 19(6), pp. 705–710.

Jeffares, D.C., et al. (2017) Transient structural variations have strong effects on quantitative traits and reproductive isolation in fission yeast, Nature Communications, 8(1), p. 14061.

Jong, M.J. de, et al. (2024) Calculating and interpreting FST in the genomics era. bioRxiv, p. 2024.09.24.614506.

Kirsche, M., et al. (2023) Jasmine and Iris: population-scale structural variant comparison and analysis, Nature Methods, 20(3), pp. 408–417.

Krumsiek, J., Arnold, R. and Rattei, T. (2007) Gepard: a rapid and sensitive tool for creating dotplots on genome scale, Bioinformatics, 23(8), pp. 1026–1028.

Kurtz, S., et al. (2004) Versatile and open software for comparing large genomes, Genome Biology, 5(2), p. R12.

Lek, M., et al. (2016) Analysis of protein-coding genetic variation in 60,706 humans, Nature, 536(7616), pp. 285–291.

Li, H., et al. (2009) The Sequence Alignment/Map format and SAMtools, Bioinformatics, 25(16), pp. 2078–2079.

Li, H. (2013) Aligning sequence reads, clone sequences and assembly contigs with BWA-MEM. *arXiv*.

Li, H. (2018) Minimap2: pairwise alignment for nucleotide sequences, Bioinformatics, 34(18), pp. 3094–3100.

Li, H. (2019) Identifying centromeric satellites with dna-brnn, Bioinformatics, 35(21), pp. 4408–4410.

Marçais, G., et al. (2018) MUMmer4: A fast and versatile genome alignment system, PLOS Computational Biology, 14(1), p. e1005944.

Minh, B.Q., et al. (2020) IQ-TREE 2: New Models and Efficient Methods for Phylogenetic Inference in the Genomic Era, Molecular Biology and Evolution, 37(5), pp. 1530–1534.

Nurk, S., et al. (2020) HiCanu: accurate assembly of segmental duplications, satellites, and allelic variants from high-fidelity long reads, Genome Research, 30(9), pp. 1291–1305.

Palmer, D., et al. (2021) Ageing transcriptome meta-analysis reveals similarities and differences between key mammalian tissues, Aging, 13(3), pp. 3313–3341.

Porubsky, D., et al. (2023) Inversion polymorphism in a complete human genome assembly, Genome Biology, 24(1), p. 100.

Quinlan, A.R. and Hall, I.M. (2010) BEDTools: a flexible suite of utilities for comparing genomic features, Bioinformatics, 26(6), pp. 841–842.

Rhie, A., et al. (2020) Merqury: reference-free quality, completeness, and phasing assessment for genome assemblies, Genome Biology, 21(1), p. 245.

Robinson, J.T., et al. (2011) Integrative genomics viewer, Nature Biotechnology, 29(1), pp. 24–26.

Rognes, T., et al. (2016) VSEARCH: a versatile open source tool for metagenomics, PeerJ, 4, p. e2584.

Servant, N., et al. (2015) HiC-Pro: an optimized and flexible pipeline for Hi-C data processing, Genome Biology, 16(1), p. 259.

Shumate, A. and Salzberg, S.L. (2021) Liftoff: accurate mapping of gene annotations, Bioinformatics, 37(12), pp. 1639–1643.

Sim, S.B., et al. (2022) HiFiAdapterFilt, a memory efficient read processing pipeline, prevents occurrence of adapter sequence in PacBio HiFi reads and their negative impacts on genome assembly, BMC Genomics, 23(1), p. 157.

Slater, G.S.C. and Birney, E. (2005) Automated generation of heuristics for biological sequence comparison, BMC Bioinformatics, 6(1), p. 31.

Smolka, M., et al. (2024) Detection of mosaic and population-level structural variants with Sniffles2, Nature Biotechnology, 42(10), pp. 1571–1580.

Stanke, M., et al. (2008) Using native and syntenically mapped cDNA alignments to improve de novo gene finding, Bioinformatics, 24(5), pp. 637–644.

Van der Auwera, G.A., et al. (2013) From FastQ Data to High-Confidence Variant Calls: The Genome Analysis Toolkit Best Practices Pipeline, Current Protocols in Bioinformatics, 43(1), p. 11.10.1-11.10.33.

Wick, R.R., et al. (2015) Bandage: interactive visualization of *de novo* genome assemblies, Bioinformatics, 31(20), pp. 3350–3352.

Xu, S. et al. (2024) Using clusterProfiler to characterize multiomics data, Nature Protocols, 19(11), pp. 3292–3320.

Zeng, T., et al. (2024) Bayesian estimation of gene constraint from an evolutionary model with gene features, Nature Genetics, 56(8), pp. 1632–1643.

Zheng, Z., et al. (2024) A sequence-aware merger of genomic structural variations at population scale, Nature Communications, 15(1), p. 960.

Zhou, Y., Song, L. and Li, H. (2024) Full resolution HLA and KIR gene annotations for human genome assemblies, *Genome Research*, p. gr.278985.124.

