## Supplementary Figures for "Hidden genetic diversity in 320 nearly-complete East Asian genome assemblies"

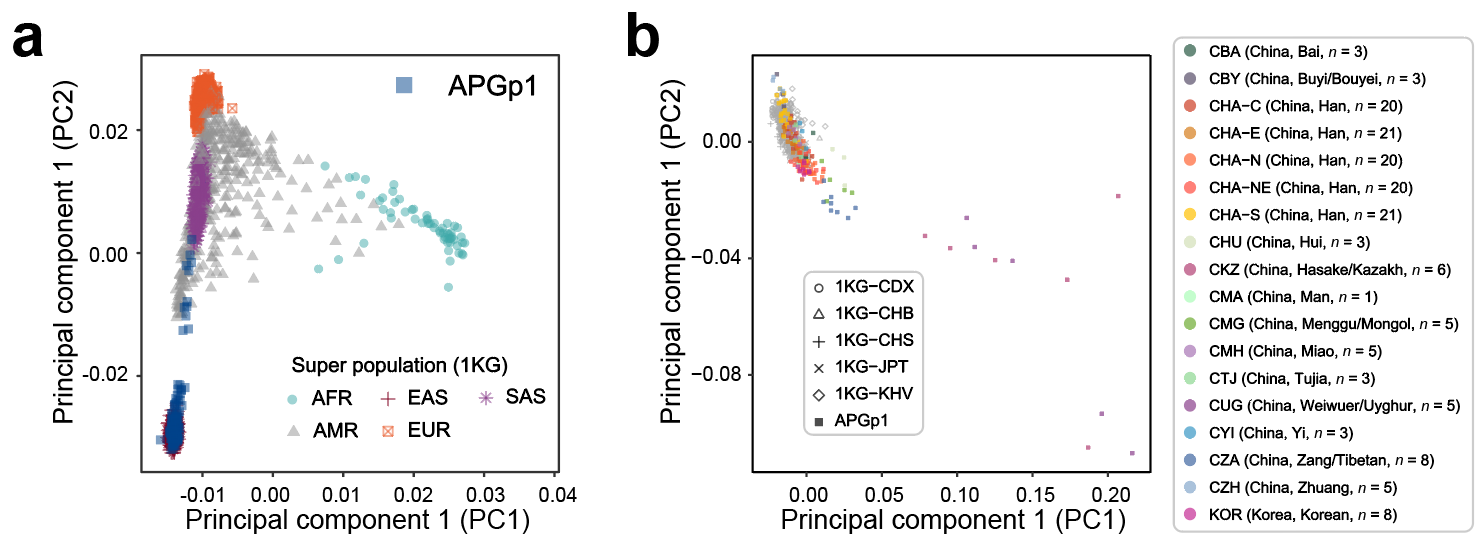

**Supplementary Fig. 1 | Genetic diversity of East Asian genomes in APG phase 1 (APGp1)**. **a**, First two components of 1KGP and APGp1 samples (blue squares), inferred from the principal component analysis (PCA) using SNPs on chromosome 1. 1KGP superpopulation labels: AFR, African ancestry; EAS, East Asian ancestry; SAS, South Asian ancestry; AMR, Admixed American ancestry; EUR, European ancestry. **b**, PCA of East Asian samples from different populations in 1KGP (gray) and APGp1. Population labels: 1KG-CDX, Chinese Dai in Xishuangbanna, China; 1KG-CHB, Han Chinese in Beijing, China; 1KG-CHS, Han Chinese South; 1KG-JPT, Japanese in Tokyo, Japan; 1KG-KHV, Kinh in Ho Chi Minh City, Vietnam; CBA, Bai, China; CBY, Buyi/Bouyei, China; CHA-C, Han Chinese from Central China; CHA-E, Han Chinese from East China; CHA-N, Han Chinese from North China; CHA-NE, Han Chinese from Northeast China; CHA-S, Han Chinese from South China; CHU, Hui, China; CKZ, Hasake/Kazakh, China; CMA, Man/Manchu, China; CMG, Menggu/Mongol, China; CMH, Miao, China; CTJ, Tujia, China; CUG, Weiwuer/Uyghur, China; CYI, Yi, China; CZA, Zang/Tibetan, China; CZH, Zhuang, China; KOR, South Korean.

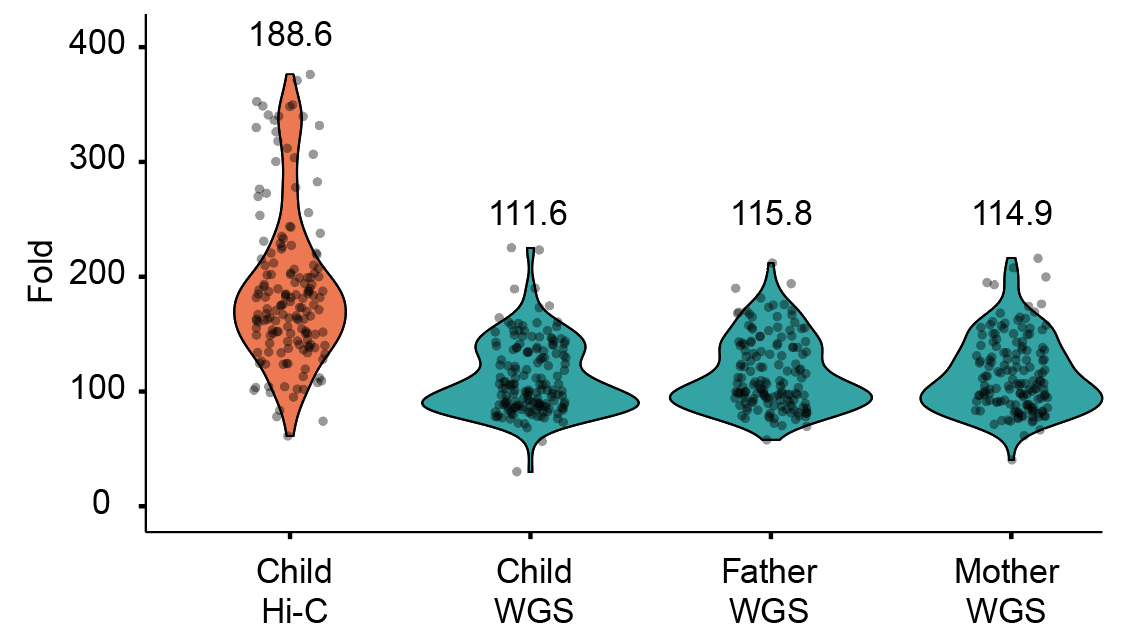

**Supplementary Fig. 2 | Sequencing coverage of NGS short reads for Hi-C and whole-genome sequencing (WGS) in APGp1 samples.** A human reference genome size of 3 Gbp is used for sequencing depth estimation. Average coverages are shown.

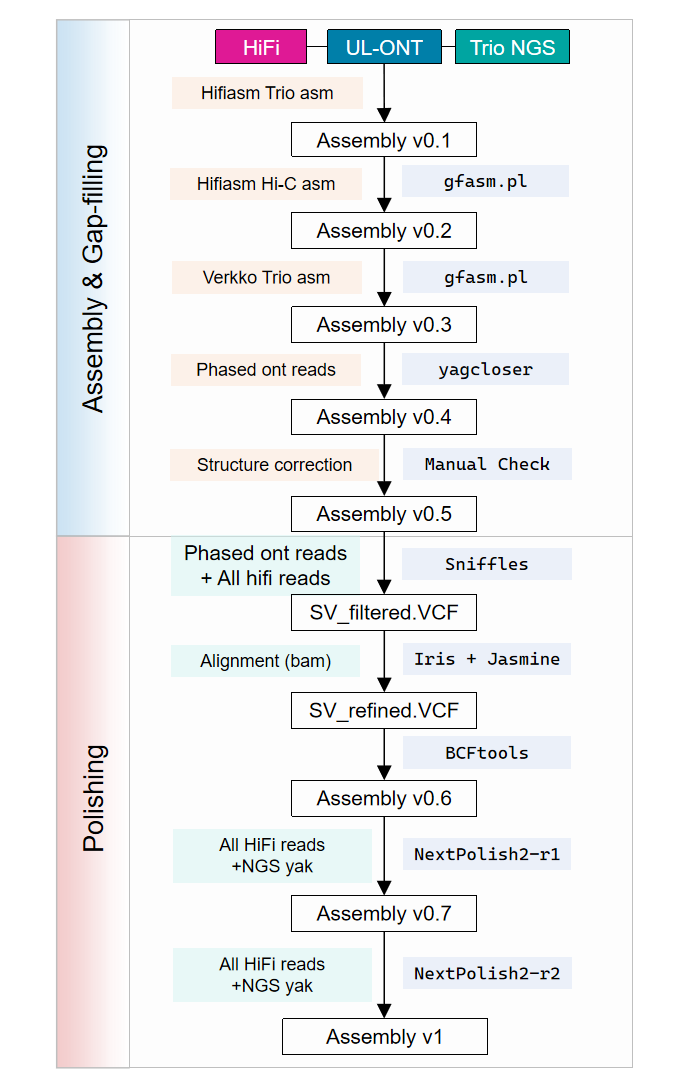

**Supplementary Fig. 3 | Primary workflow for genome assembly, gap filling and polishing in APGp1 for individuals with trio data.** For APGp1 samples with trio information, multiple phased assembly versions are generated using long reads (PacBio HiFi and ultra-long ONT) and phasing data (trio and Hi-C), followed by multiple rounds of gap filling and polishing.

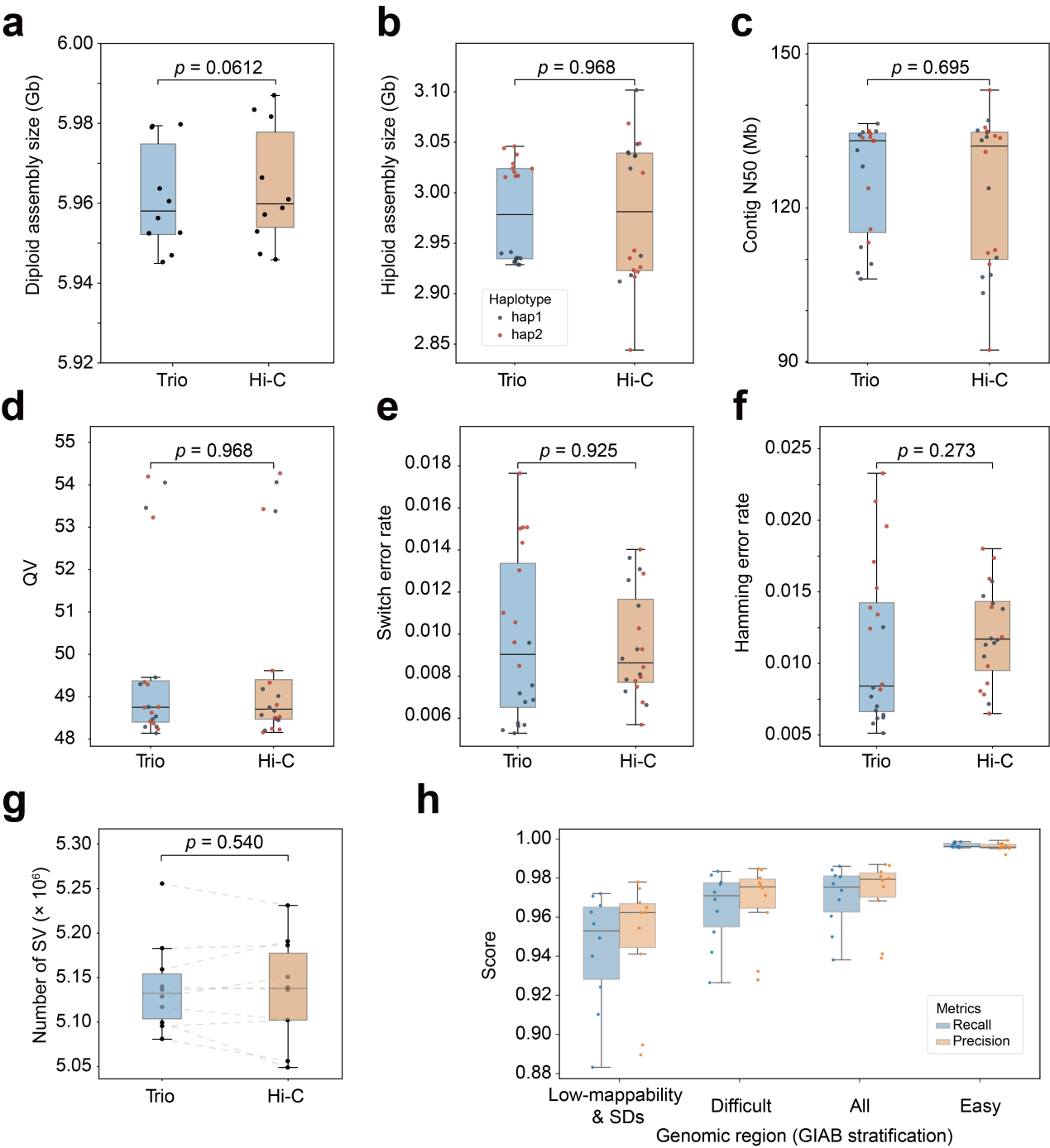

**Supplementary Fig.4 | Comparison between assemblies using trio-based and Hi-C-based phasing strategies.** Haplotype-resolved assemblies of ten individuals using two strategies in hifiasm with available trio and Hi-C data. **a**, diploid assembly sizes. The significance is calculated using a paired *t*-test. **b**, haploid assembly sizes (Wilcoxon rank sum test). **c**, contig N50. **d**, QV. **e**, switch error rates. **f**, hamming error rates. **g**, total SVs called by LSGvar. **h**, SV consistency across GIAB-defined genomic stratification regions, compared to final SV calls for each individual.

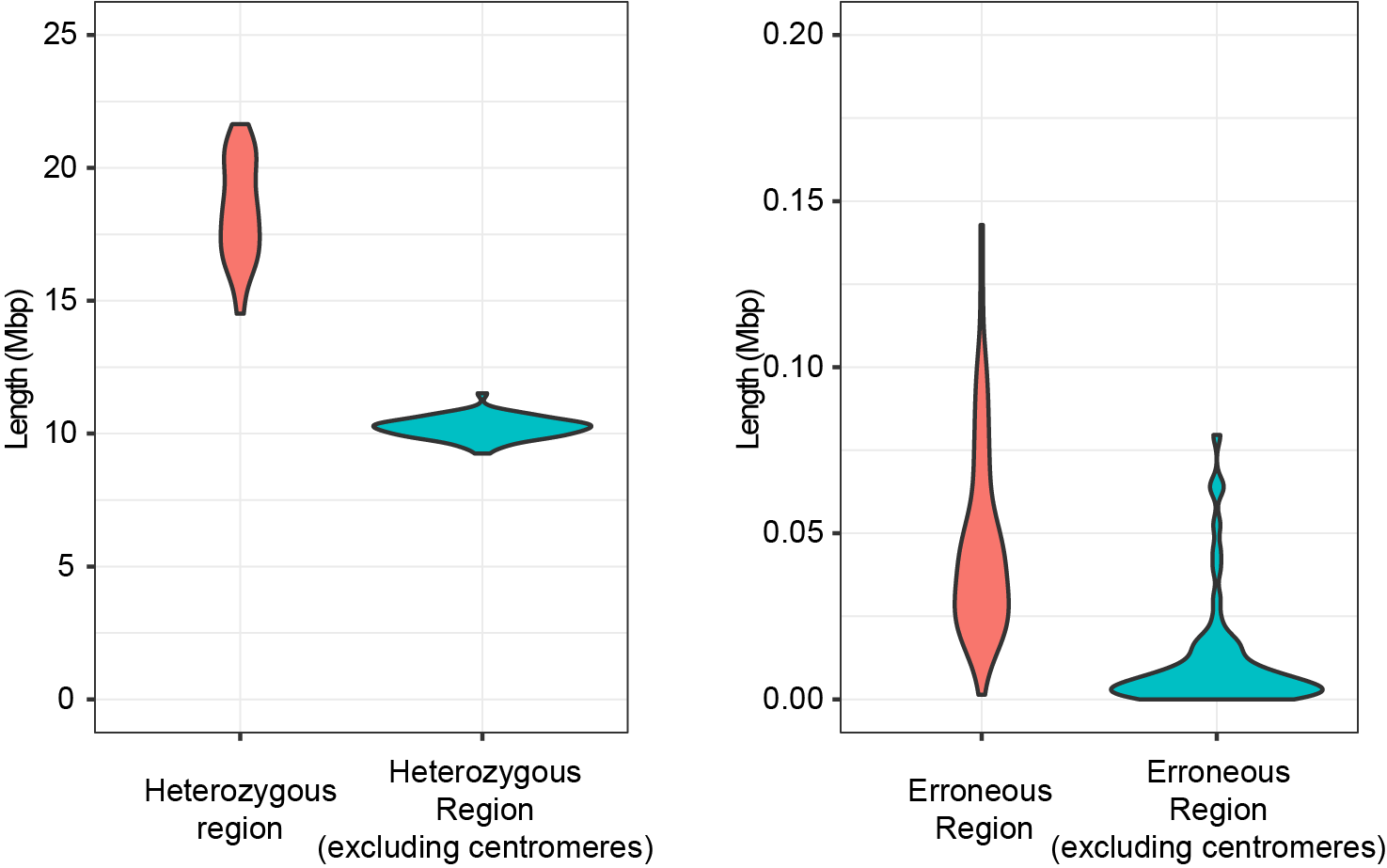

**Supplementary Fig. 5 | VerityMap evaluation of potential heterozygous and erroneous assembly regions.** VerityMap calculates the distances between consecutive rare *k*-mers in the assembly and in a PacBio HiFi read (https://github.com/ablab/VerityMap). Regions with 20%-80% deviated reads are flagged as heterozygous sites, while those with above 80% deviated reads indicate assembly errors. Potential issue regions (heterozygous and erroneous) are counted in 200-bp sliding windows and merged.

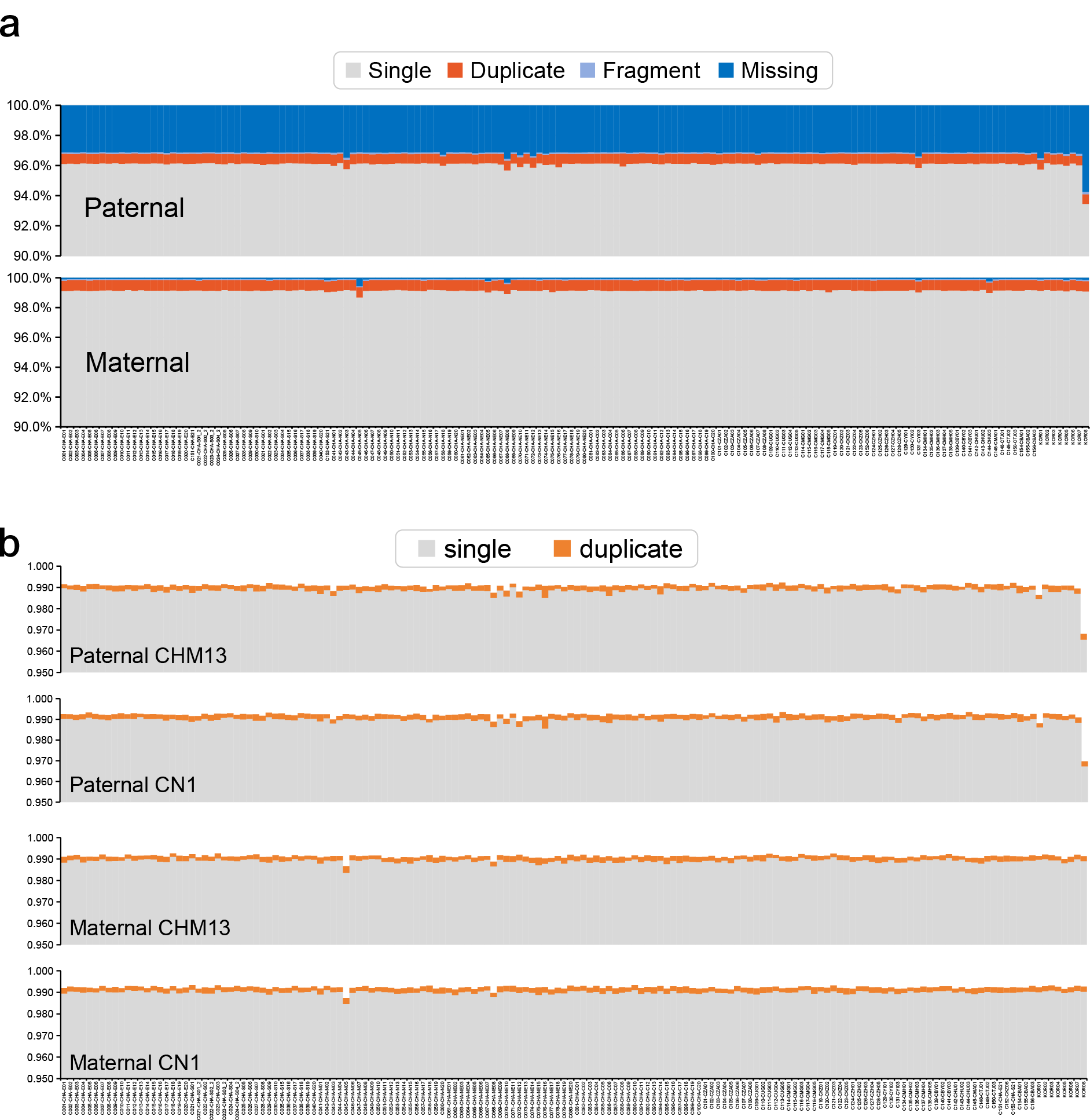

**Supplementary Fig. 6 | Assembly completeness evaluation using reference gene sets. a**, Gene completeness (percentage of BUSCO-like orthologs) for each haplotype assembly, assessed by compleasm (https://github.com/huangnengCSU/compleasm). Paternal assemblies have a ~3% missing rate due to chromosome Y. **b**, Assembly completeness evaluated via asmgene (https://github.com/lh3/minimap2). Potential assembly collapses are evaluated by aligning to T2T-CN1 and T2T-CHM13 gene references, respectively.

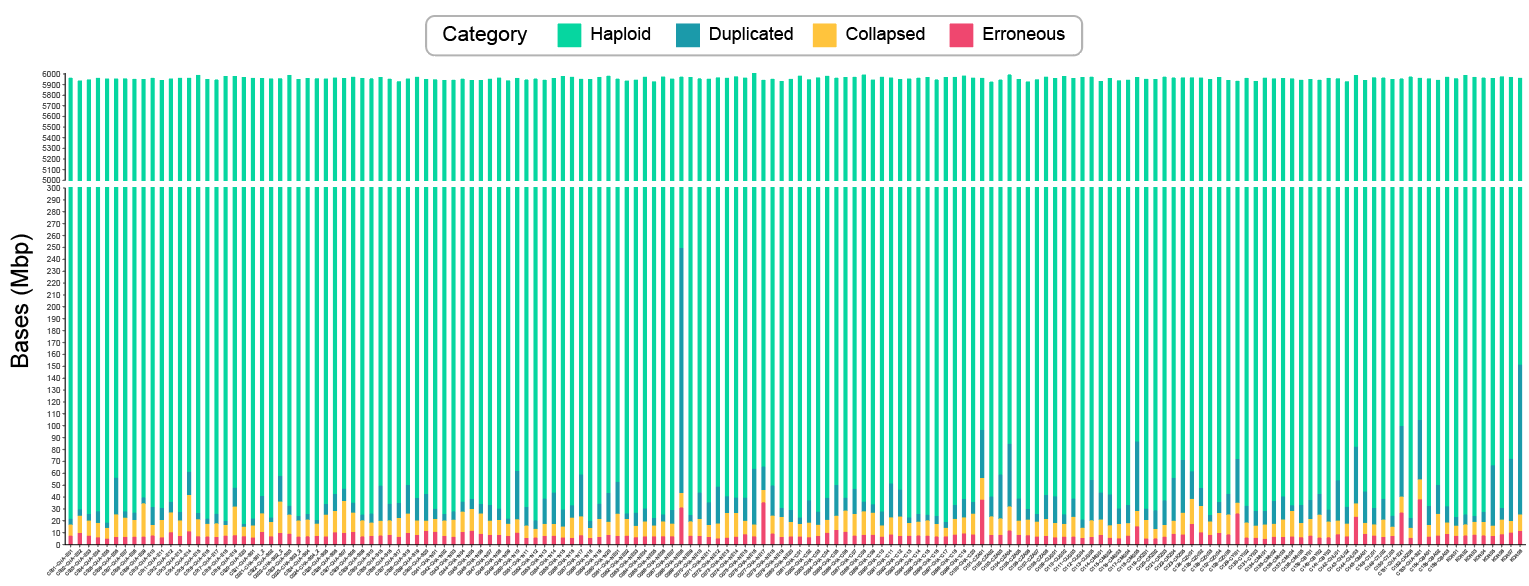

**Supplementary Fig. 7 | Assembly reliability evaluation on 160 diploid APGp1 genomes using Flagger.** The maternal and paternal assemblies for each individual are merged as a diploid for evaluation.

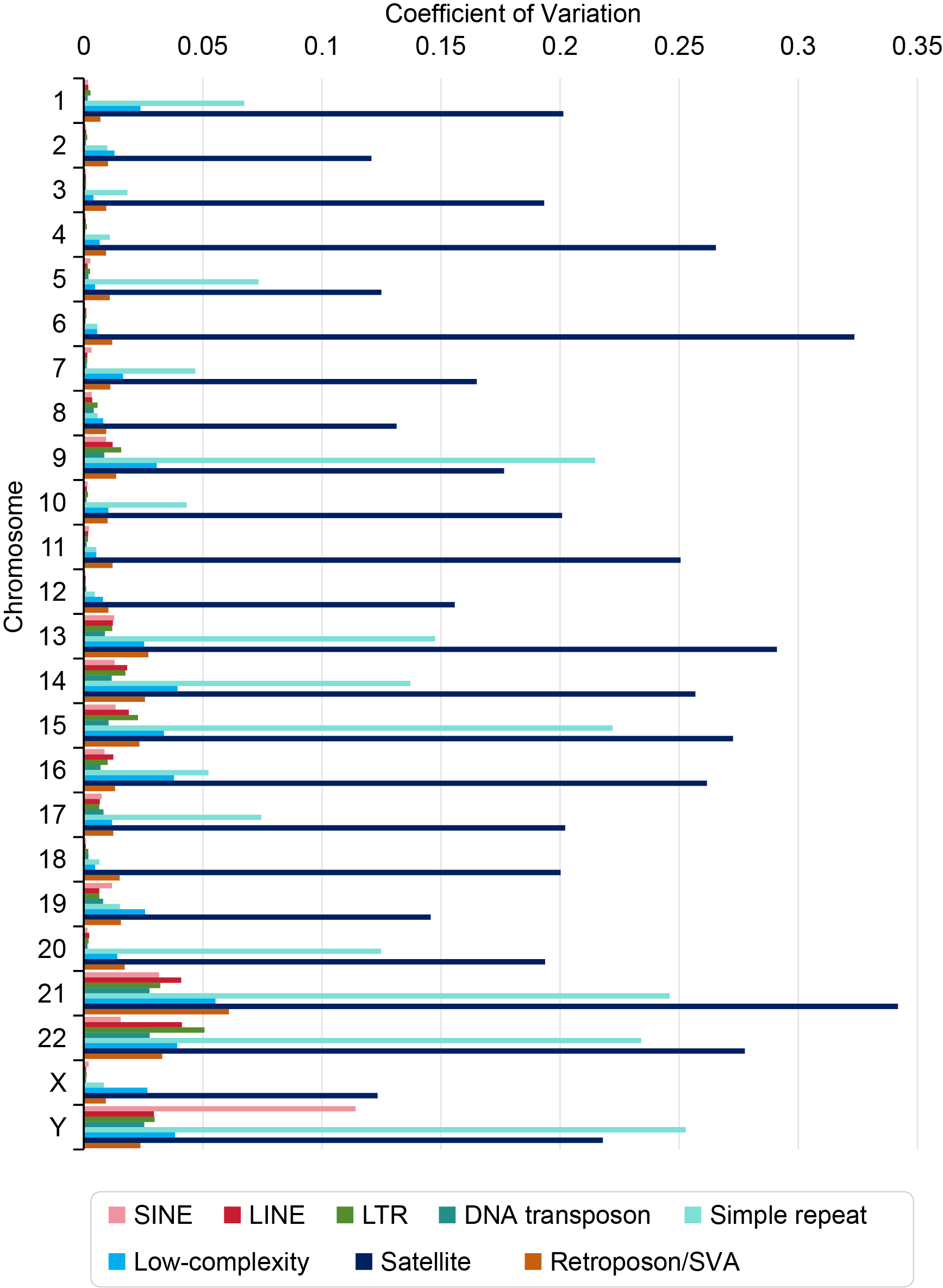

**Supplementary Fig. 8 | Coefficient of variation (CV) in sequence sizes of different repeat elements across APGp1 assemblies.** The different types pf repeat elements are annotated by RepeatMasker (v4.1.2).

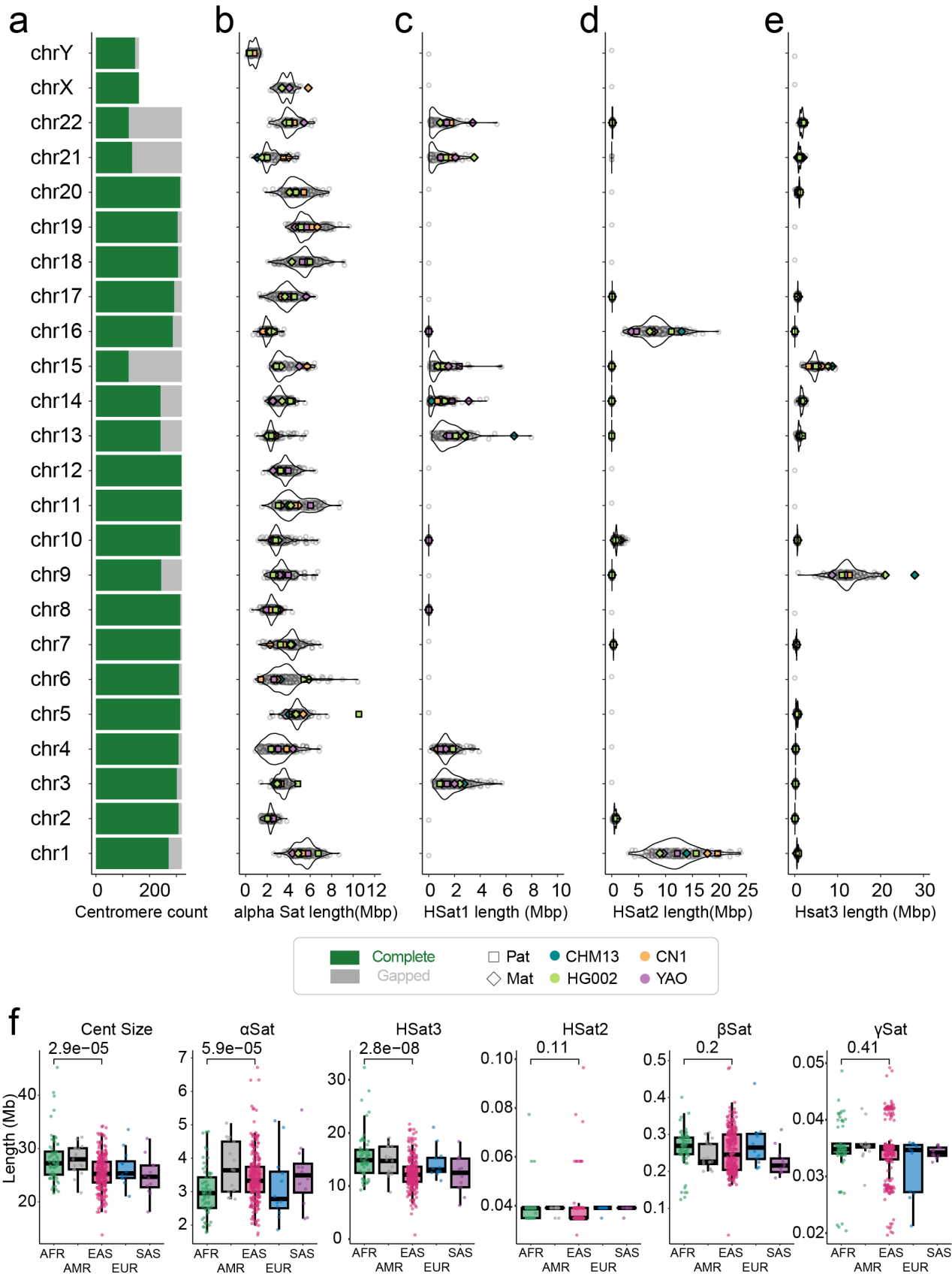

**Supplementary Fig. 9 | Centromere size diversity in EAS genomes. a**, Completeness of centromere assembly per chromosome. Green bars for each chromosome indicate the number of complete centromere assemblies with no internal gaps. **b-e**, Total lengths of alpha satellite (**b**), Hsat1 (**c**), Hsat2 (**d**) and Hsat3 (**e**) in centromeres across EAS genome assemblies. Previous human reference-level assemblies are included and highlighted (phased assemblies of T2T-CN1, YAO and HG002, and haploid assembly T2T-CHM13). Yq12 region from chromosome Y is not included here, although it is predominantly composed of human satellites. **f**, Satellite composition of centromeres on chromosome 9 across five superpopulations. Statistic significance is calculated using Wilcox test.

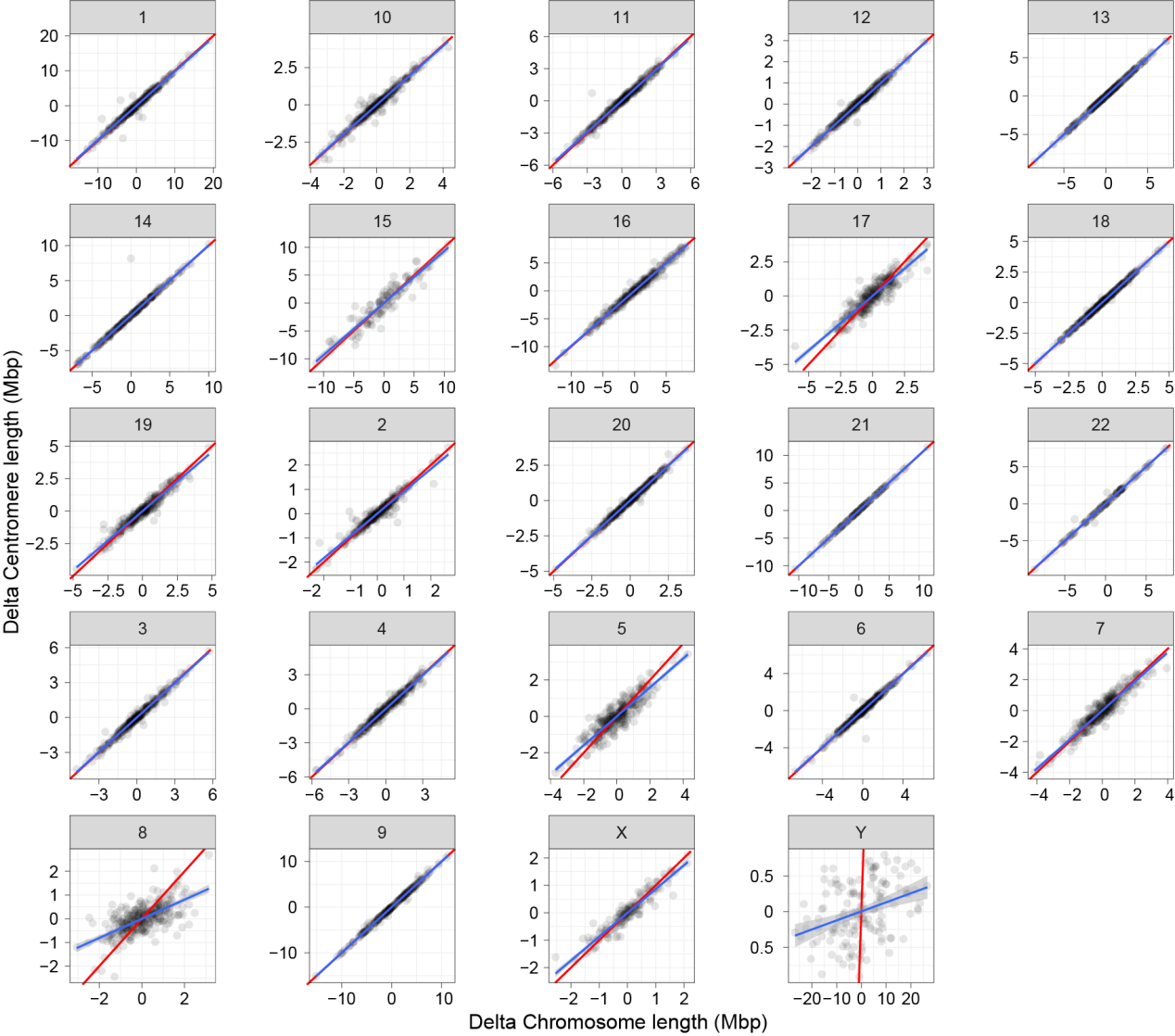

**Supplementary Fig. 10 | Variation in centromere and whole chromosome sizes per chromosome.** Delta chromosome (*x*-axis) and centromere (*y*-axis)length represent size differences. Blue lines denote fitting curves, and red lines denote the *y* = *x* reference.

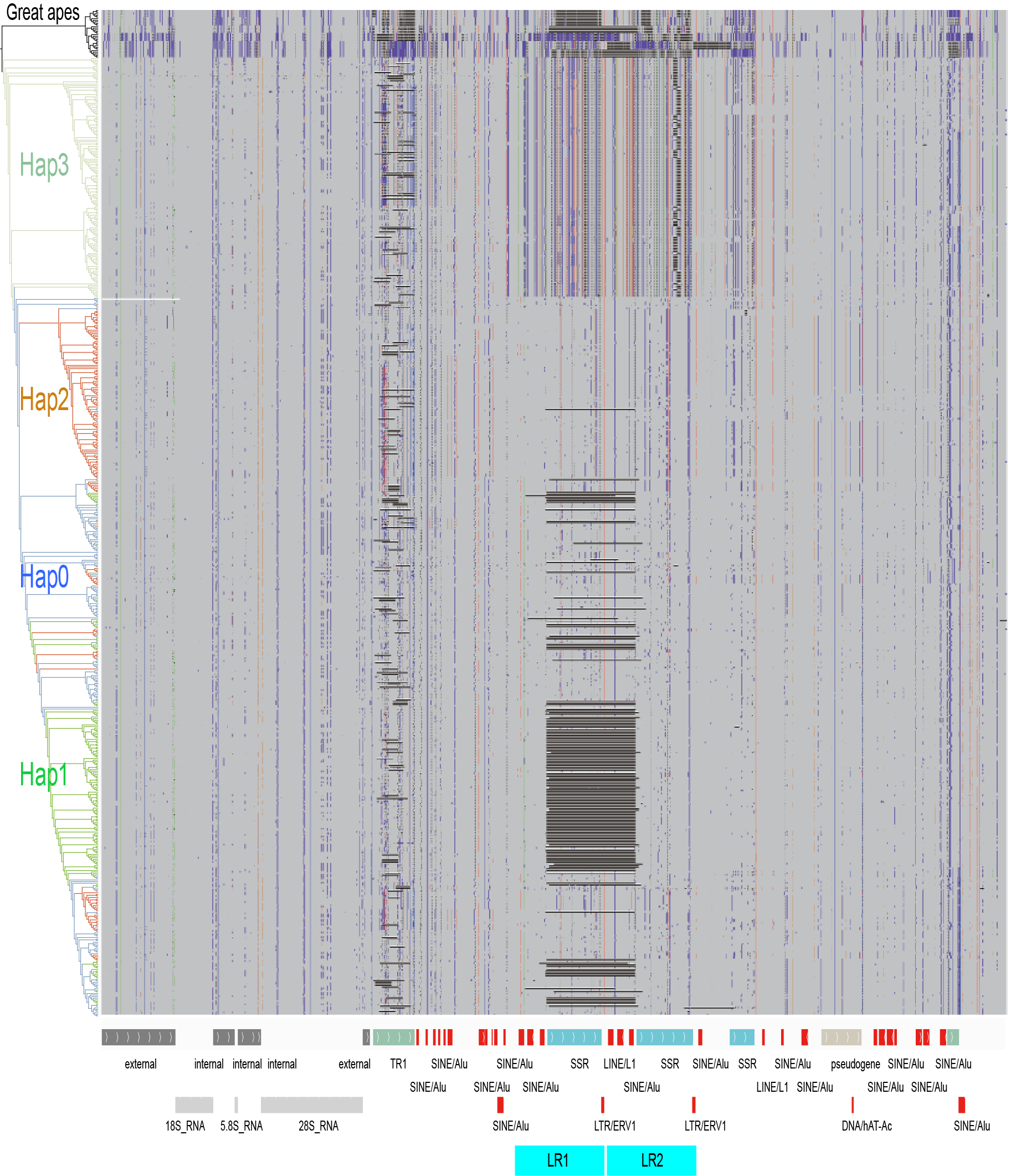

**Supplementary Fig. 11 | Variation and phylogeny of human rDNA sequences.** Intact rDNA sequences from phased T2T assemblies of non-human primate genomes (including chimpanzee, bonobo, gorilla, Sumatran orangutan, Bornean orangutan and siamang gibbon) serve as outgroup copies. Left, a maximum-likelihood phylogenetic tree of representative intact rDNA sequences built using iq-tree under the TVM+F+R10 model. Right, IGV screenshots revealing their alignments against the reference rDNA sequence KY962518. Beneath, repeat element annotation.

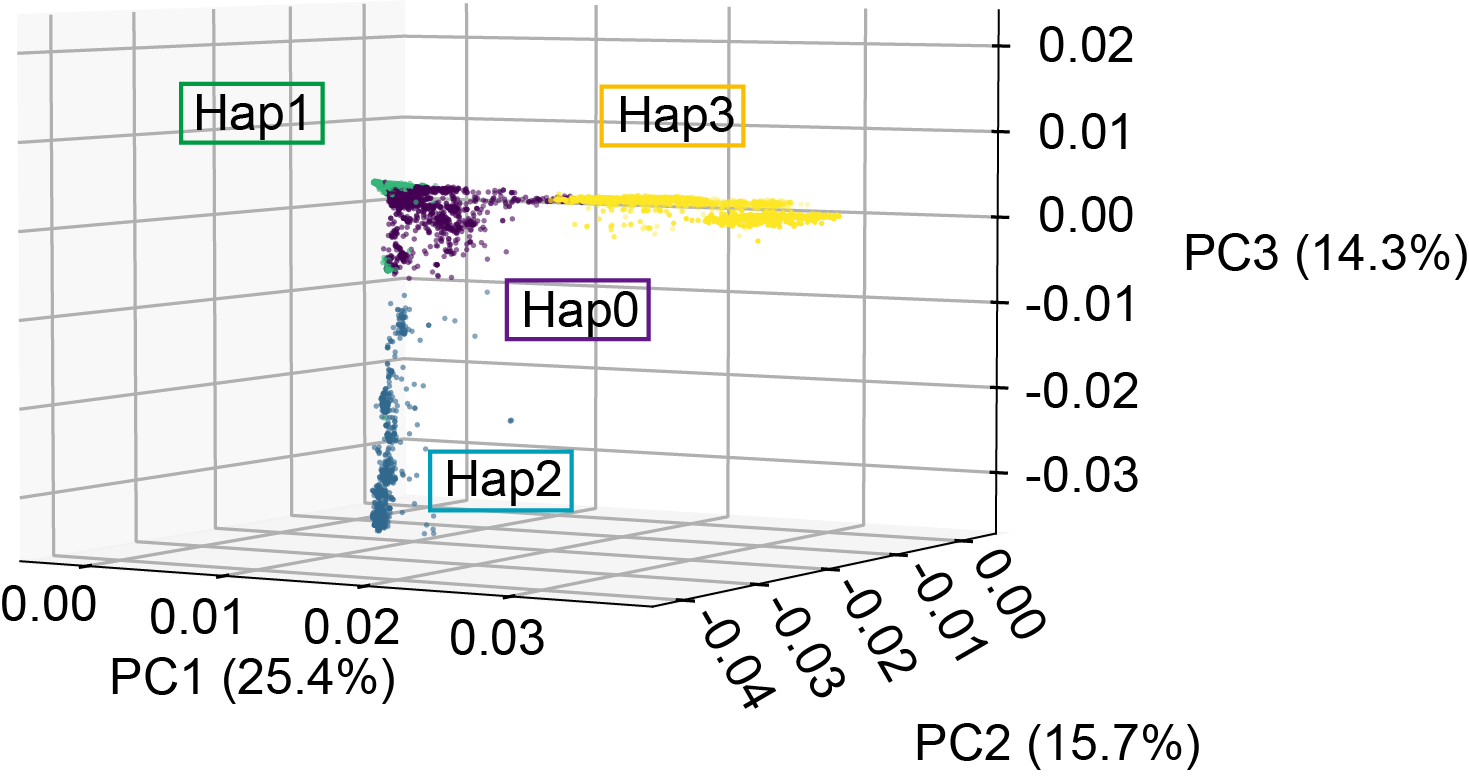

**Supplementary Fig. 12 | 3-dimention PCA plot of human rDNA elements.** Intact rDNA arrays are included in the PCA analysis by using Plink. The first three principal components are plotted.

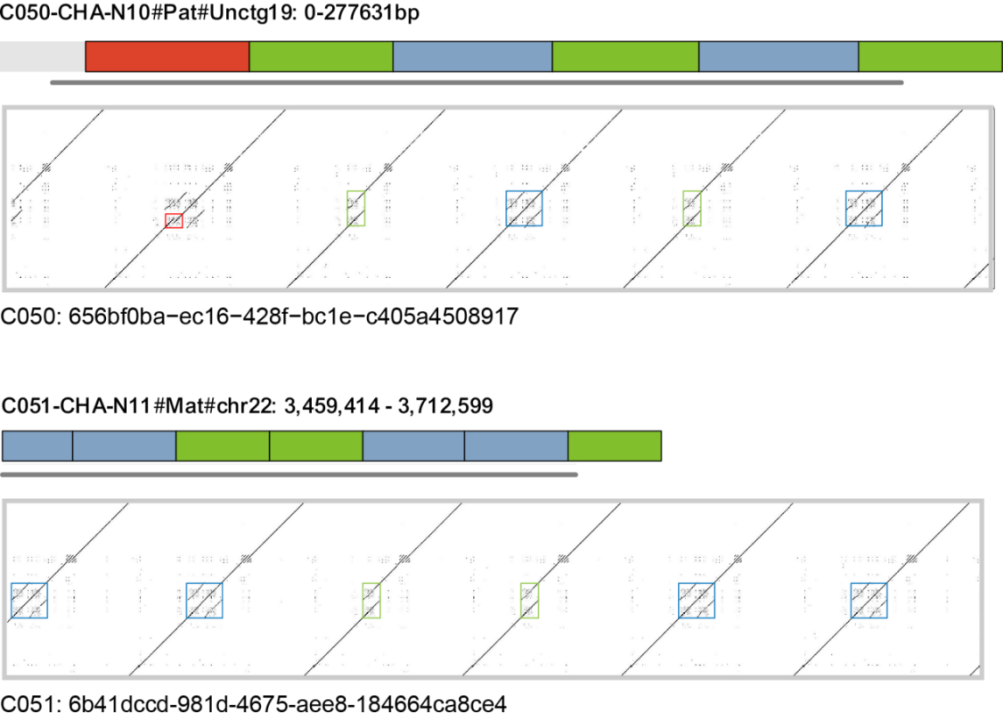

**Supplementary Fig. 13 | Examples of rDNA haplotyping comparison between ONT long reads and genome assemblies.** In the nucleotide dot plot, the vertical axis represents the reference sequence of the human rDNA element (KY962518) and the horizontal axis represents the full length of a single ONT read. The colors denote different rDNA haplotypes.

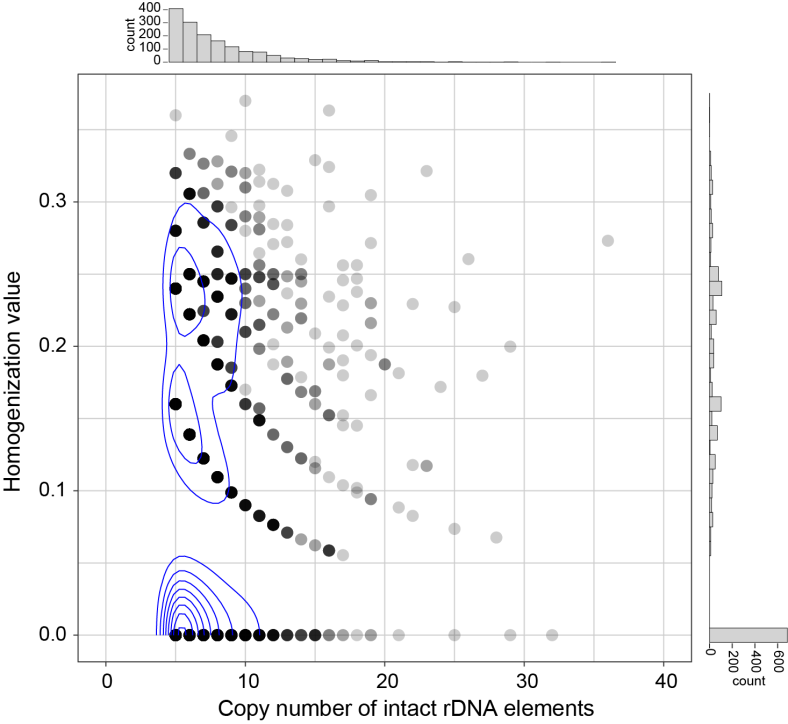

**Supplementary Fig. 14 | Homogenization pattern of human rDNA arrays.** A total of 1,569 large rDNA arrays (with at least five intact rDNA copies) are included for analysis. Homogenization values, reflecting haplotype diversity within each intact rDNA array, are defined as the probability that any two rDNA units originate from distinct haplotypes; a homogenization value of 0 indicates that all repeats within the array originate from a single uniform haplotype.

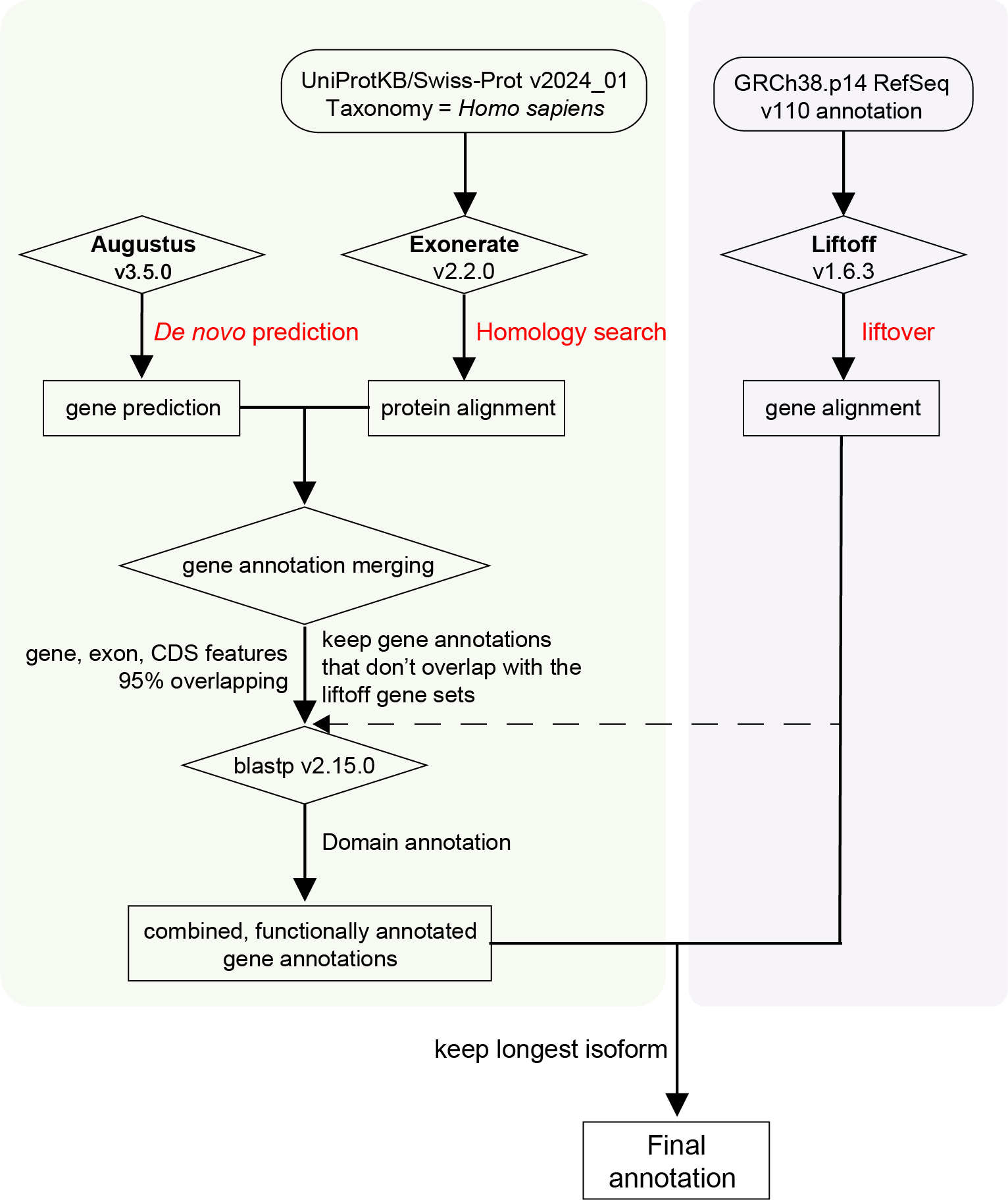

**Supplementary Fig. 15 | Workflow for gene annotation.** Three annotation approaches are recruited, including liftover from GRCh38.p14 annotation, homology search and *de novo* prediction.

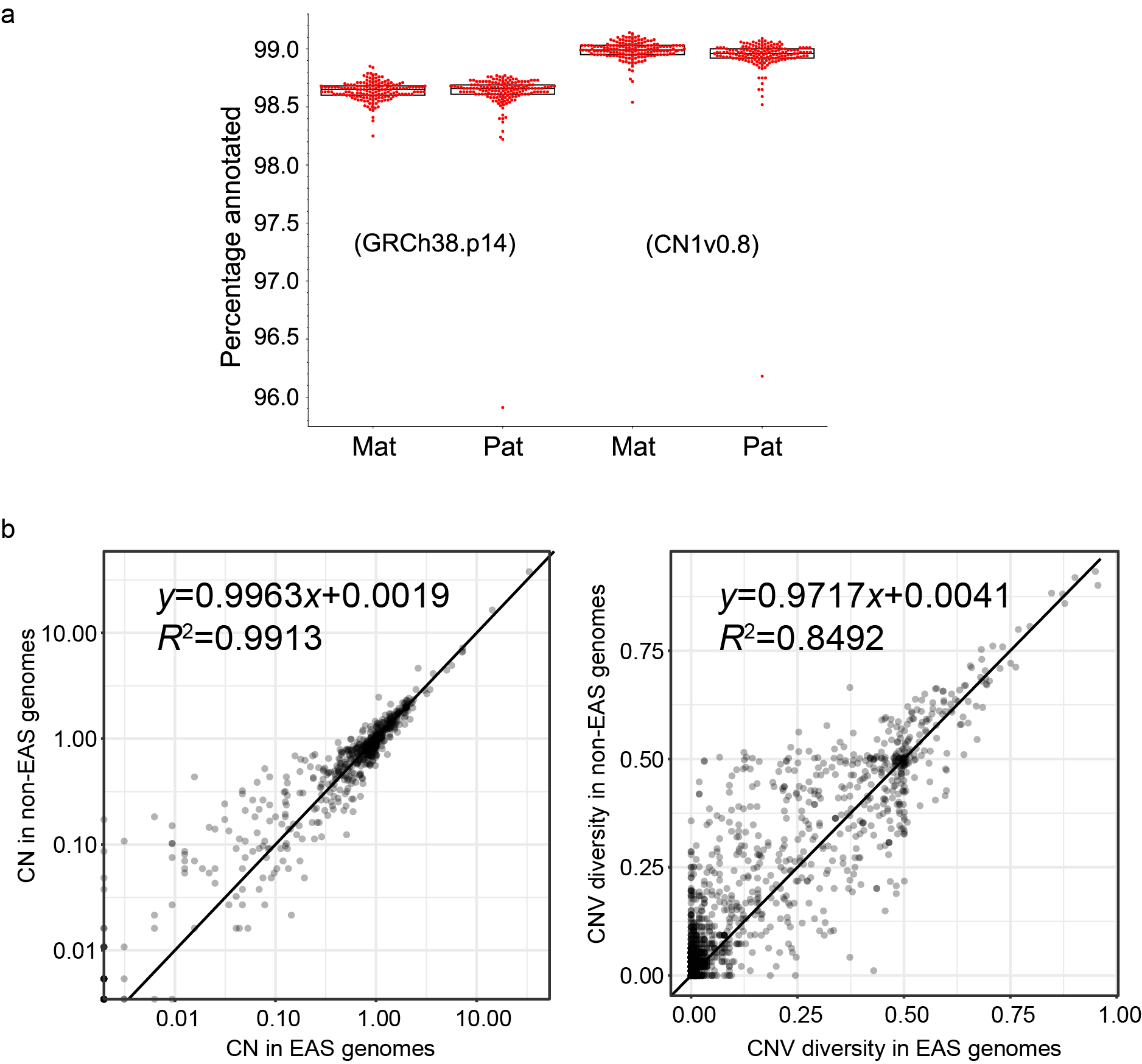

**Supplementary Fig. 16 | Gene annotation of APGp1 phased assemblies. a**, Percentages of protein-coding genes annotated from the reference sets (GRCh38p14 and T2T-CN1 v0.8) for APGp1 maternal and paternal assemblies. **b**, Comparison of gene copy number (CN) and copy number variation (CNV) diversity between EAS samples from APGp1, and EAS genomes from HPRCy1 and HGSVC3.

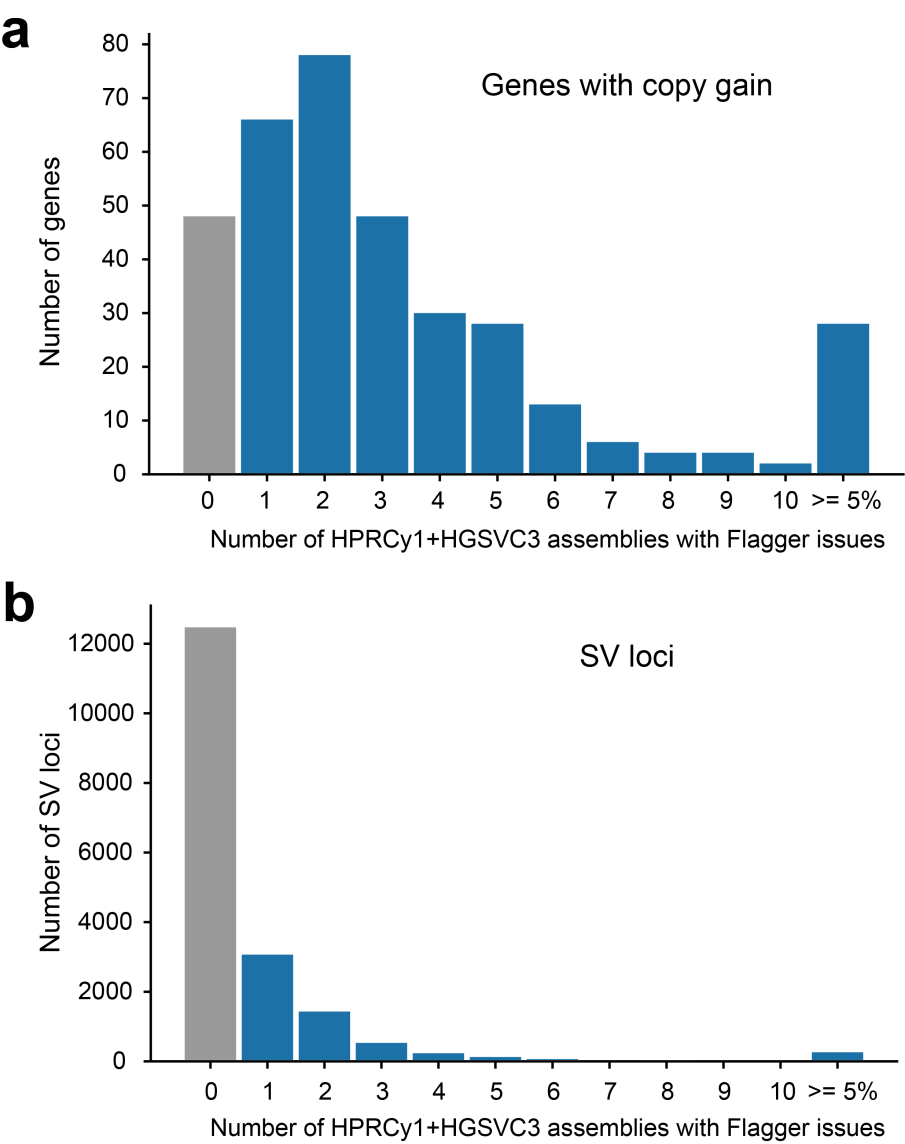

**Supplementary Fig. 17 | Distribution of Flagger issues across APGp1-specific genes with copy gain (a) and SV loci (b) in HPRCy1+HGSVC3 assemblies.** The x-axis represents the number of non-APGp1 assemblies in which Flagger issues were detected for each genomic feature. Integer bins (0-10) show the exact count of affected assemblies, while the rightmost bin (≥5%) aggregates all features where ≥5% of non-APGp1 assemblies contain Flagger issues.

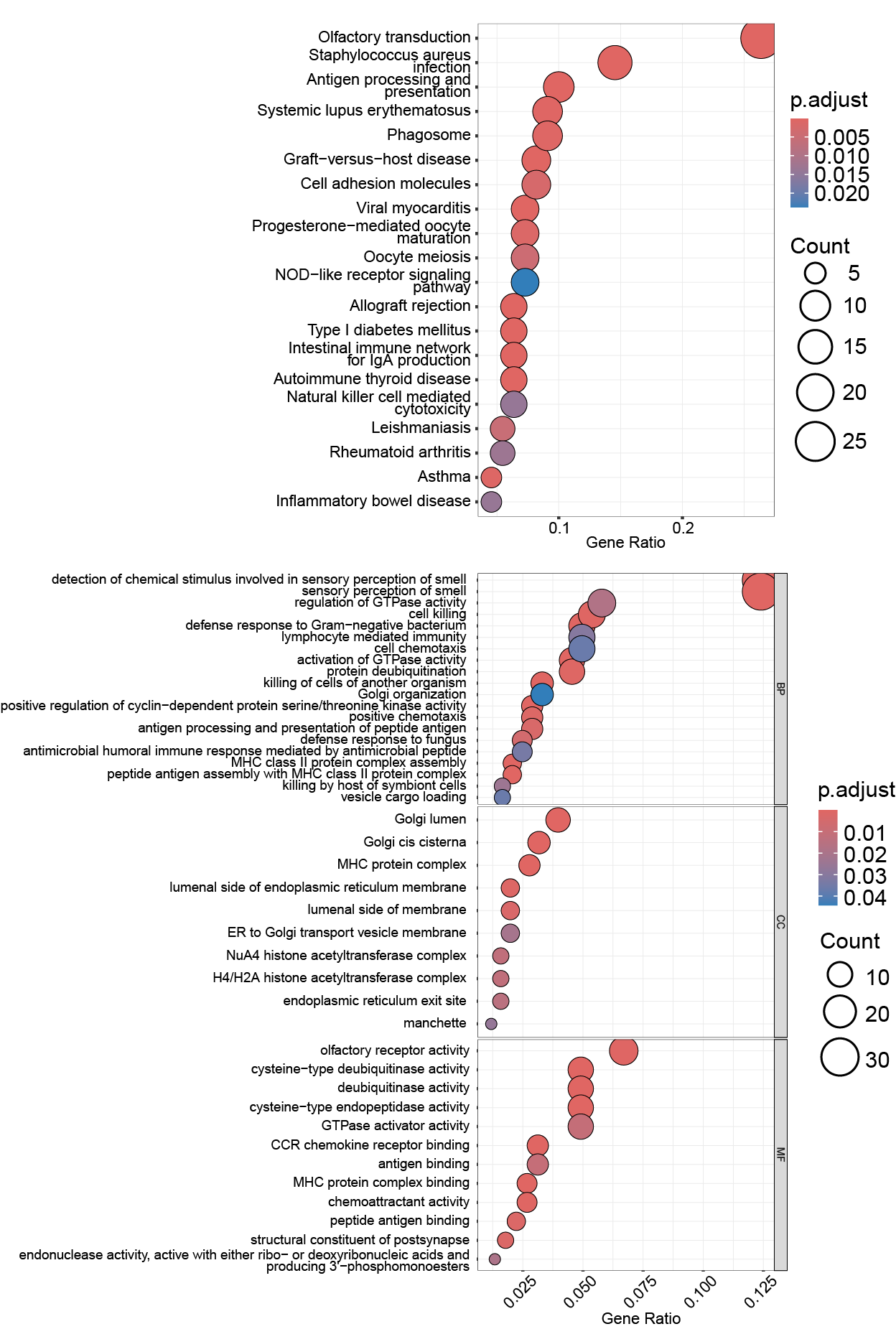

Supplementary Fig. 18 | Functional enrichment analysis of genes with high copy number variation (**CNV) diversity in APGp1 assemblies.** Genes with CNV diversity value greater than 0.26 (top 2% threshold , *n* = 384 ) are included in the analysis. Upper panel: Kyoto Encyclopedia of Genes and Genomes (KEGG) pathway. Lower panel: Gene Ontology (GO) enrichment across biological processes, molecular functions, and cellular components.

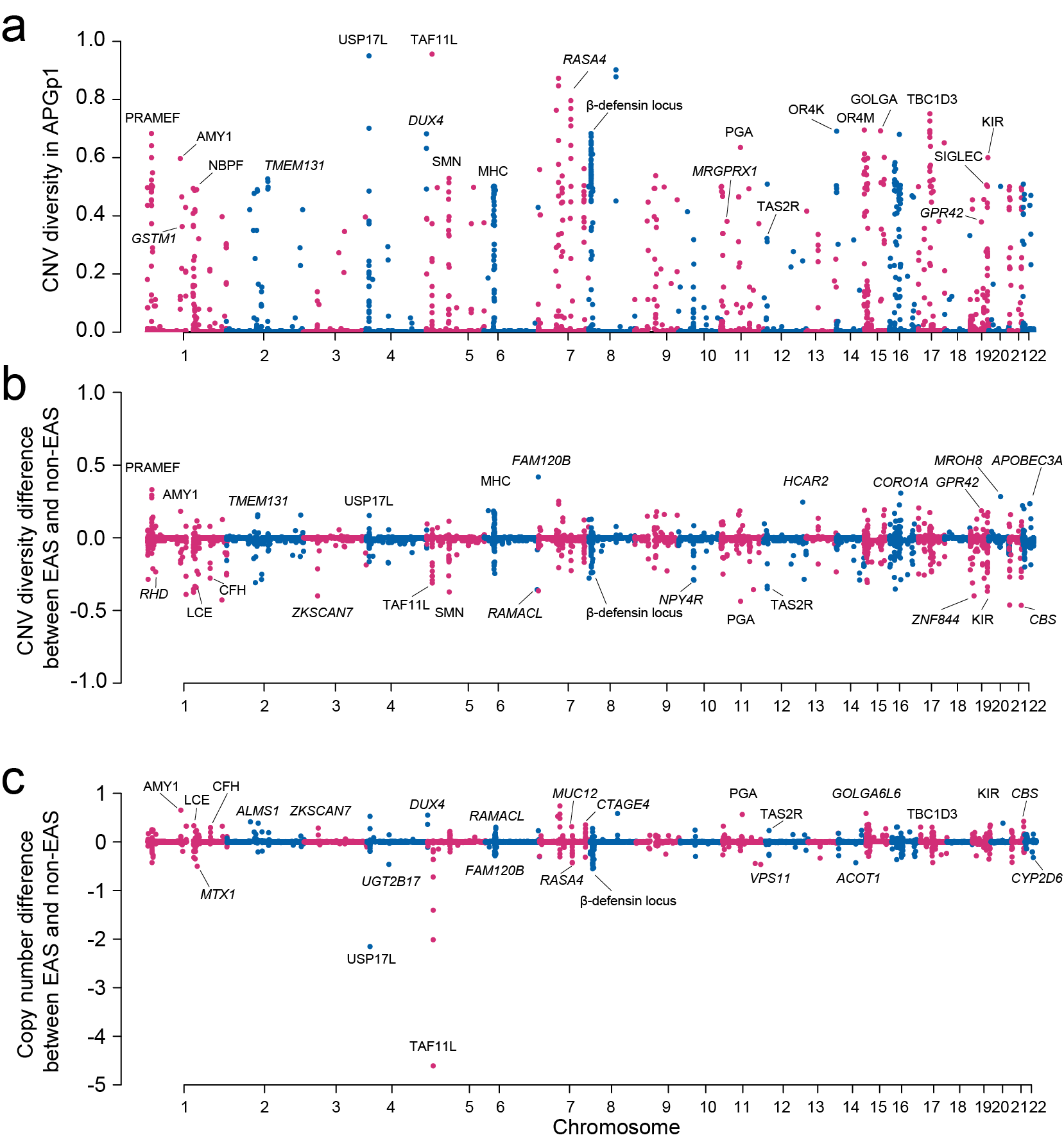

**Supplementary Fig. 19 | CNV diversity and differentiation in human populations.** **a**, CNV diversity along 22 human autosomal chromosomes in APGp1 genomes. **b**, Difference in CNV diversity between EAS and non-EAS human genomes, where positive values indicate higher CNV diversity in EAS. **c**, Absolute average copy number difference between EAS and non-EAS genomes, with positive values denoting higher average gene copy number in EAS.

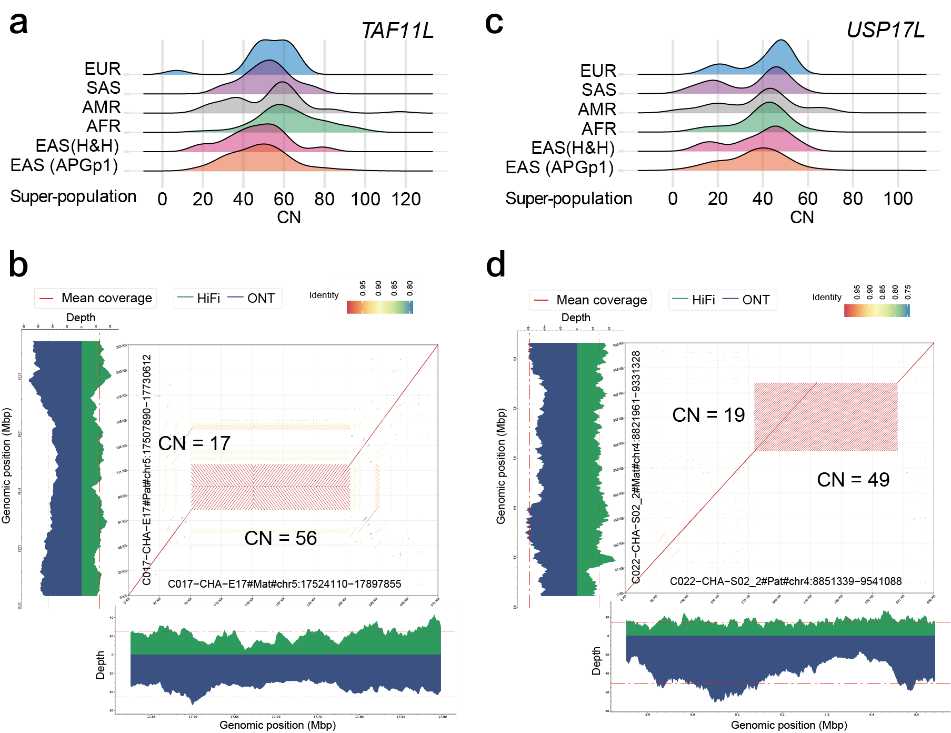

**Supplementary Fig. 20 | CNV in the *TAF11L* and *USP17L* gene clusters.** **a**, *TAF11L* gene copy number distribution among super populations. **b**, Assembly validation and comparison of the *TAL11L* cluster with different copy numbers between assemblies C017-CHA-E17-01#Mat (CN = 56) and C017-CHA-E17-01#Pat (CN = 17). Mapping coverages are calculated from curated alignments in GCI. **c**, *USP17L* gene copy number distribution. **d**, Assembly validation and comparison of the *USP17L* cluster between assemblies C022-CHA-S02_02-01#Pat (CN = 49) and C022-CHA-S02_02-01#Mat (CN = 19).

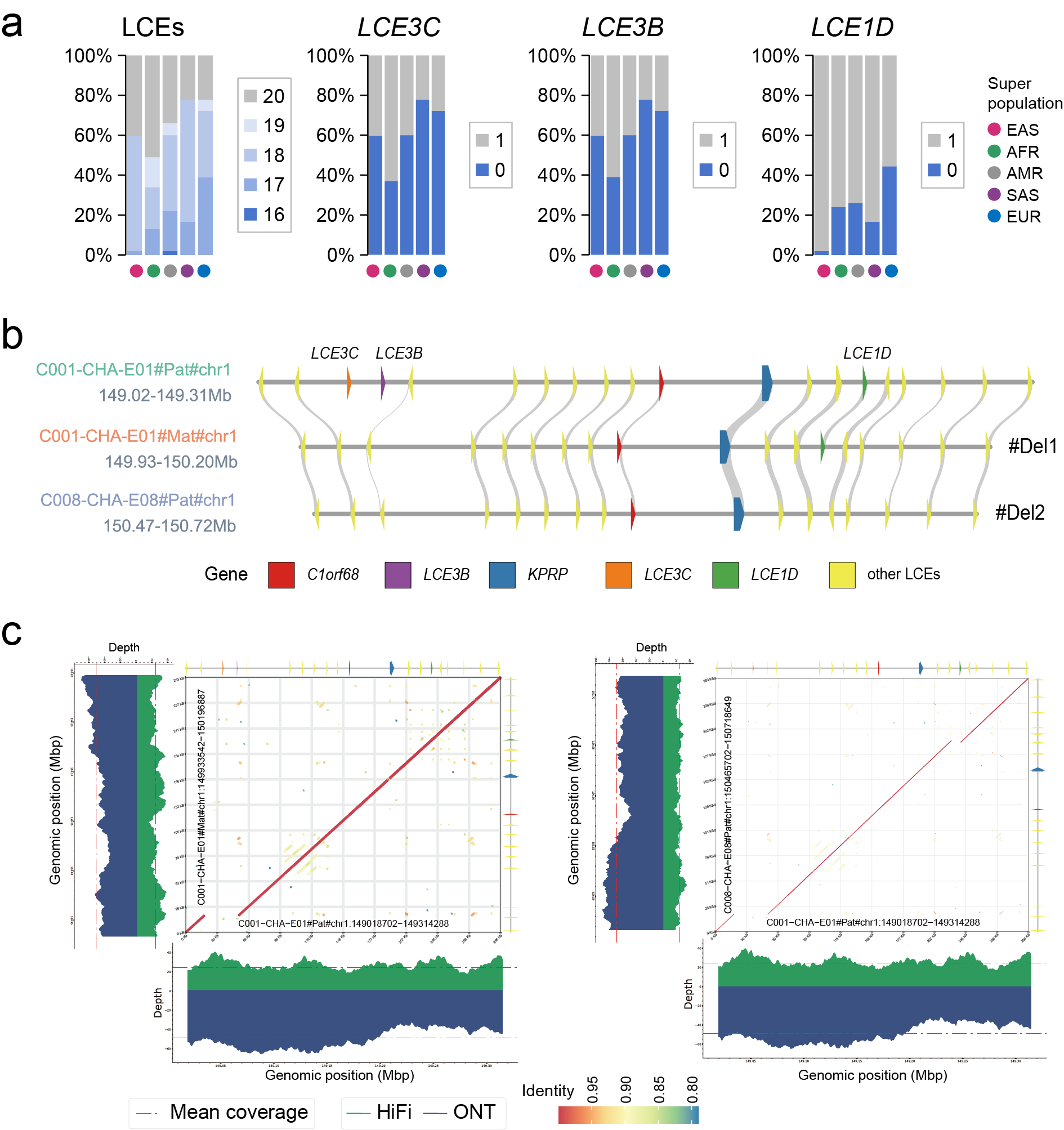

**Supplementary Fig. 21** | **CNV in the LCE gene cluster**. **a**, CNV frequency of human Late Cornified Envelope (LCE) genes across five super populations. **b**, Schematic diagram of CNV in the LCE gene cluster. **c**, Assembly validation and comparison among LCE clusters in assemblies C001-CHA-E01-01#Pat, C001-CHA-E01-01#Mat and C008-CHA-E08-01#Pat, representing three alleles.

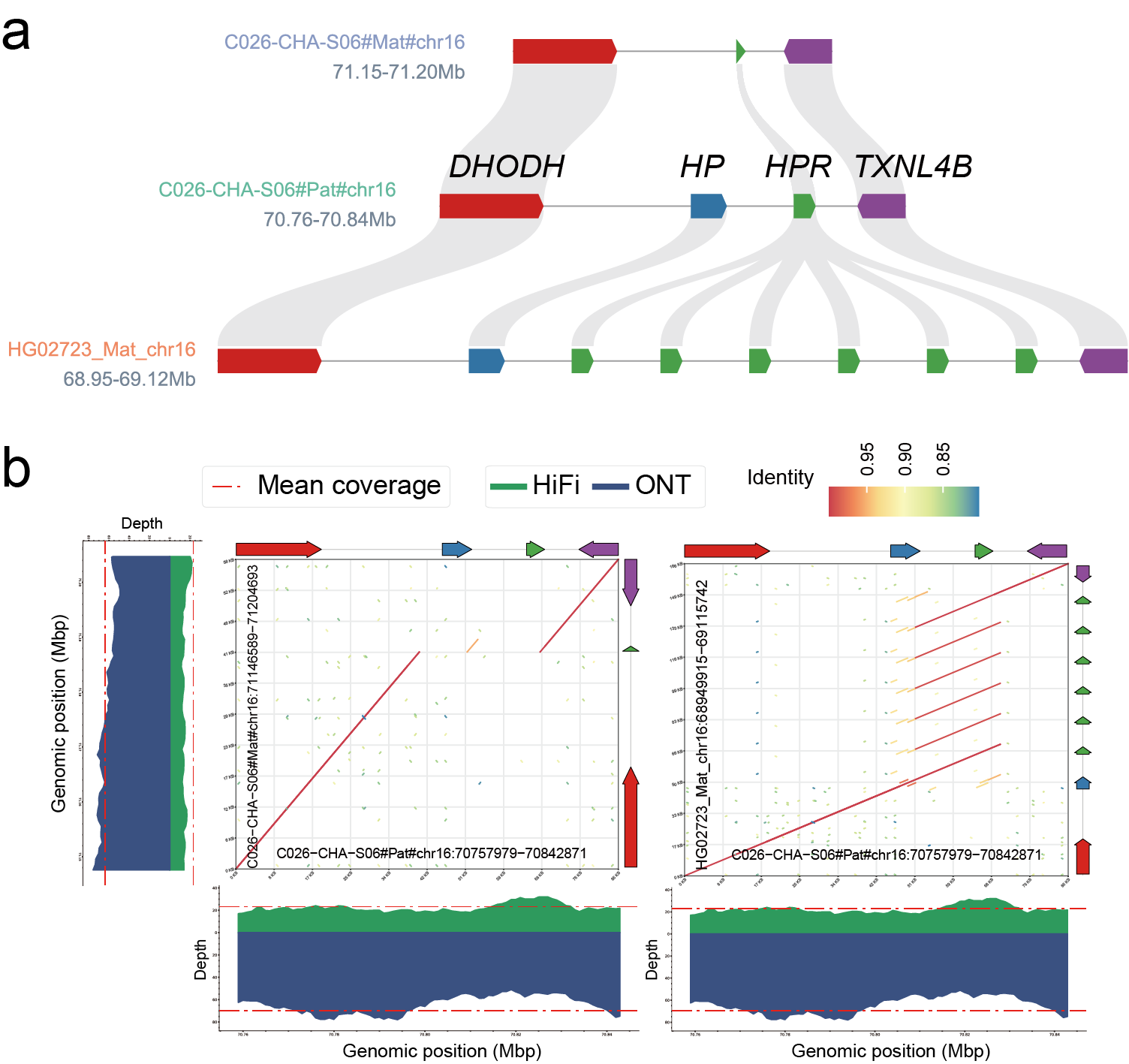

**Supplementary Fig. 22 | CNV in *HP* and *HPR* genes. a**, Schematic diagram of CNVs in genes *HP* and *HPR*. **b**, Assembly validation and comparison in the *HP* and *HPR* gene region among assemblies C026-CHA-S06-01#Mat, C026-CHA-S06-01#Pat, and HG02723#Mat.

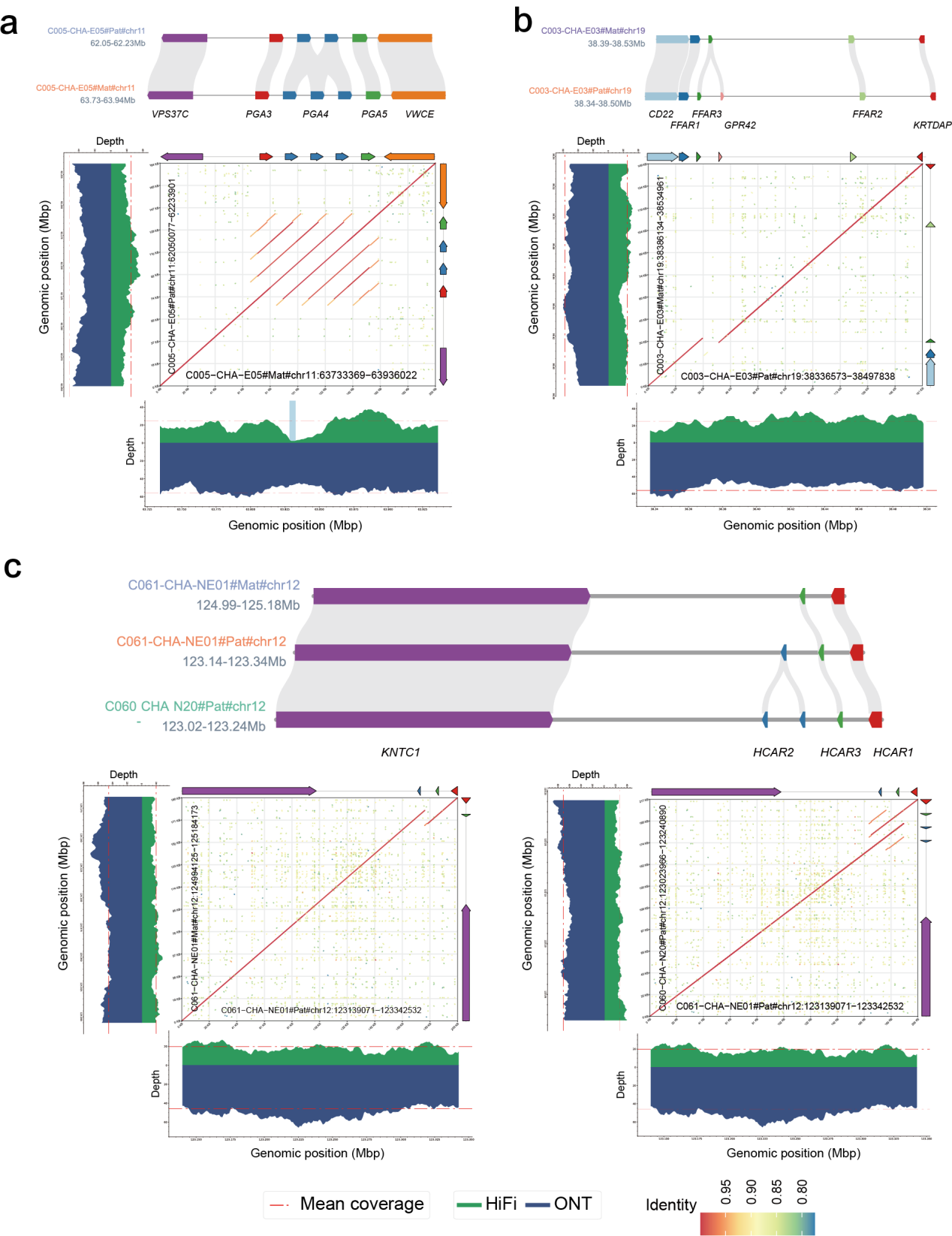

**Supplementary Fig. 23 | CNV in diet-related genes. a**, CNV and assembly validation of gene *PGA4*. **b**, Gene *GPR42*. **c**, Gene *HCAR2*.

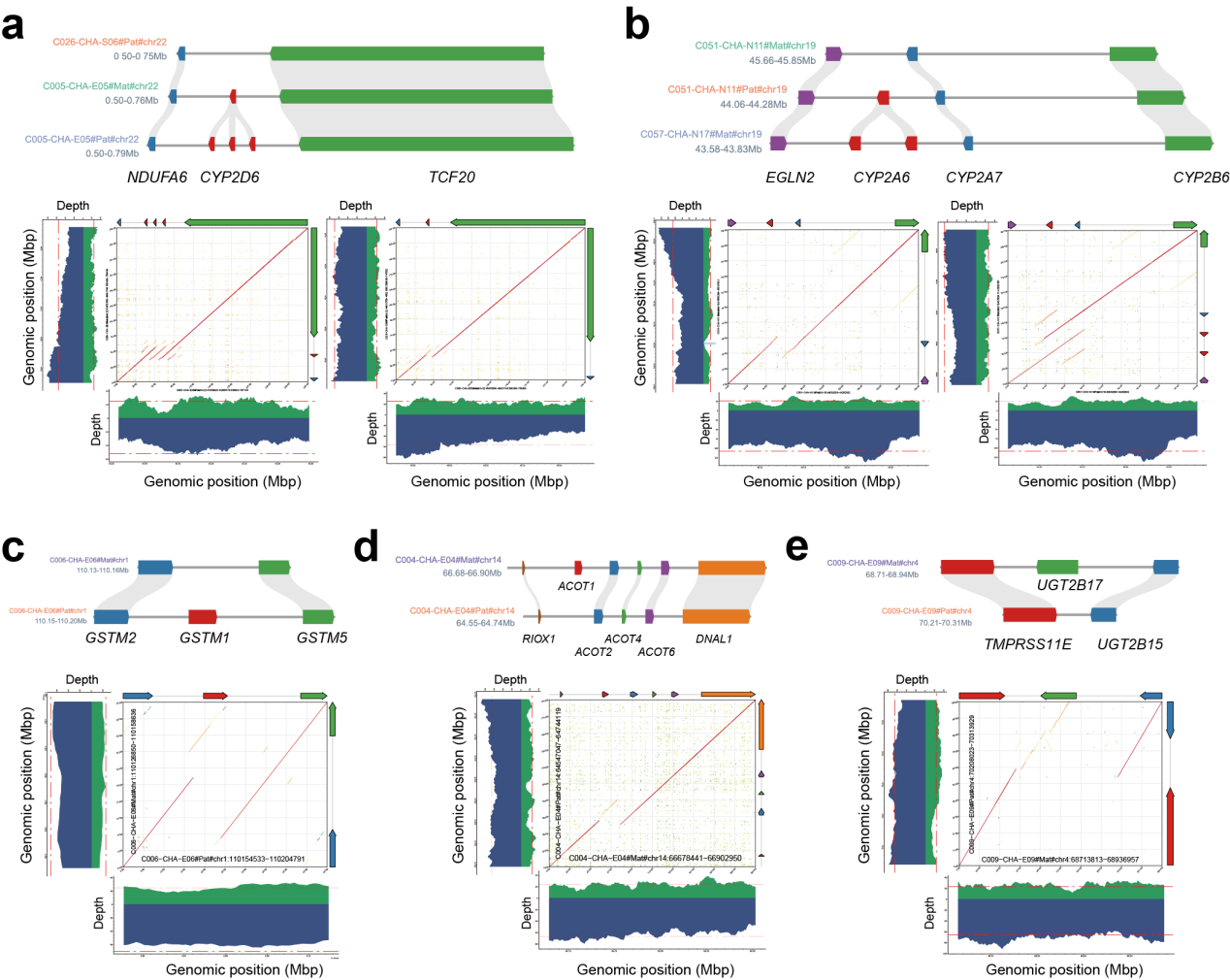

**Supplementary Fig. 24 | CNV in drug-metabolism-related genes. a**, Gene *CYP2D6*. **b**, Gene *CYP2A6*. **c**, Gene *GSTM1*. **d**, Gene *ACOT1*. **e**, Gene *UGT2B17*.

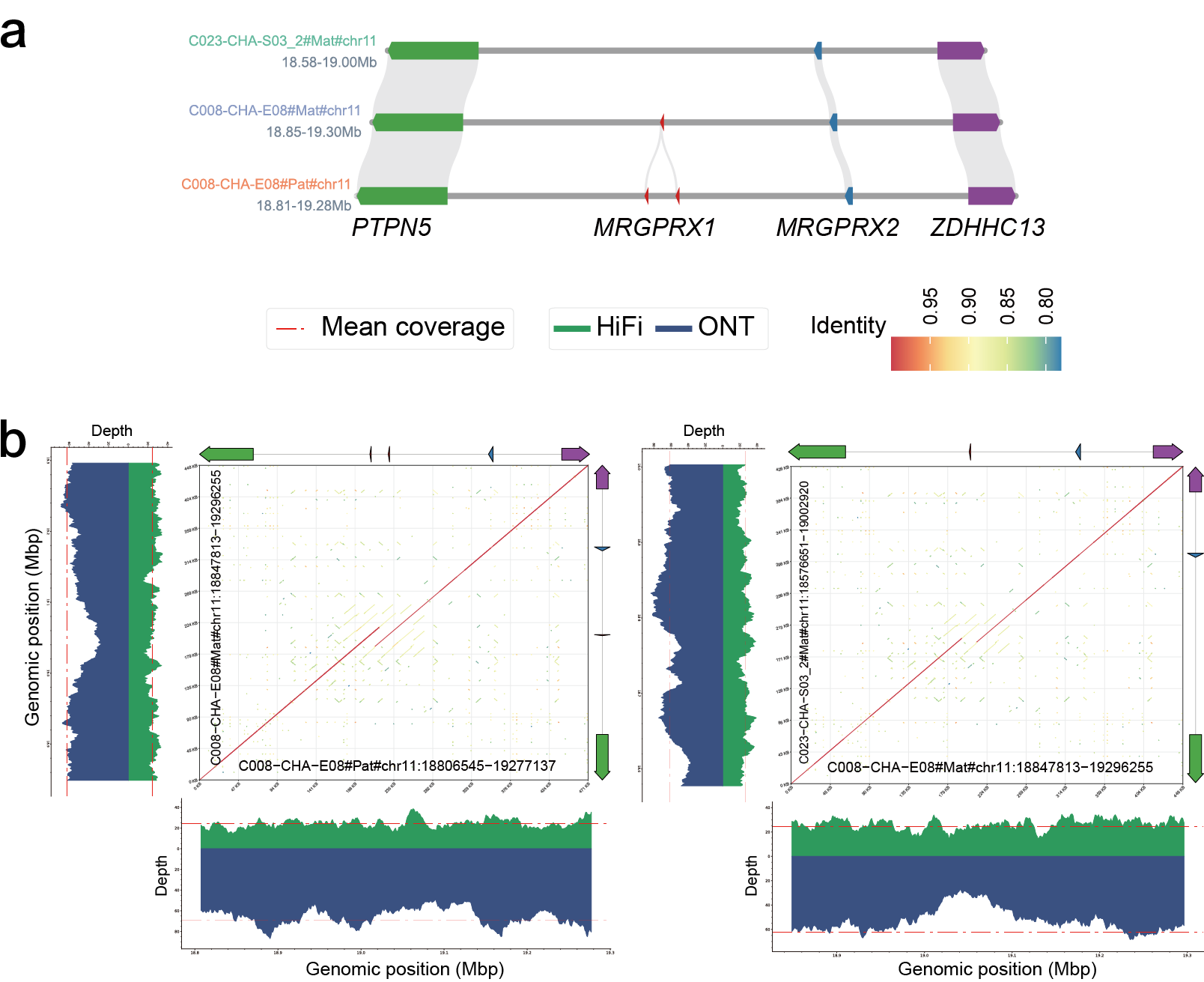

**Supplementary Fig. 25** | **CNV in gene *MRGPRX1*. a**, Schematic diagram of CNVs for the gene *MRGPRX1*. **b**, Assembly validation and comparison of gene *MRGPRX1* among assemblies C008-CHA-E08-01#Mat, C023-CHA-S03_2-01#Pat, and C008-CHA-E08-01#Pat.

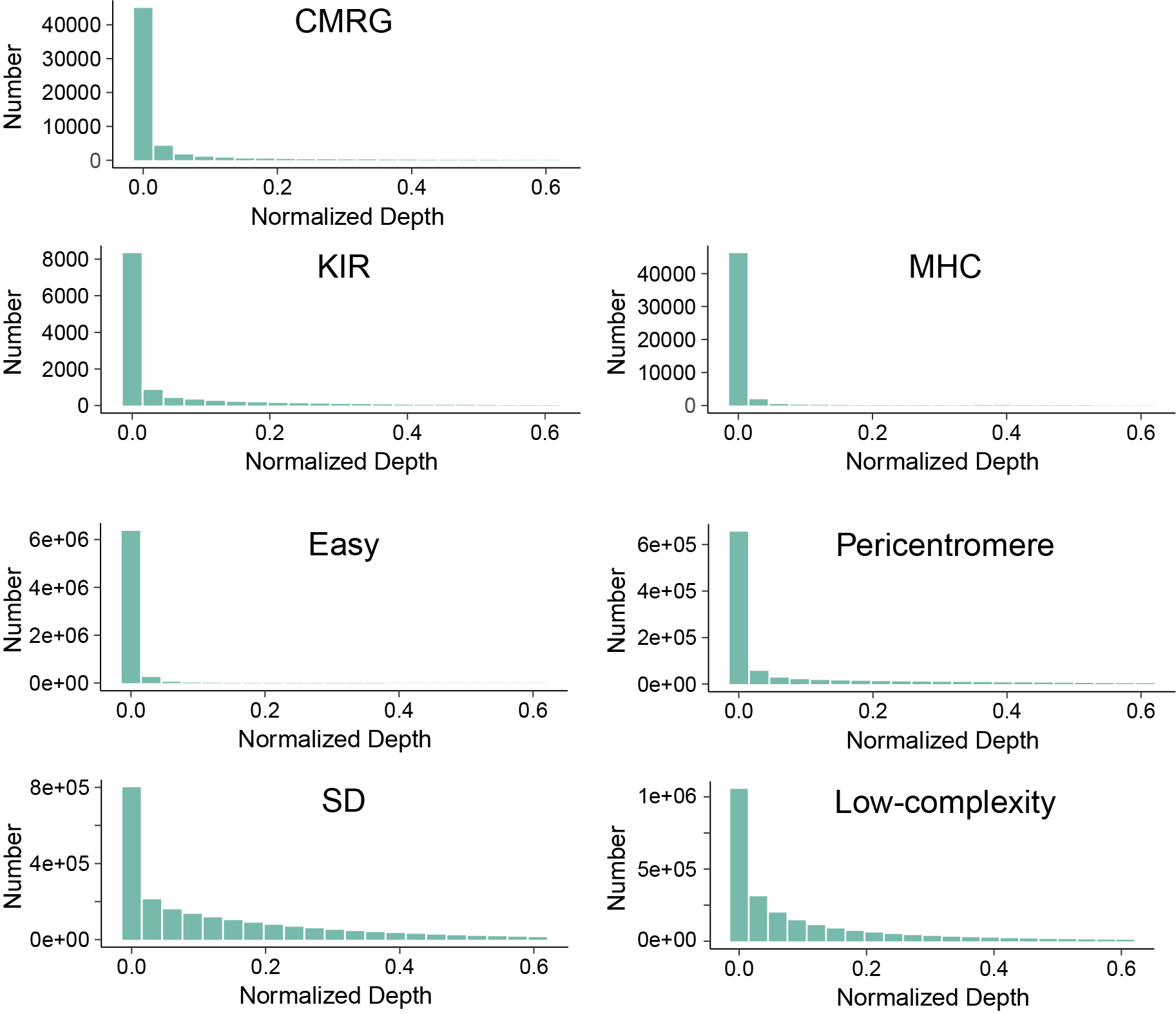

**Supplementary Fig. 26 | Mapping depth of NGS short reads on off-target edges in the APGp1 T2T-CN1-referenced MC pangenome graph.** Off-target alignments represent the reads failing to target the haplotypic path through the graph. CMRG, Challenging Medically Relevant Gene.

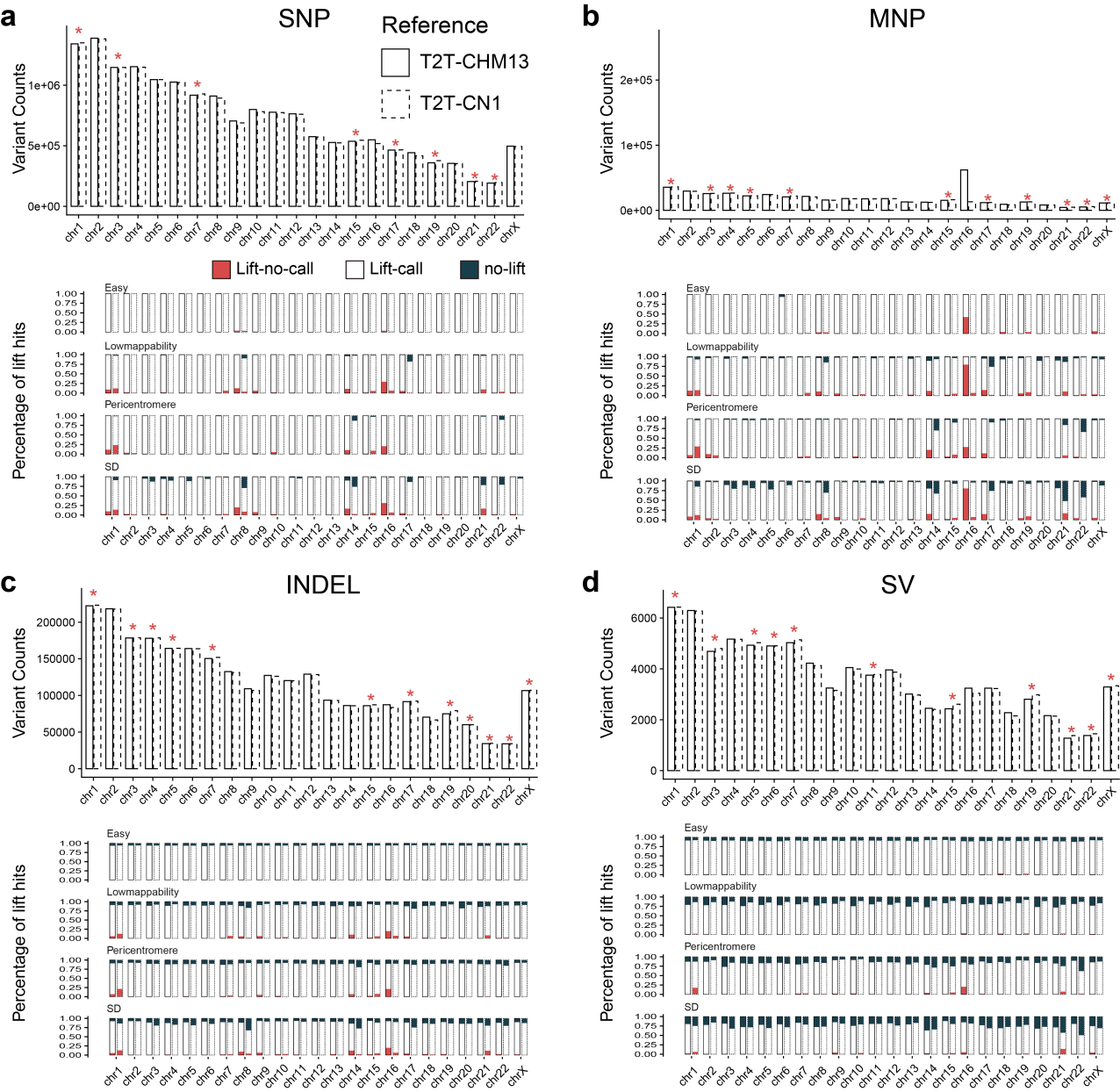

**Supplementary Fig. 27 | Comparative assessment of variant representation and coordinate lift efficiency between T2T-CHM13 and T2T-CN1-referenced APGp1 graphs. a-d,** Comparison of variant statistics and coordinate lifting across four variant classes: SNPs (**a**), MNPs (**b**), InDels (**c**), and SVs (**d**). For each panel, the upper plot illustrates the total variant count per chromosome. Solid bars represent the APGp1 graphs using T2T-CHM13 as backbone, while dashed-outline bars represent the T2T-CN1-based graph. Red asterisks denote chromosomes where the variant count in the T2T-CN1-referenced graph exceeds that of the T2T-CHM13-referenced graph. The lower plot provides a percentage-based breakdown of pairwise lifting results across stratified genomic regions of varying complexity, including ‘Easy’, ‘Lowmappability’, ‘Pericentromere’ and ‘SD’ (detailed definitions in **Methods**). Lifting outcomes are categorized into three classes: ‘no-lift’ (no alignable coordinates on the alternative reference), ‘lift-no-call’ (successful coordinate lift-over but no overlapping variant call), and ‘lift-call’ (successful lift-over with a corresponding overlapping variants).

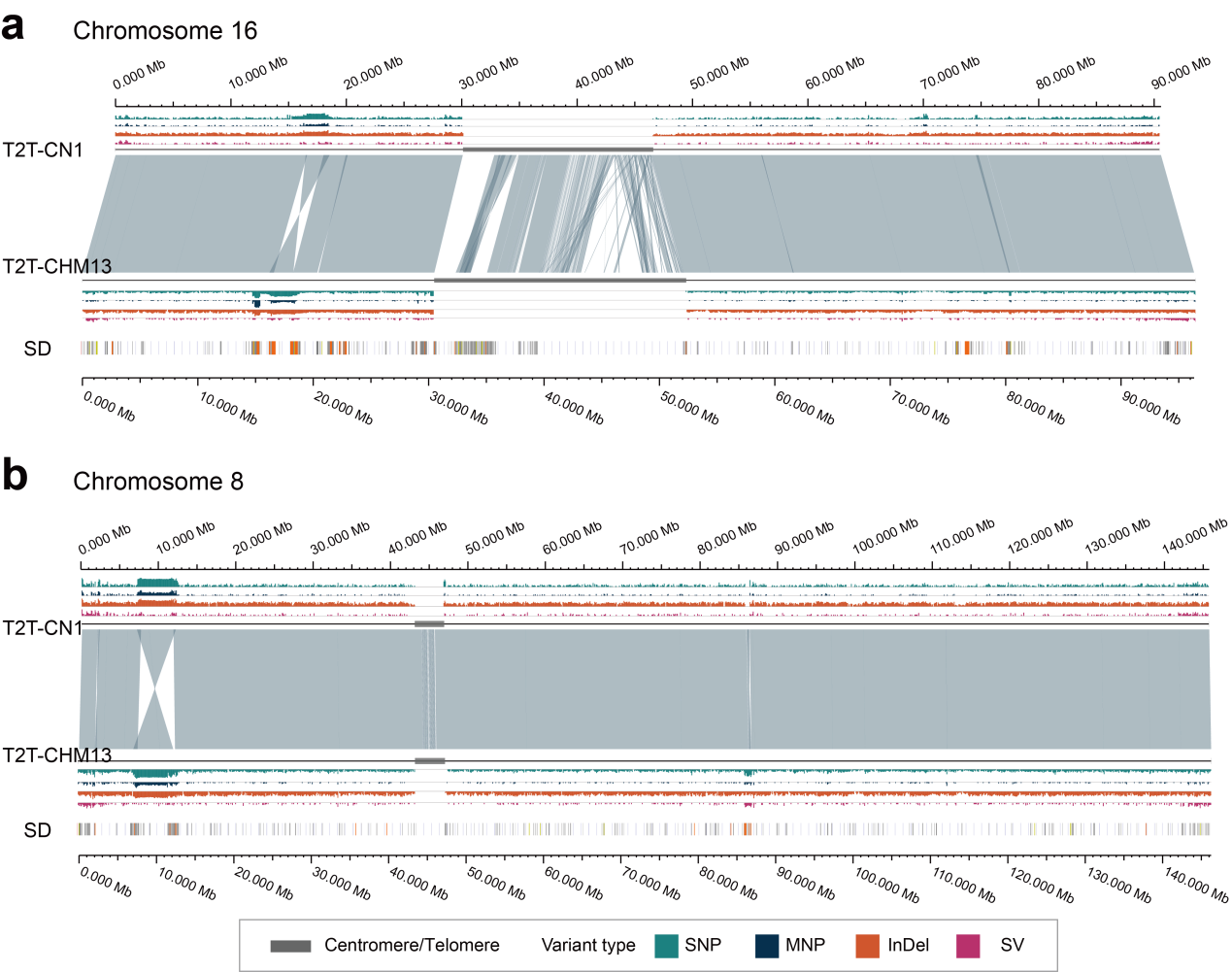

**Supplementary Fig. 28 | Landscape of variant lift failures between T2T-CN1- and T2T-CHM13-referenced pangenome graphs on chromosomes 16 and 8.** The middle tracks display sequence alignments and synteny between T2T-CN1 and T2T-CHM13. Lift failures of variant calls per 100-kbp window are shown for SNPs, MNPs, InDels and SVs, respectively. Centromere regions are masked. Lift failures include two categories: ‘lift-no-call’ (variants that can be lifted but fail to overlap with the target graph's variants) and ‘no-lift’ (variants with no alignable coordinates in the target graph). The segmental duplication (SD) annotations on the T2T-CHM13 are provided as the bottom track.

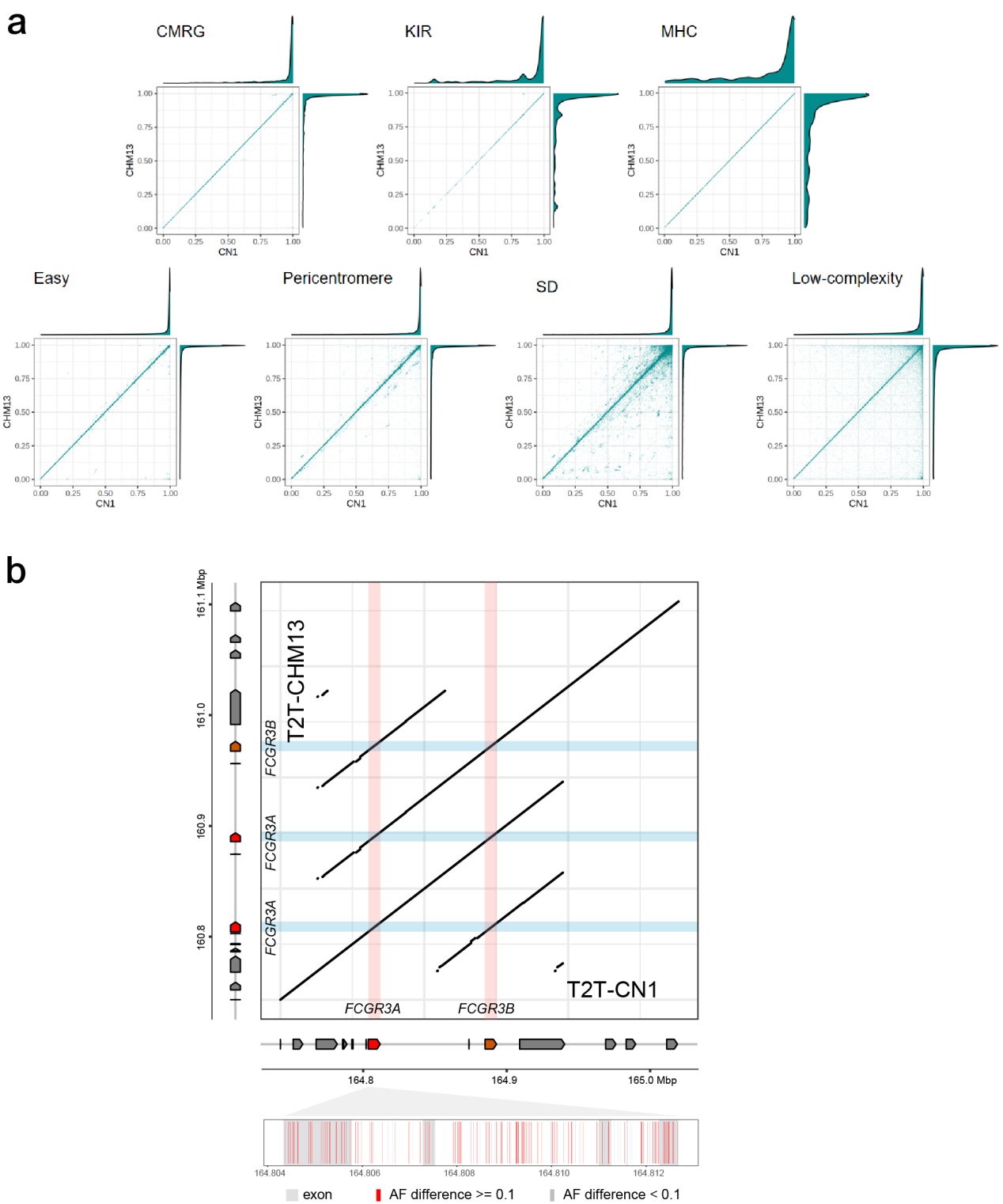

**Supplementary Fig. 29 | Reference bias in allele frequency of variants decoded from MC graphs with different backbones.** **a**, Allele frequency differences for variants decomposed from T2T-CN1 and T2T-CHM13-referenced graphs across different genomic regions. CMRG, Challenging Medically Relevant Gene. **b**, Allele frequency differences of variants in the *FCGR3* gene cluster, with observed gene copy number variation between T2T-CHM13 and T2T-CN1.

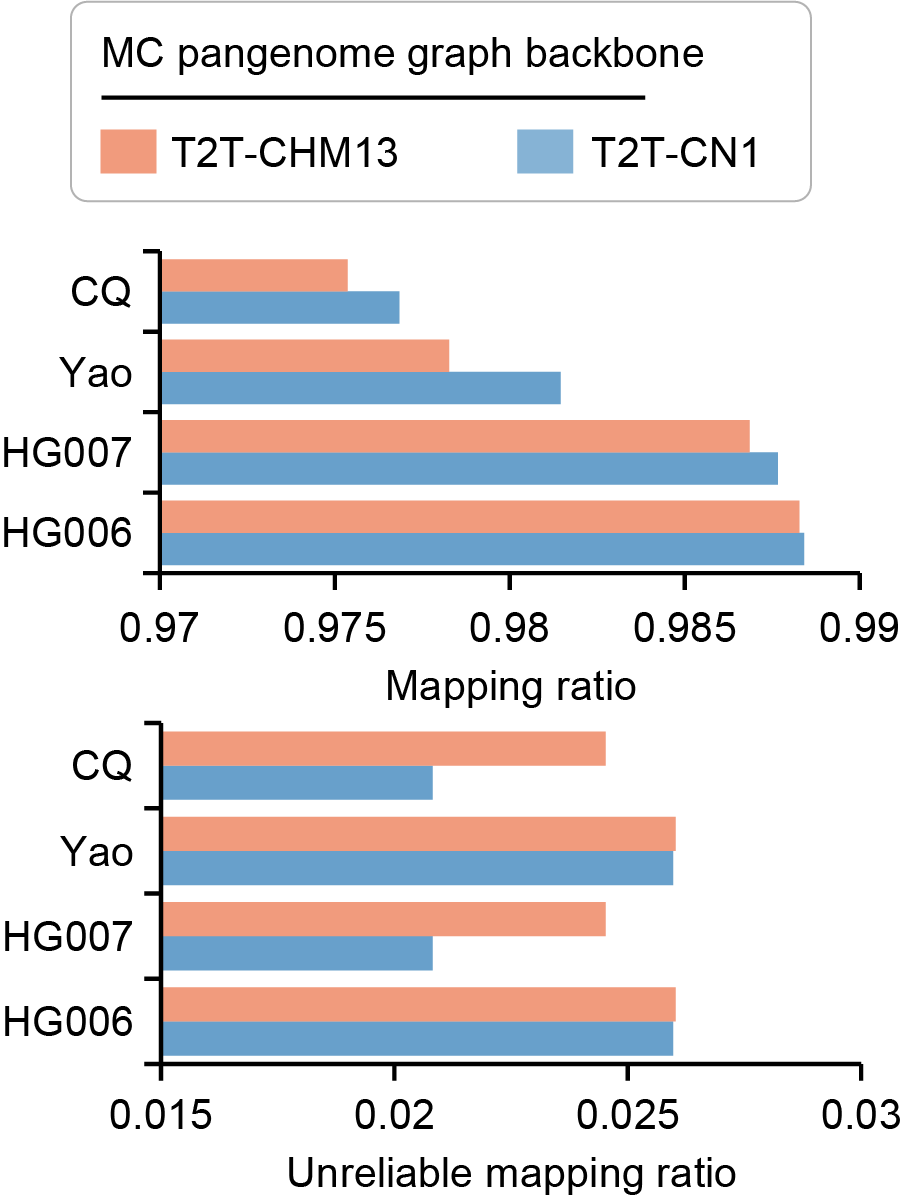

**Supplementary Fig. 30 | Mapping performance of short and long reads from East Asian (EAS) genomes on T2T-CN1 and T2T-CHM13-based MC graphs.** NGS reads of HG006 and HG007 from GIAB are aligned against graphs using vg Giraffe (https://github.com/vgteam/vg), and PacBio HiFi reads of Chinese Quartet (CQ) and Yao samples are aligned using GraphAligner (https://github.com/maickrau/GraphAligner). Upper panel: mapping ratio; Lower panel: unreliable mapping ratio, defined as the proportion of bases with abnormal mapping coverage (i.e. < 0.1 or > 5 times the whole-genome average depth), across the whole pangenome graph.

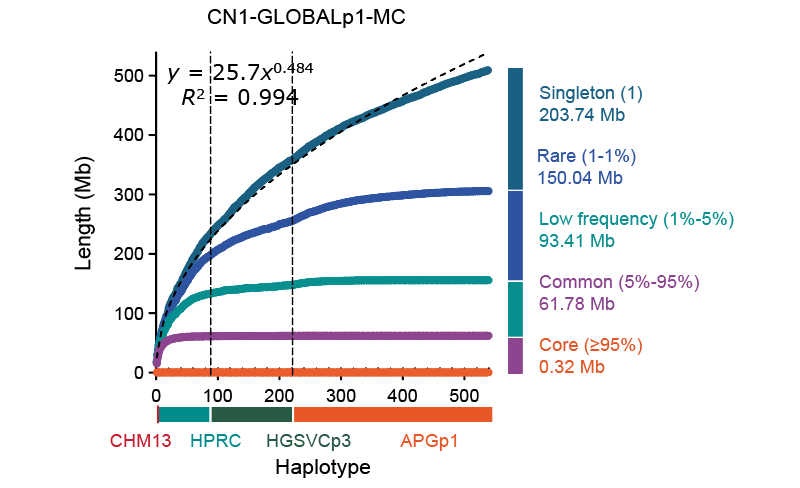

**Supplementary Fig. 31 | Growth curve of the human pangenome graph CN1.GLOBALp1.MC.** This graph is constructed using Minigraph-Cactus (MC) by integrating global human genome assemblies from APGp1, HPRCy1, and HGSVC3, and T2T-CHM13, with T2T-CN1 set as the reference backbone. The dashed line represents a fitting curve, with its formula and correlation displayed. Four categories of genomic sequences in the final graph are categorized: singleton, rare, low-frequency and core sequences, defined by the occurrence frequency across 540 human genome assemblies.

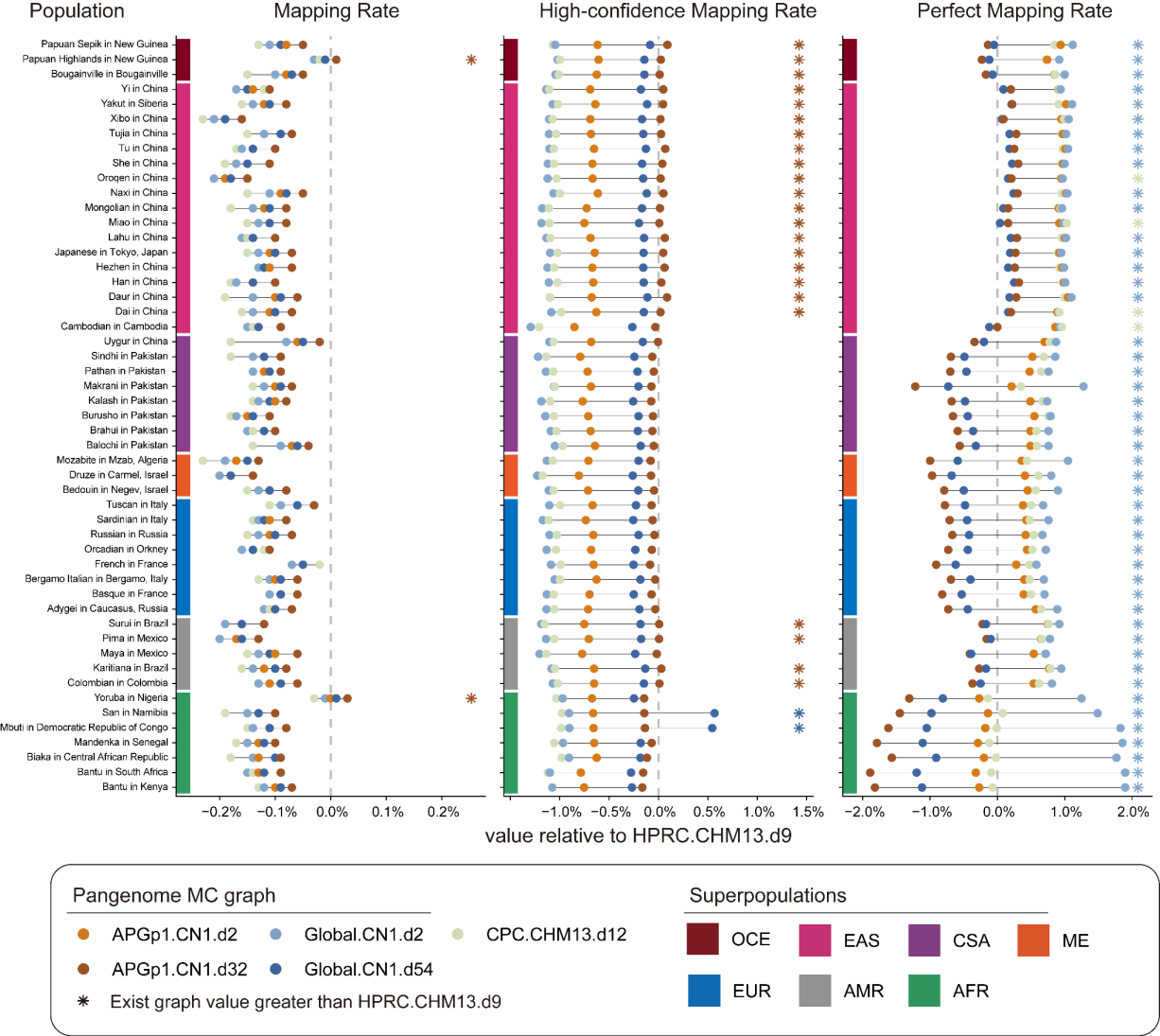

**Supplementary Fig. 32 | Read mapping performance across human ancestries using multiple human pangenome graphs.** A total of 51 human populations from AFR (Africa), EUR (Europe), ME (Middle East), SAS (South Asia), EAS (East Asia), OCE (Oceania) and AMR (America) in HGDP are included. As the MC graph IDs show, the six graphs are constructed using diverse human genome assembly collections (HPRCy1, CPC, APGp1 and GLOBALp1). The GLOBALp1 panel integrates assemblies from APGp1, HPRCy1 and HGSVC3. The reference assemblies T2T-CHM13 and T2T-CN1 are set as graph backbones, respectively. Two frequency filtration criteria are implemented: allele counts (AC) ≥ 2 and allele frequency (AF) ≥ 0.1. Specifically, ‘d2’ denotes alleles with at least two counts; ‘d32’ represents variants with AF ≥ 10% across all APGp1 assemblies; ‘d54’ denotes variants with AF ≥ 10% across all APGp1+HPRCy1+HGSVC3 assemblies. The pangenome graphs HPRCy1.CHM13.d9 and CPC.CHM13.d12 were retrieved from previous published studies (Liao *et al*., 2023; Gao *et al*., 2023).

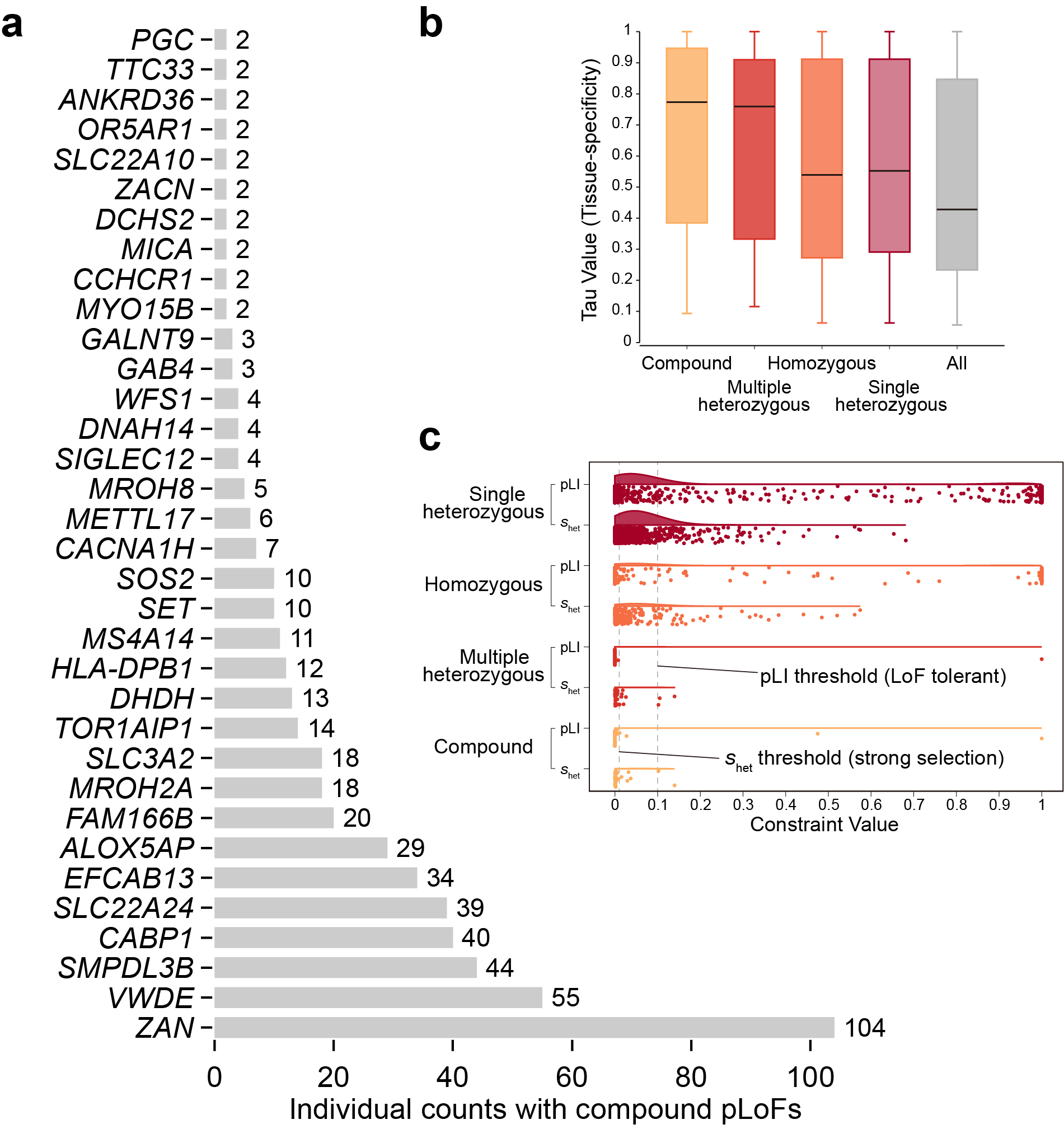

**Supplementary Fig. 33 | Population frequency, tissue specificity and constraint across pLOF genes.** **a**, non-singleton compound pLoF genes in 160 APGp1 individuals. The bar plot displayed the sample size that exhibited compound pLoFs for each gene. **b**, Tissue specificity of genes from different pLOF categories, measured by the tau (*τ*) value. Values approaching 1 indicate tissue-specific expression, whereas values near zero indicate ubiquitous expression. Genes with *τ* ≥ 0.6 are defined as highly tissue-specific. **c**, Constraint scores for genes in the four categories with different pLoF variants in APGp1 samples. Values of pLI (the probability of being LoF intolerant) and *s*_het_ (selective effects for heterozygous protein-truncating variants) close to 1 indicate stronger constraint. Constraint value thresholds for LoF-tolerent genes and genes under strong selection are set at pLI = 0.1 and *s*_het_ = 0.01, respectively.

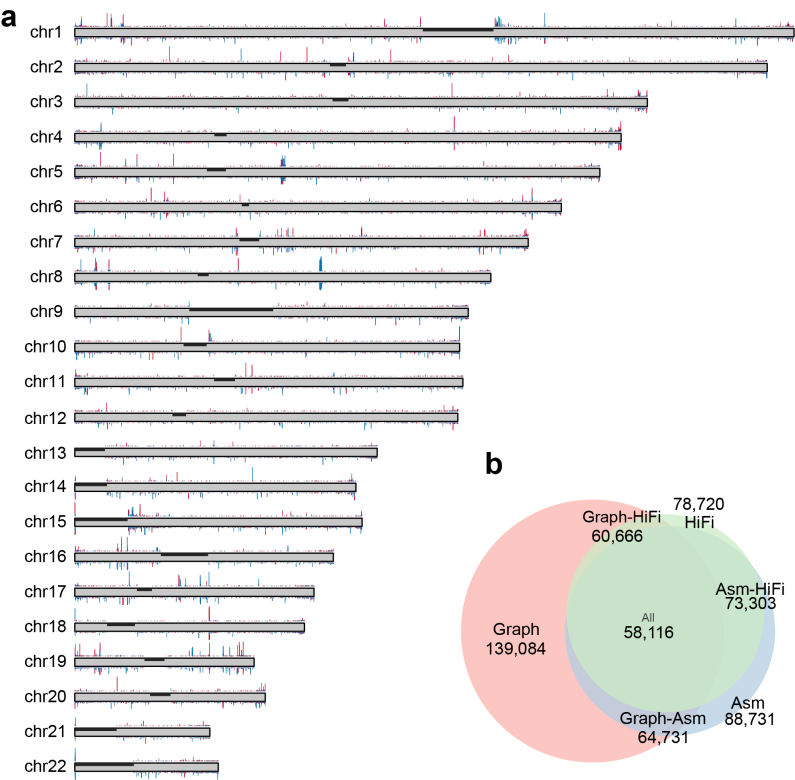

**Supplementary Fig. 34 | Comparison of SV sets derived from pangenome graphs and assemblies. a**, Distribution of SV lengths along genome coordinates for SV sets specific to pangenome graphs (red) and to assemblies (blue). **b**, Venn plot of three SV sets on autosomal chromosomes.

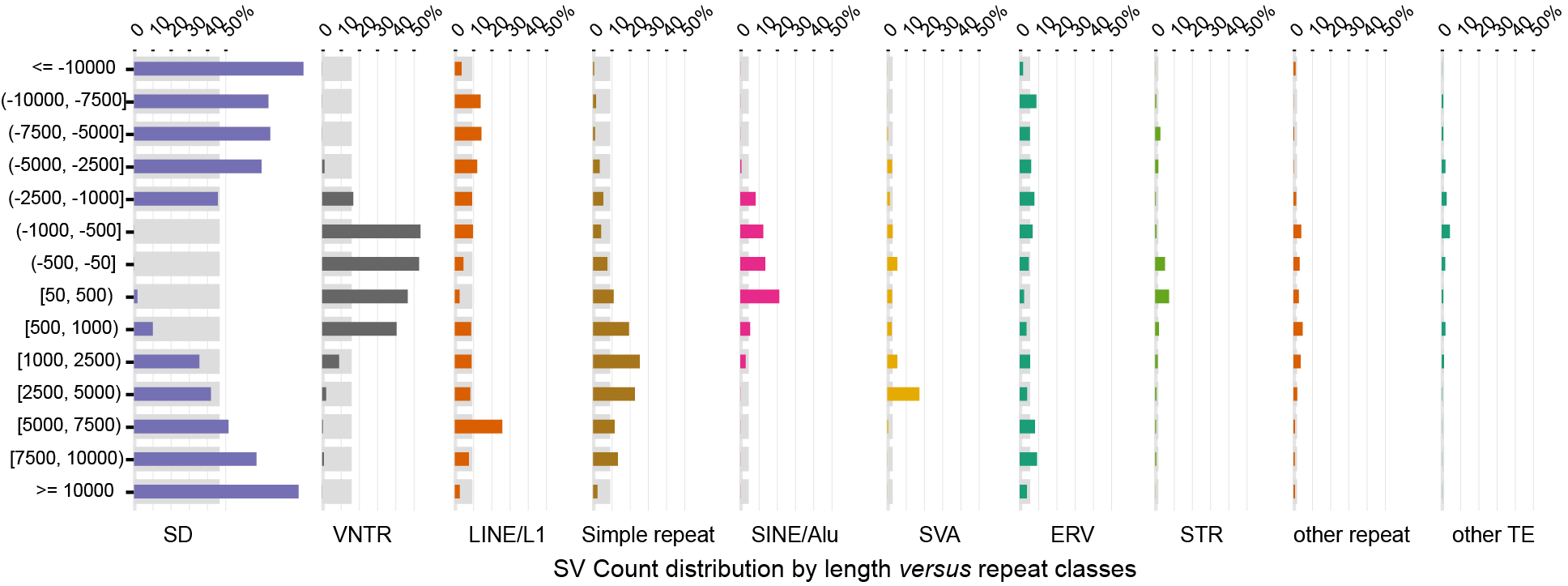

**Supplementary Fig. 35 | Distribution of SV lengths across different repeat classes.** Horizontal bars show the percentage of SVs within each repeat class falling into specified length ranges (colored bars) compared to the average percentage across all repeat classes (grey background bars). SV lengths are categorized by range (*y*-axis) and repeat class (*x*-axis). SD, segmental duplication; VNTR, variable number tandem repeat; STR, short tandem repeat.

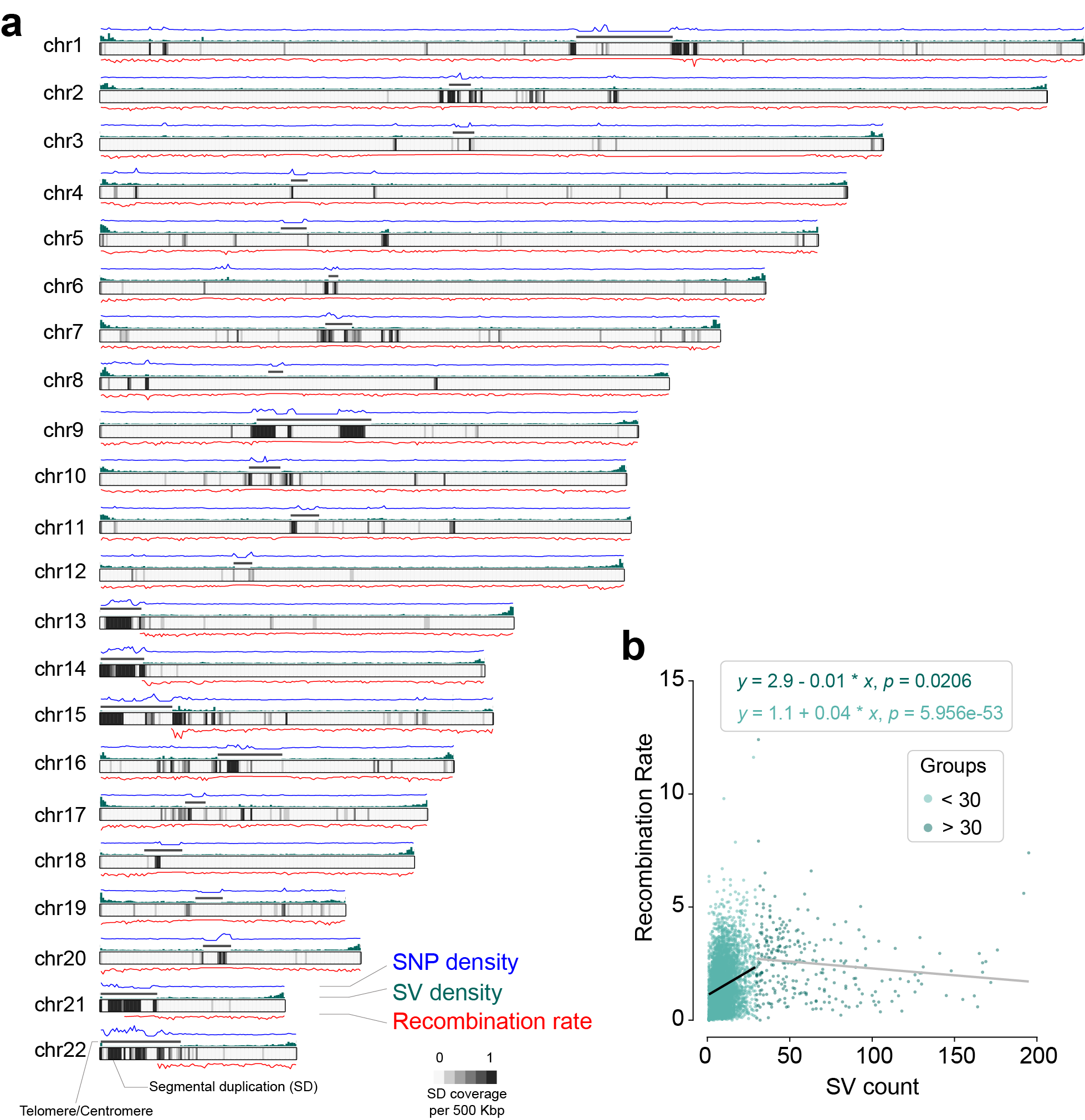

**Supplementary Fig. 36 | Distribution and inter-relationships of SVs, SNPs, recombination and SDs. a**, For each chromosome, the upper panel displays SNP density (blue line) and SV frequency (green histogram) in 500-kbp windows, and the red line below the chromosome ideogram depicts the average recombination rate within corresponding windows. Color-gradient blocks in each chromosome ideogram illustrate the SD coverage in 500-Kbp windows. **b**, Correlation analysis between SV density and recombination rate.

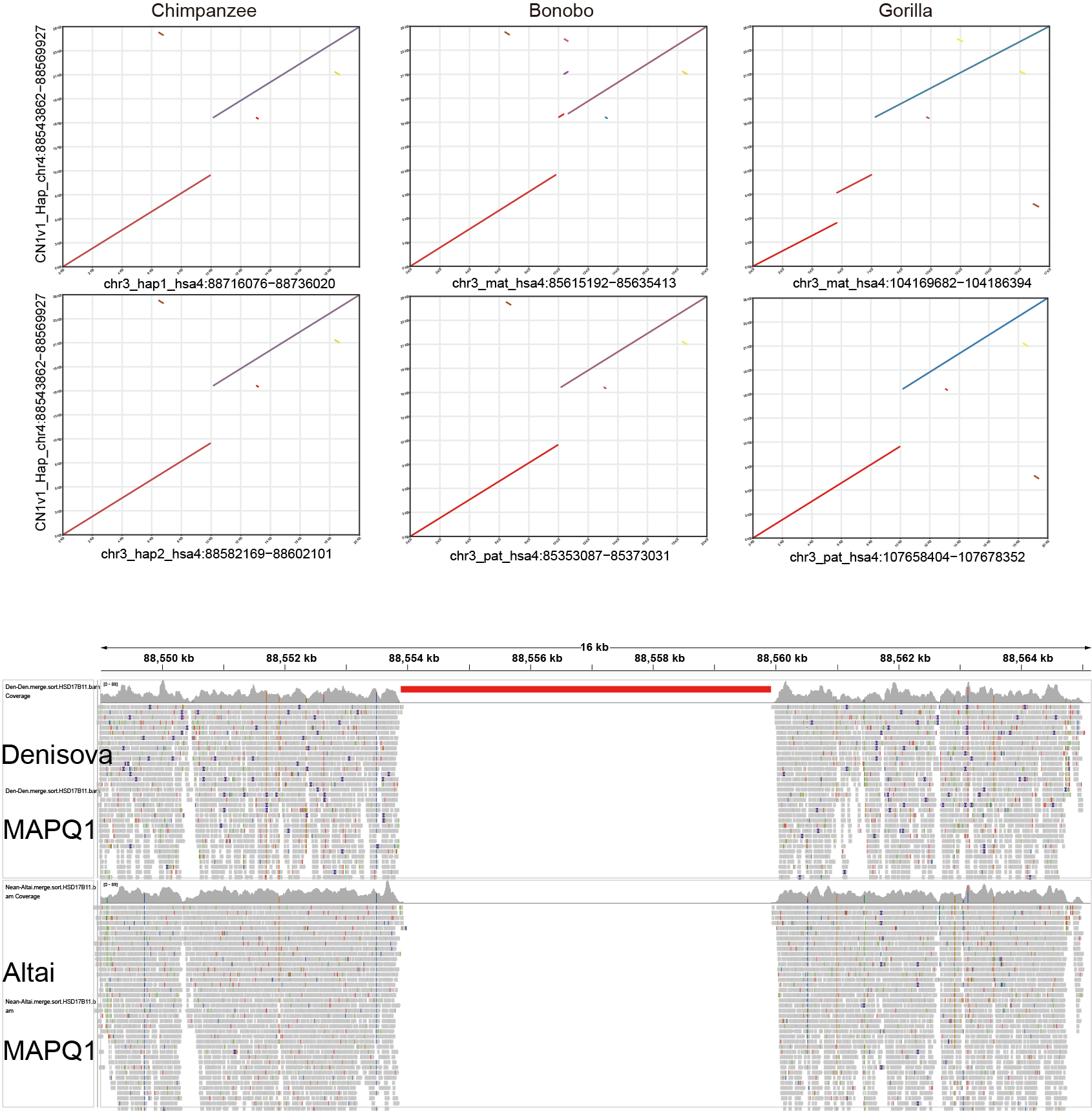

**Supplementary Fig. 37 | Absence of the LINE-1 insertion in *HSD17B11* across great apes and Neanderthal archaic genomes.** Haplotype assemblies for chimpanzee, bonobo and gorilla are from T2T-Primates project (https://github.com/marbl/Primates). The lower panel displays short-read alignments from Denisovan and Altai Neanderthal genomes to T2T-CHM13, with the red bar denoting the LINE-1 insertion.

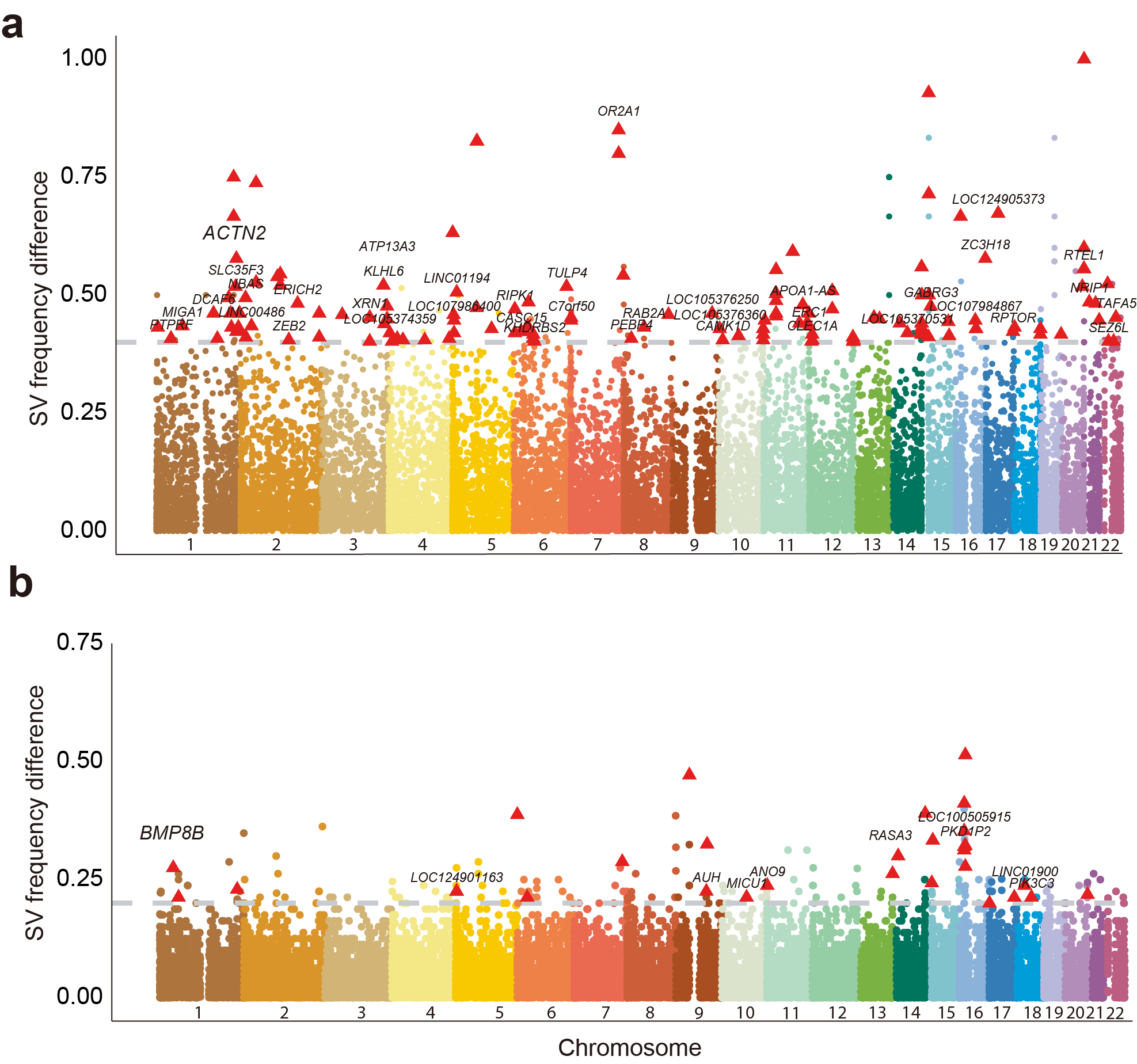

**Supplementary Fig. 38 | SV frequency differences within EAS populations. a**, Han populations (CHA-S, CHA-E, CHA-C and CHA-N; *n* = 164) *versus* Tibetan population (CZA; *n* = 16). **b**, Northern Han (CHA-N and CHA-NE; *n* = 80) *versus* Southern Han (CHA-S and CHA-E; *n* = 84). Dots represent absolute SV frequency differences between compared populations at each locus. Gray dashed lines indicate the corresponding top 1% significance thresholds. Triangles mark gene-associated SVs with significant frequency differences.

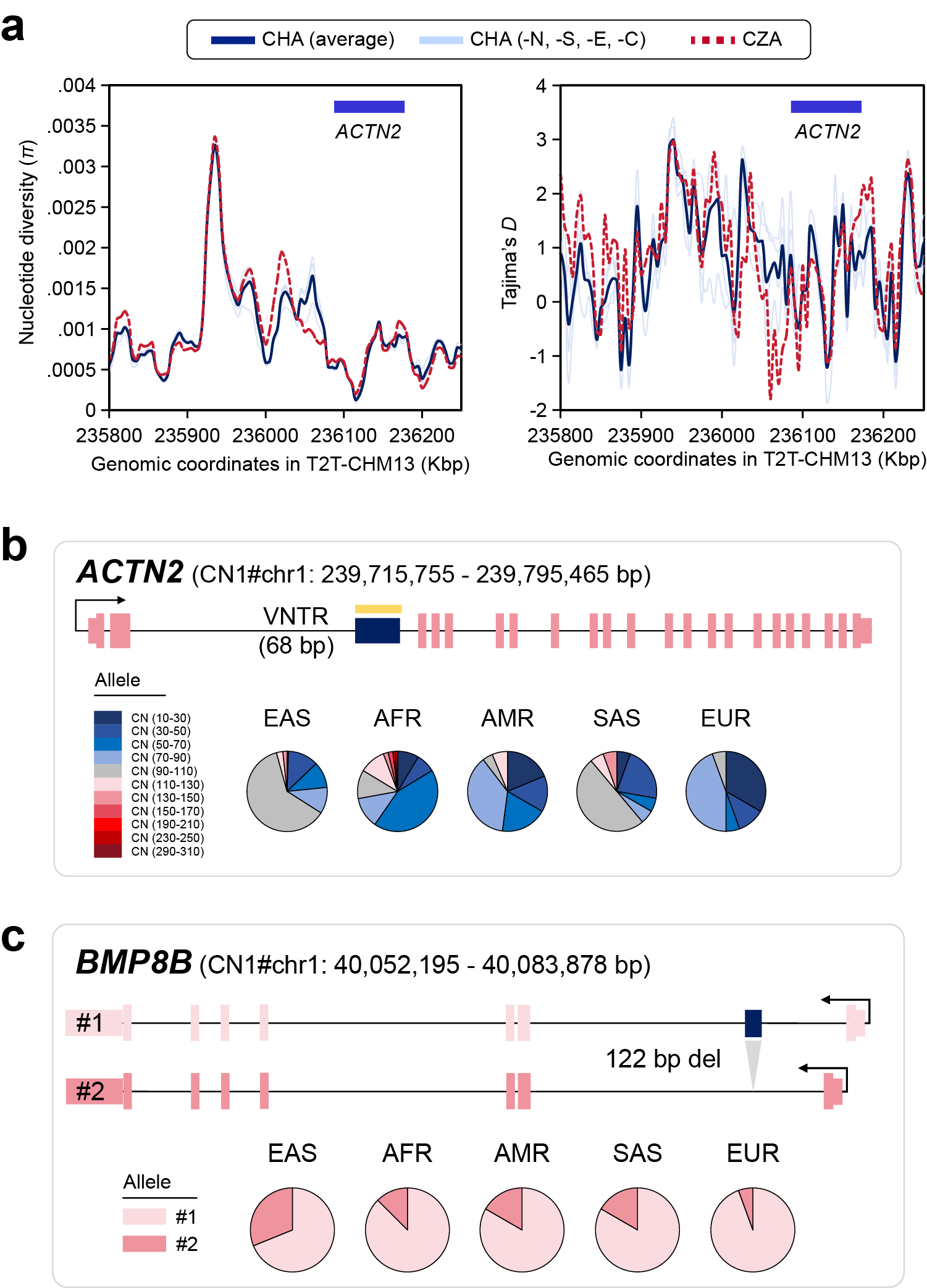

**Supplementary Fig. 39 | Selection signatures and allele frequency of SVs in *ACTN2* and *BMP8B* across superpopulations.** **a**, Nucleotide diversity (*π*) and Tajima’s *D* for Han (CHA) and Tibetan (CZA) populations around the genomic position of *ACTN2*. Tajima’s *D* 95% confidence interval is [-1.791, 1.984] for population size of *n* = 15 (Tajima, 1989, *Genetics*). **b**, Allele frequency of the *ACTN2* 68-bp VNTR copy number across superpopulations. **c**, Allele frequency of the *BMP8B* deletion among superpopulations.

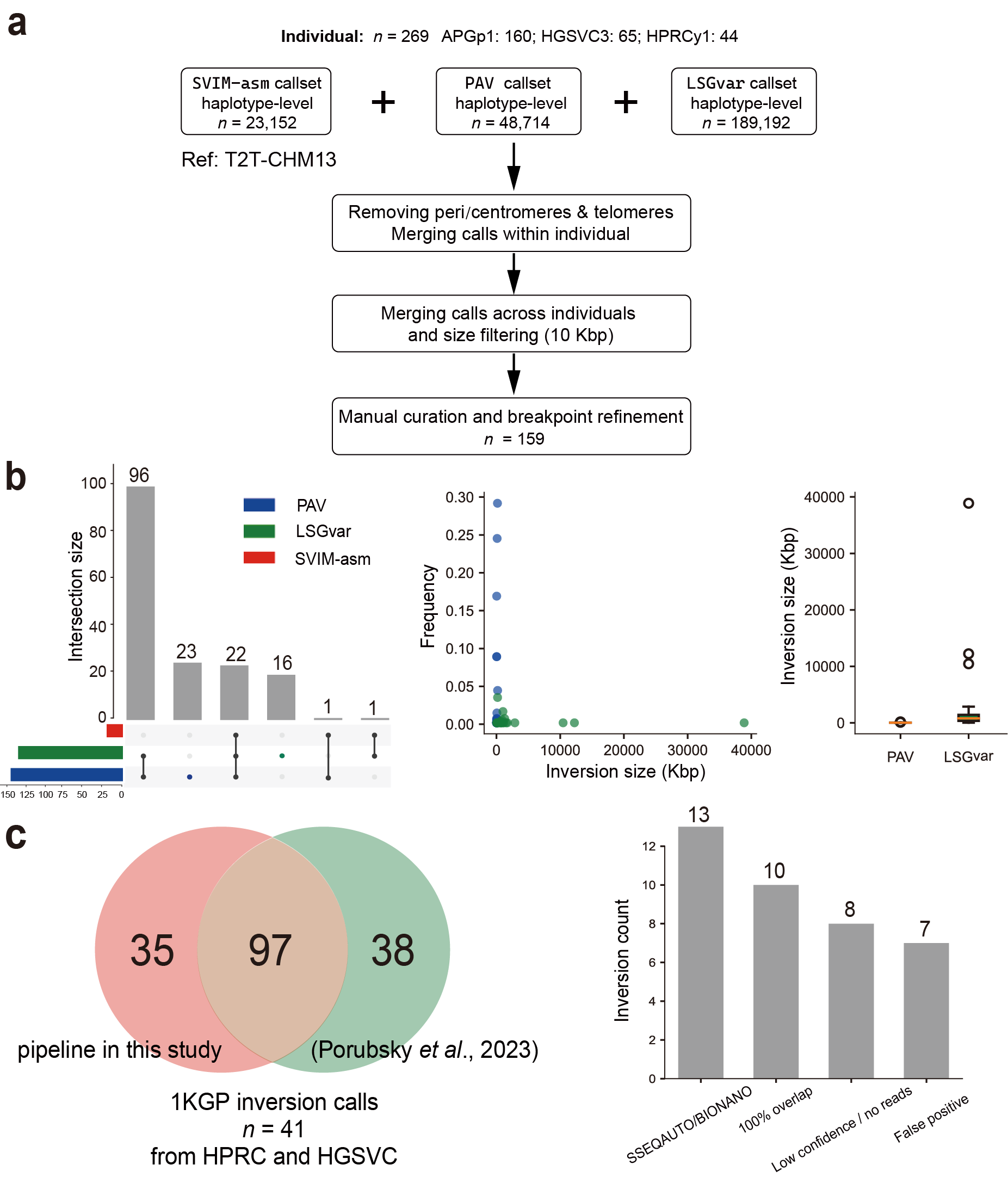

**Supplementary Fig. 40 | Large inversion calling and benchmarking in human genomes. a**, Flowchart illustrating the generation of a unified inversion call set (≥10 Kbp) on chromosome arms relative to the reference assembly T2T-CHM13 (*n* = 159). **b**, Upset plot depicting the number of inversions uniquely identified by each method: PAV, SVIM-asm, and LSGvar. The scatter plot and boxplot illustrate the length and frequency distribution of inversions uniquely called by PAV or LSGvar. **c**, Left: Venn diagram comparing our calls with a public dataset for 41 unrelated human genomes (Porubsky *et al*., 2023, *Genome Biology*). Right: Bar plots summarizing the reasons for undetected inversions in the previous study. Benchmarking revealed that 71.9% (97/135) of previously reported inversions are successfully identified, with 35 novel inversions discovered. Most undetected inversions are attributed to technological limitations (e.g. Strand-seq: chr11:1751546-1768606; Bionano: chr17:19045523-19102543 ) or low-confidence calls, or other factors in the prior studies.

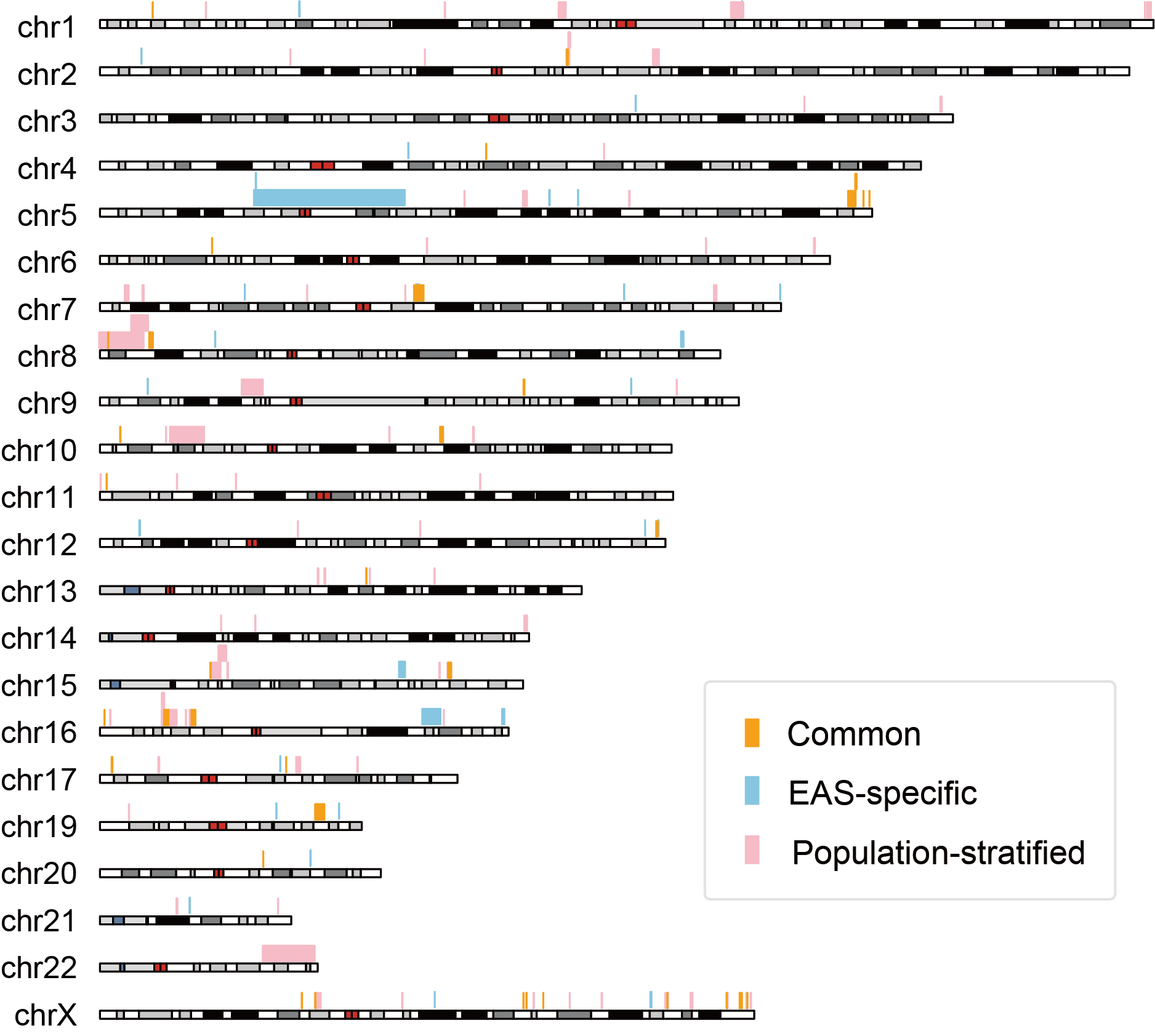

**Supplementary Fig. 41 | Large inversions on autosomes and chromosome X across global human genomes.** Inversions nested within larger inversions are staggered in the plots. EAS specific and population-stratified inversions are highlighted in blue and red, respectively. We determine the inversions with population stratification between EAS and non-EAS populations by testing the significance of their inversion frequency difference using two-tailed Fisher’s exact test. We note that all inferred population-level patterns should be interpreted with caution, owing to limited high-quality assemblies available for non-EAS populations.

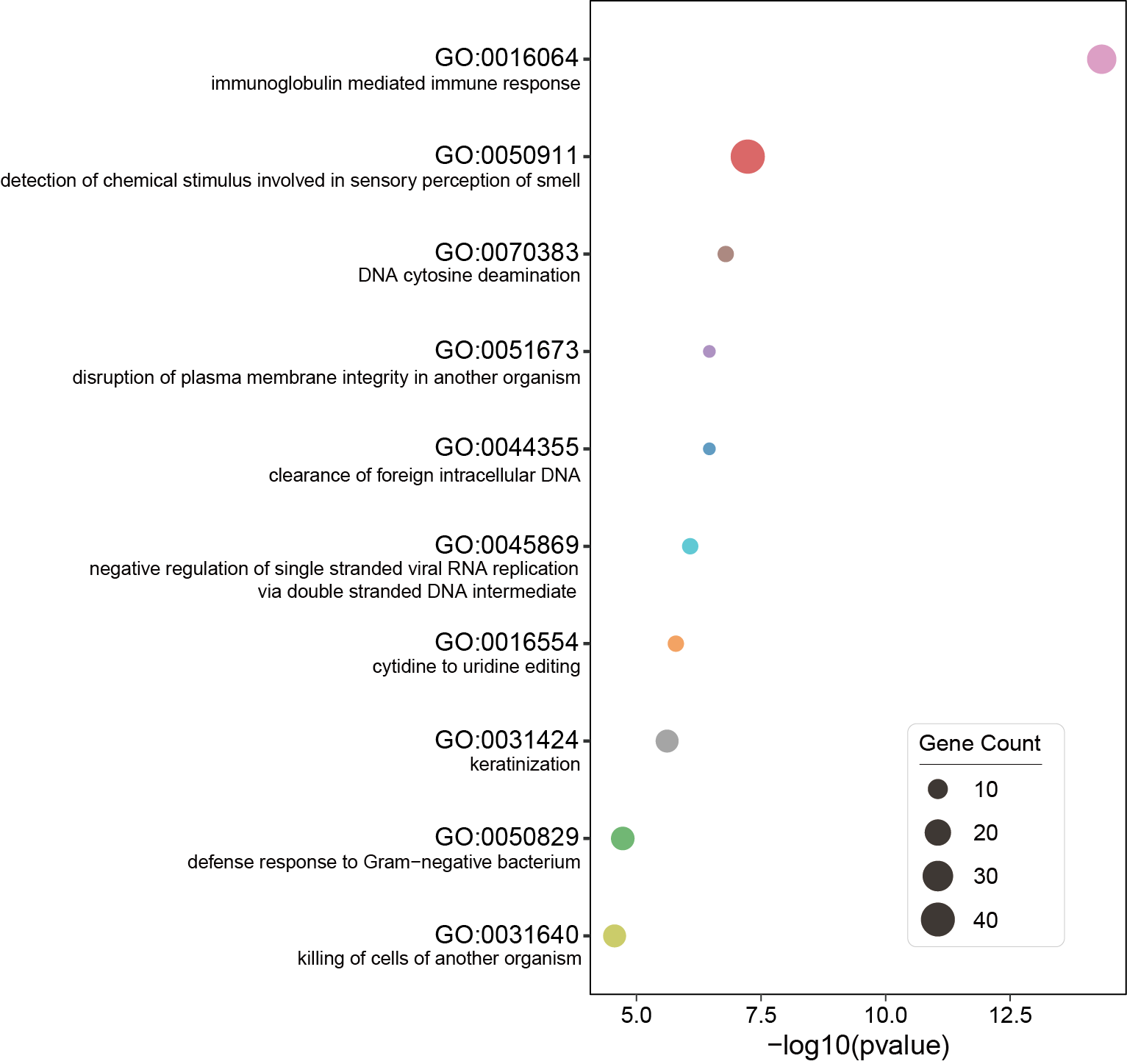

**Supplementary Fig. 42 | GO biological pathway enrichment of genes within large inversions and their 50-kbp flanking regions**.

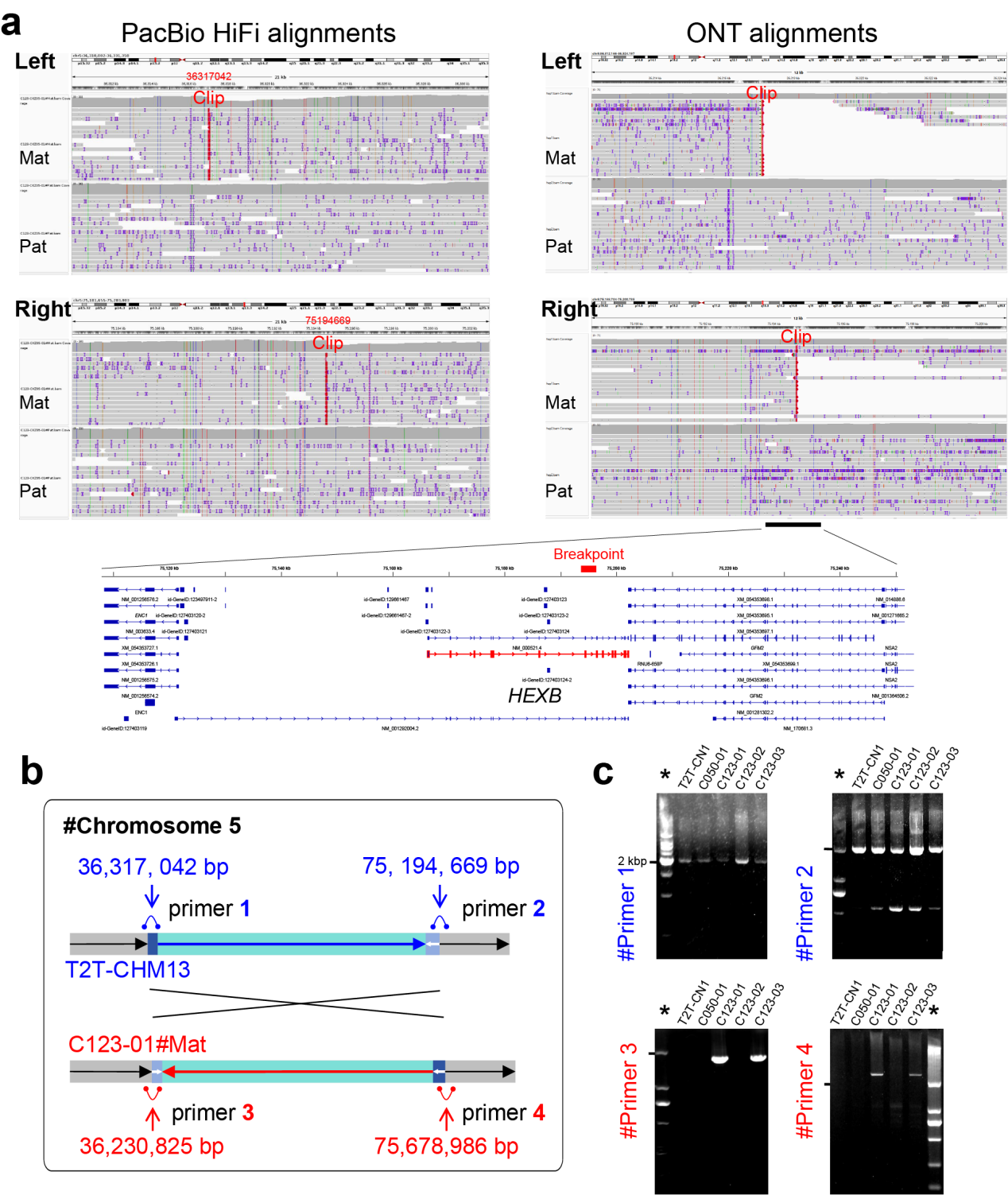

**Supplementary Fig. 43 | Validation of 5p13-5q13 ultra-large pericentric inversion.** **a**, Alignments of phased long reads (PacBio HiFi and ONT) to the T2T-CHM13 reference assembly around the breakpoints, with detailed clipping positions displayed. Gene-related annotation tracks are shown beneath the alignments. **b**, Schematic of PCR primer design for the reference (T2T-CHM13) and inverted alleles (C123-CKZ01-01#Mat). **c**, PCR products from five genomes using four primers. Molecular weight markers are indicated with an asterisk. Detailed PCR results are provided in Supplementary Table 19.

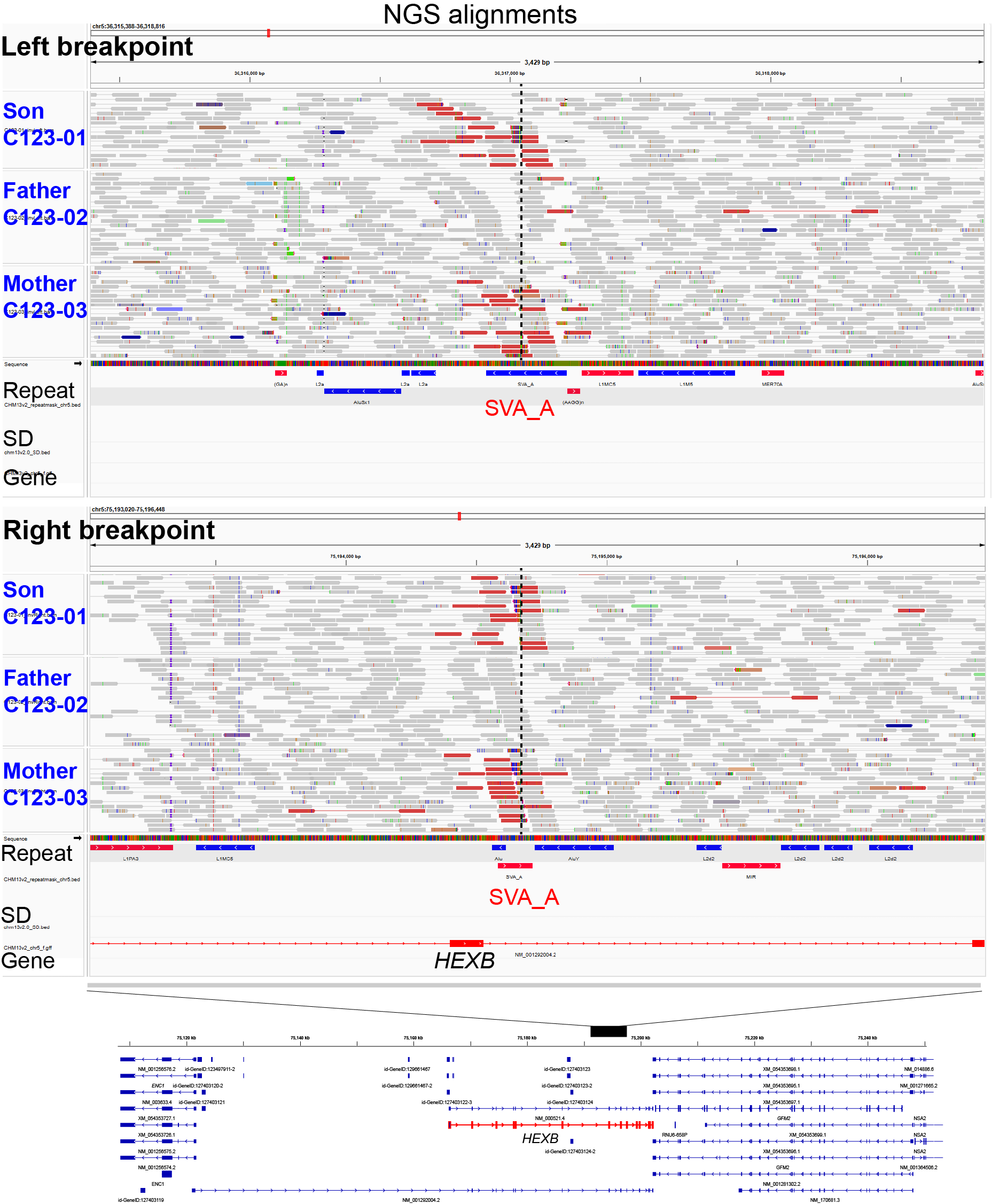

**Supplementary Fig. 44 | NGS read alignments of the C123 family around the two breakpoints of the 5p13.2-5q13.3 ultra-large inversion.** PCR-free paired-end short reads from C123-01 (child), C123-02 (father) and C123-03 (mother) are aligned and visualized in IGV. Read pairs with abnormal insert sizes (>10 Mbp) are highlighted in red. Repeat elements, segmental duplications (SDs) and gene annotations are detailed at the bottom.

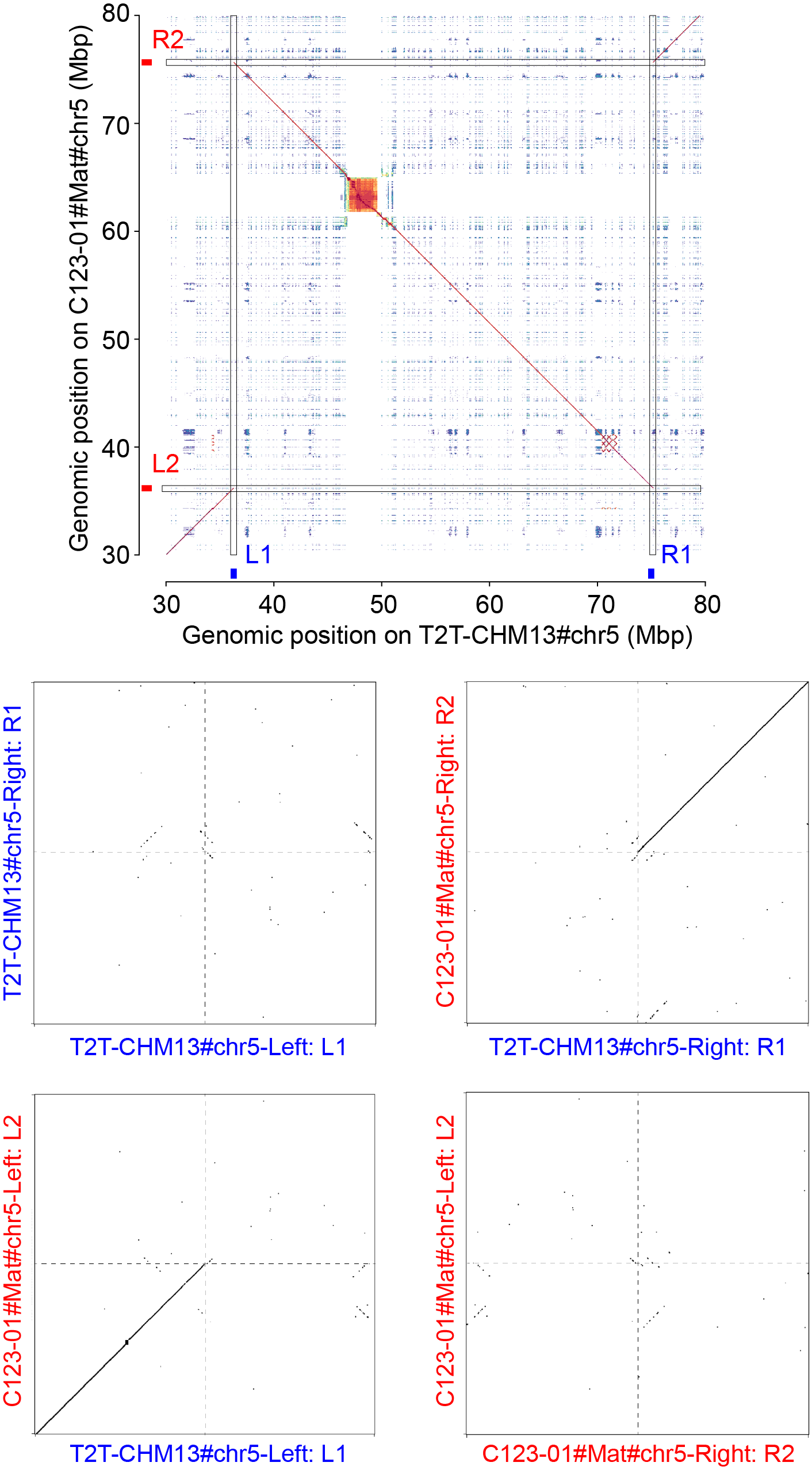

**Supplementary Fig. 45 | Sequence alignment between C123-01#Mat and T2T-CHM13 around the ultra-large inversion at 5p13.2-5q13.3.** Zoomed dotplots illustrate a pair of inverted repeat around the breakpoints in T2T-CHM13 (blue) and C123-01#Mat (red).

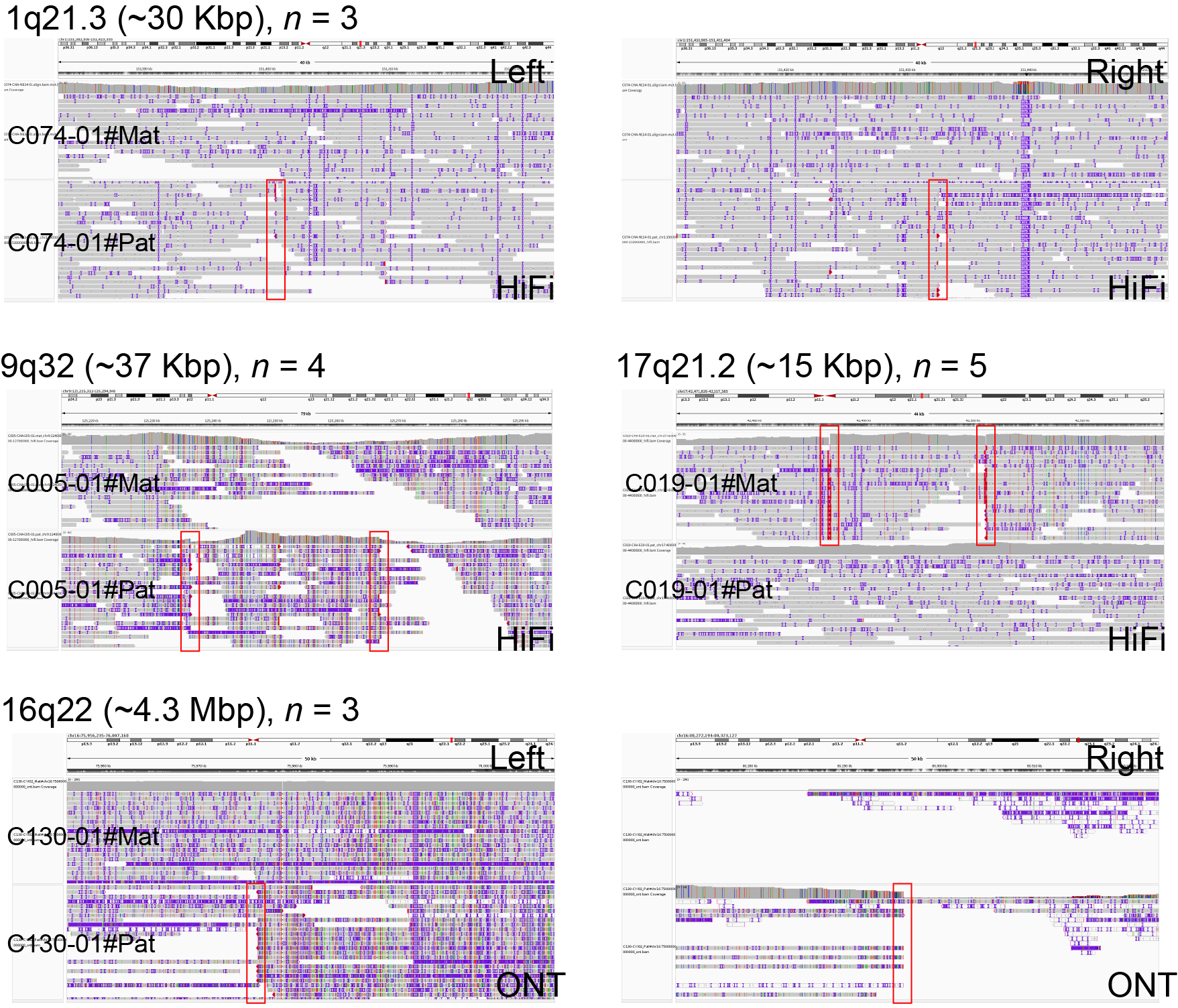

**Supplementary Fig. 46 | Validation of EAS-specific large inversions by long-read alignments.** Inversions observed in at least three assemblies are shown, with phased PacBio HiFi or ONT reads aligned to the T2T-CHM13 reference assembly. Red boxes denote genomic locations of inversion breakpoints.

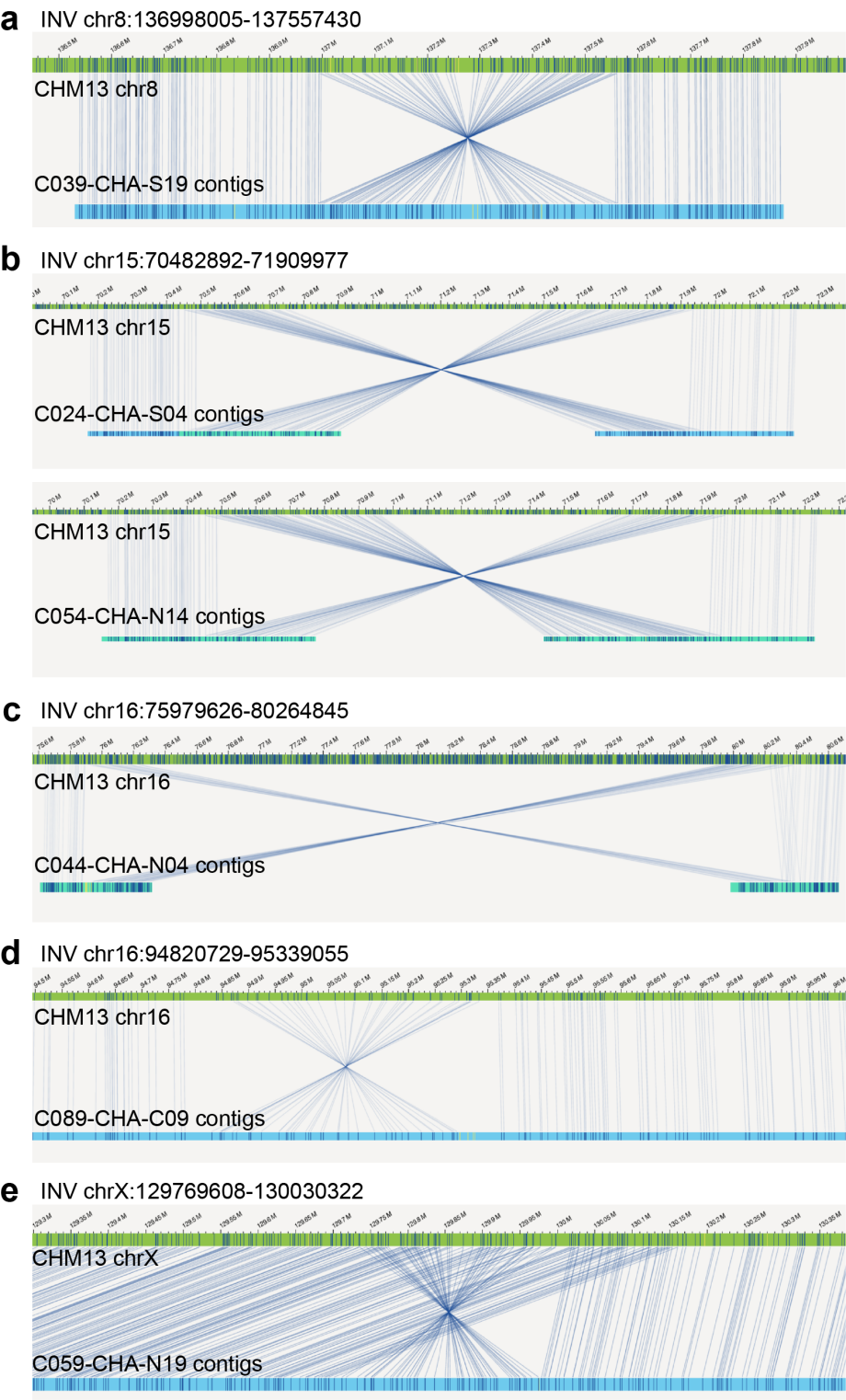

**Supplementary Fig. 47 | Validation of EAS-specific large inversions (>100 Kbp) by Bionano optical maps.** Five >100-Kbp inversions are visualized in Bionano Access, including these at 8q24.23 (**a**), 15q24.1 (**b**), 16q22.1 (**c**), 16q24.3 (**d**), and Xq26.2 (**e**). The remaining three >100-Kbp inversions (5p13.2-5q13.3, 7q11.23, 12p13.31) were not investigated due to insufficient DNA samples for Bionano data generation.

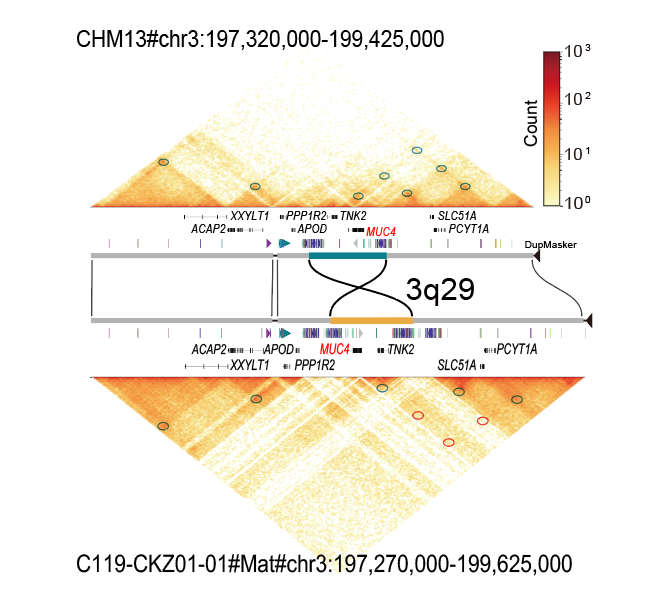

**Supplementary Fig. 48 | Topological differences at 3q29 between C119-01#Mat and T2T-CHM13. H**eatmap colors represent the intensity of contacts between proximal genomic regions, ranging from low (yellow) to high (red) interaction frequency. Regions with different contact levels between two matrices are highlighted by red circles, indicating potential chromatin loop alterations.

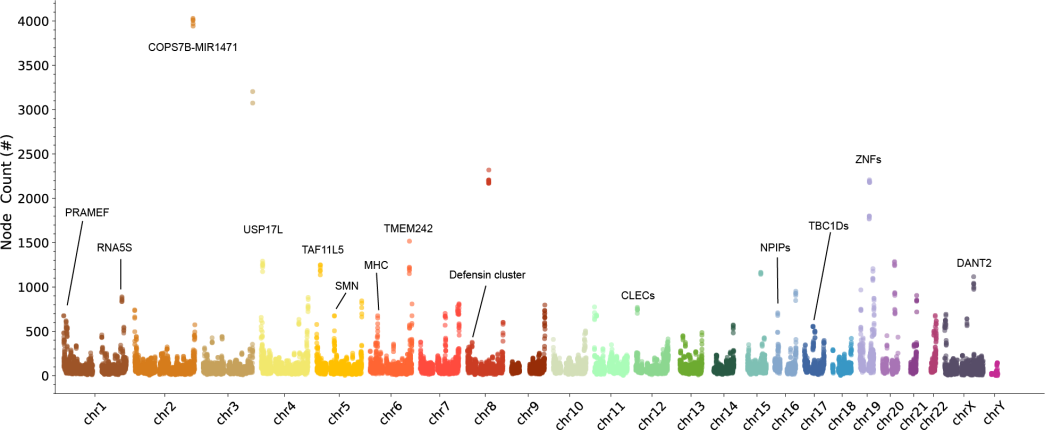

**Supplementary Fig. 49 | Complexity of human pangenome graph.** The pangenome graph here is constructed using Minigraph, including assemblies from APGp1, HPRCy1, and HGSVC3. Nodes are counted in 500-kbp windows, with high-complexity regions annotated.

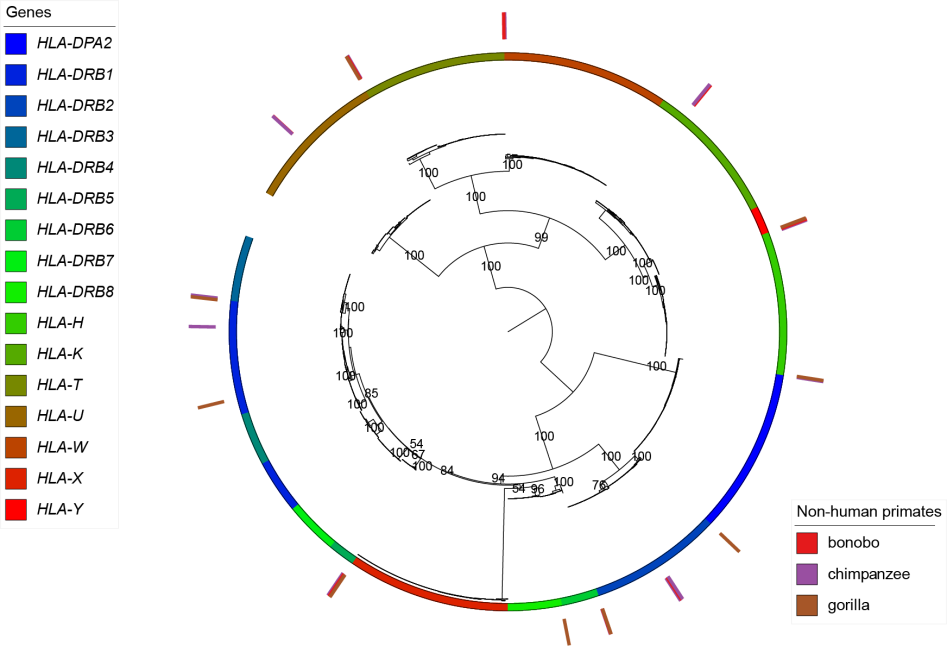

**Supplementary Fig. 50 | A maximum-likelihood phylogenetic tree of all annotated multi-copy HLA genes.** Bootstrap values from 1000 replicates are shown at major branches. HLA genes from three non-human primates (chimpanzee, bonobo, and gorilla) are included as outgroups for each gene, with *HLA-X*, *HLA-Y* and *HLA-DRB4* unannotated in the T2T primate genomes (https://github.com/marbl/Primates).

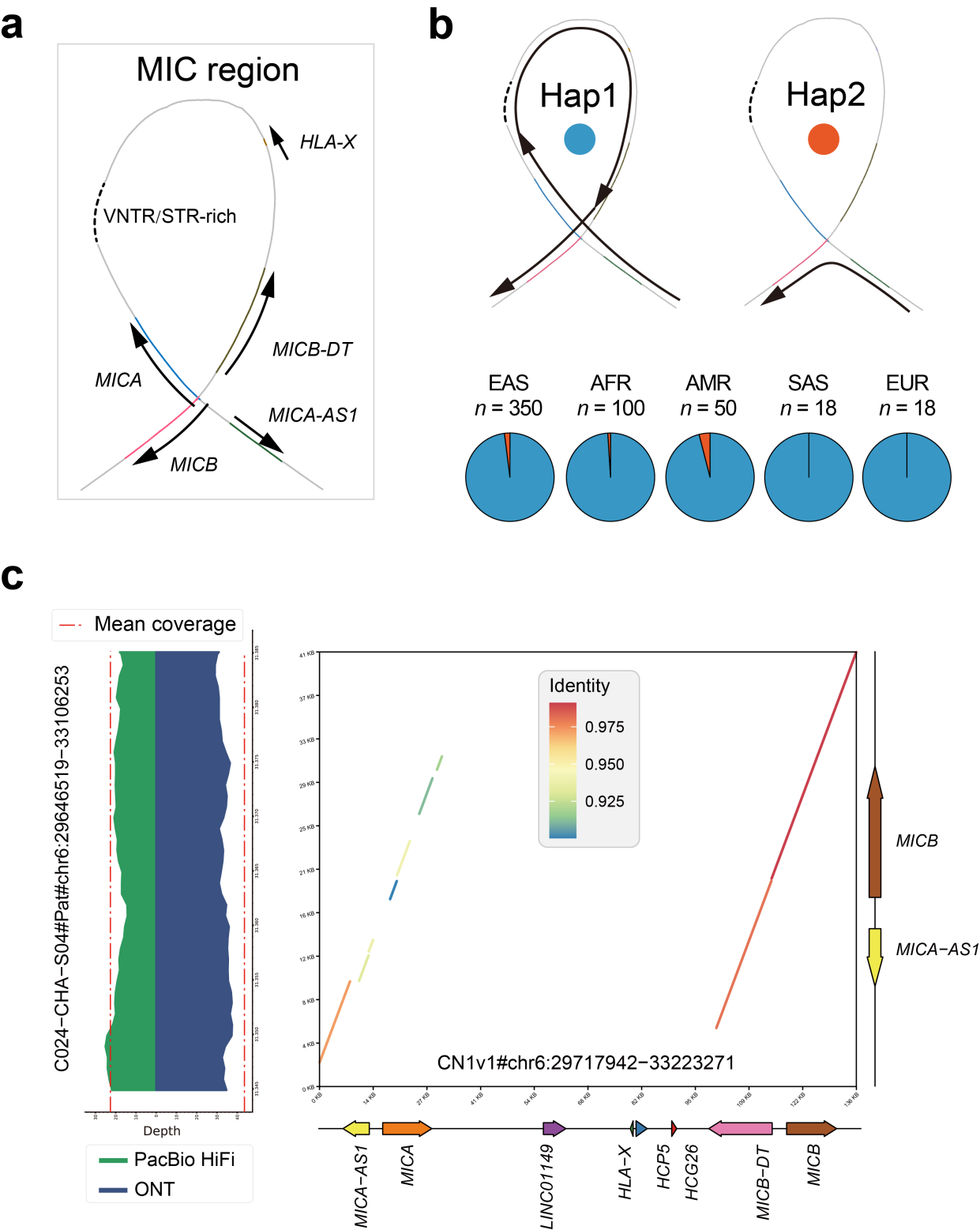

**Supplementary Fig. 51 | Structural haplotypes of human MIC sequences and their frequency across five super populations.** **a**, Local MC graph of the human MIC region. Genes are depicted as colored lines with orientation arrows. A VNTR-rich region is simplified by a dashed line due to extreme complexity in the MC graph. **b**, Two structural haplotypes of the MIC region and their frequency in five super populations. **c**, Local assembly validation of the MIC deletion allele.

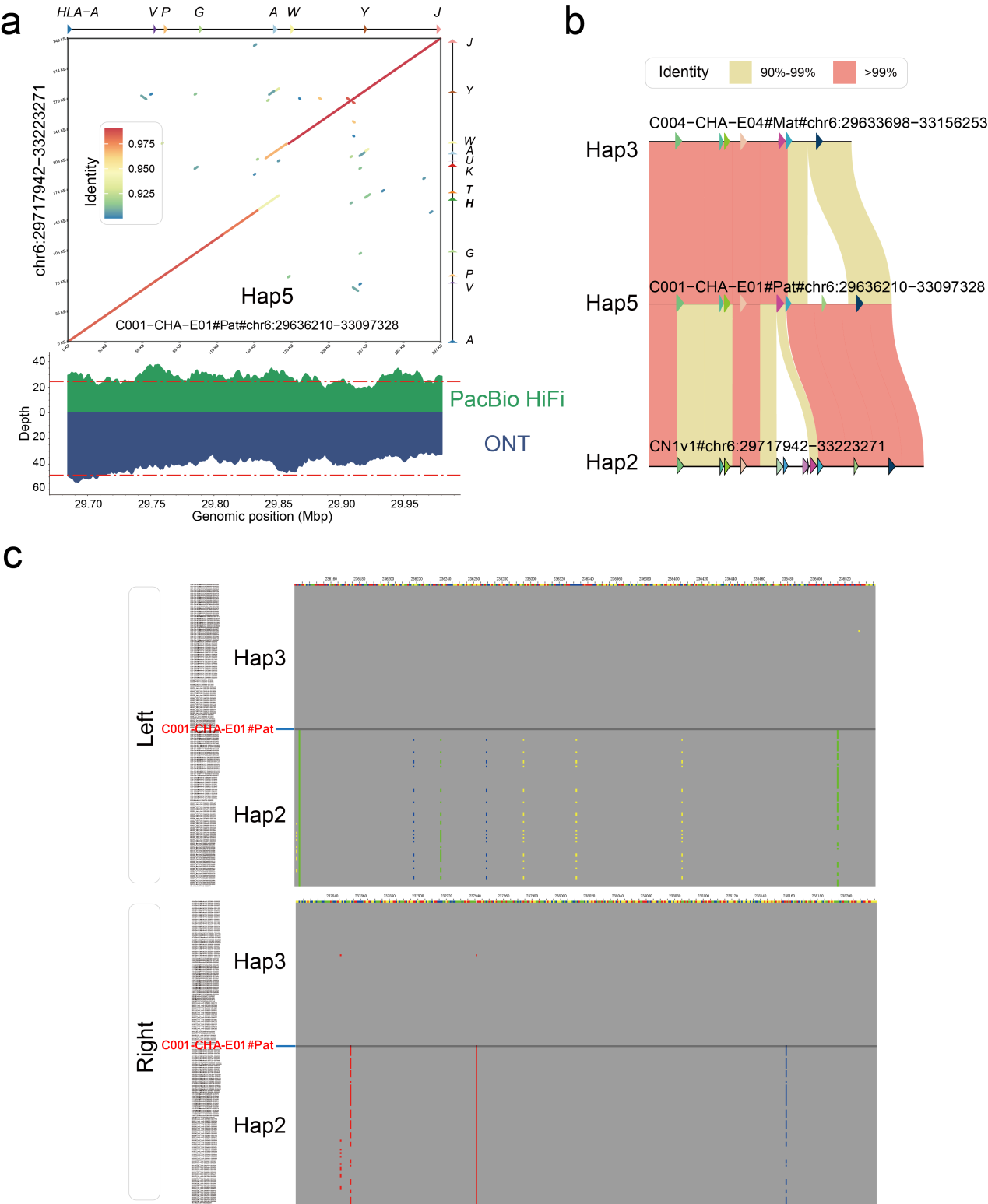

**Supplementary Fig. 52 | Recombination origin of EAS-specific haplotype Hap5 in the HLA-A region. a**, Sequence synteny between Hap5 (C001-CHA-E01-01#Pat) and T2T-CN1, and assembly validation by examining curated alignments of PacBio HiFi and ONT reads. **b**, Sequence identity among Hap5, Hap2 and Hap3, revealing the recombination origin of Hap5. **c**, Sequence alignments showing shared variations in the HLA-A region between Hap5 (C001-CHA-E01-01#Pat) and Hap3 on the left and Hap2 on the right, respectively.

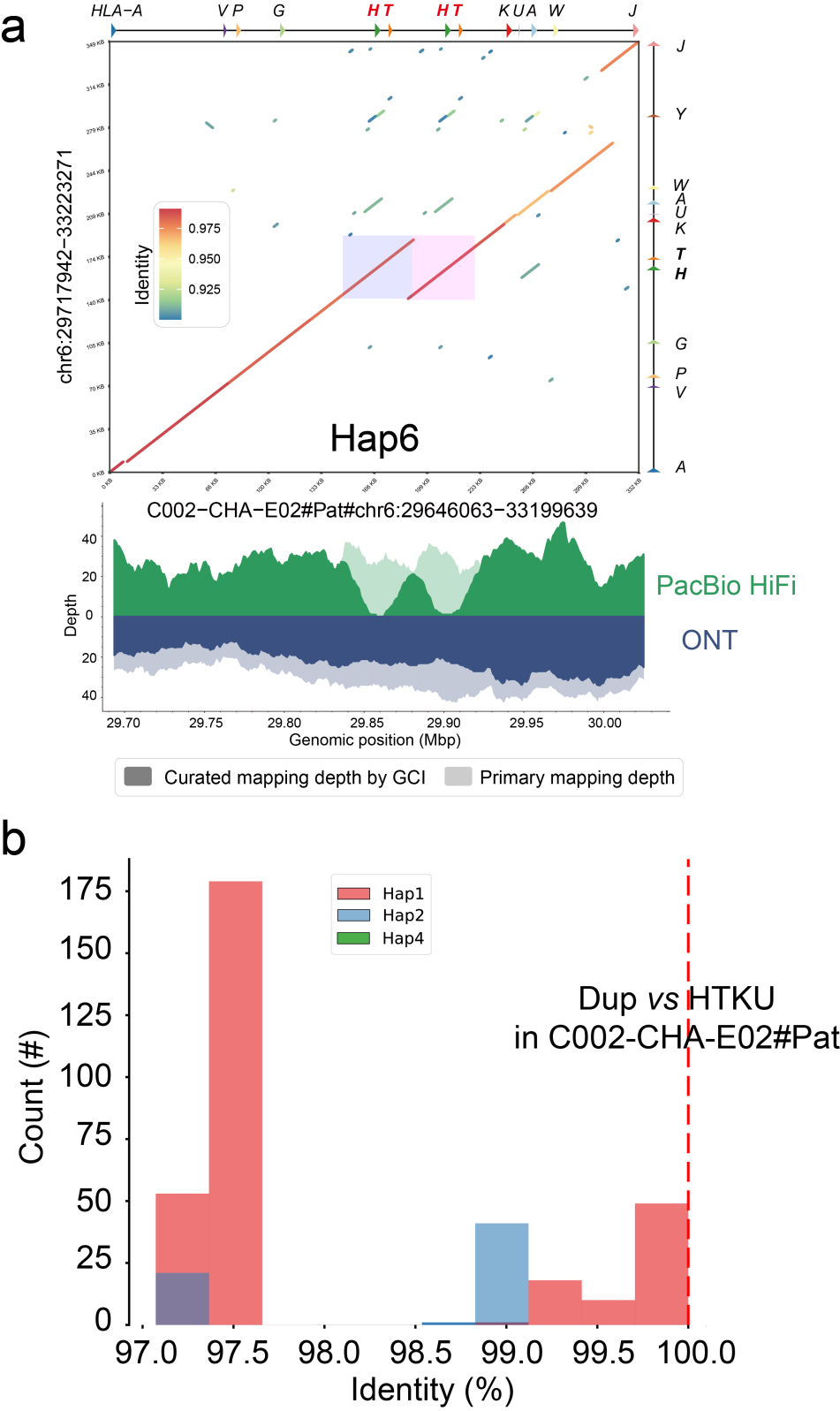

**Supplementary Fig. 53 | Recent segmental duplication in a novel HLA-A haplotype Hap6. a**, Sequence synteny between Hap6 (C002-CHA-E02-01#Pat) and T2T-CN1, and assembly validation by examining curated primary alignments of PacBio HiFi and ONT reads. The near-identical nature of the two copies induces two coverage valleys in PacBio HiFi alignments, as detected by GCI. **b**, Sequence identity between the HTKU sequence of C002-CHA-E02-01#pat assembly and its duplicate (vertical red dashed line), compared with other HTKU sequences from Hap1, Hap2 and Hap4 assemblies.

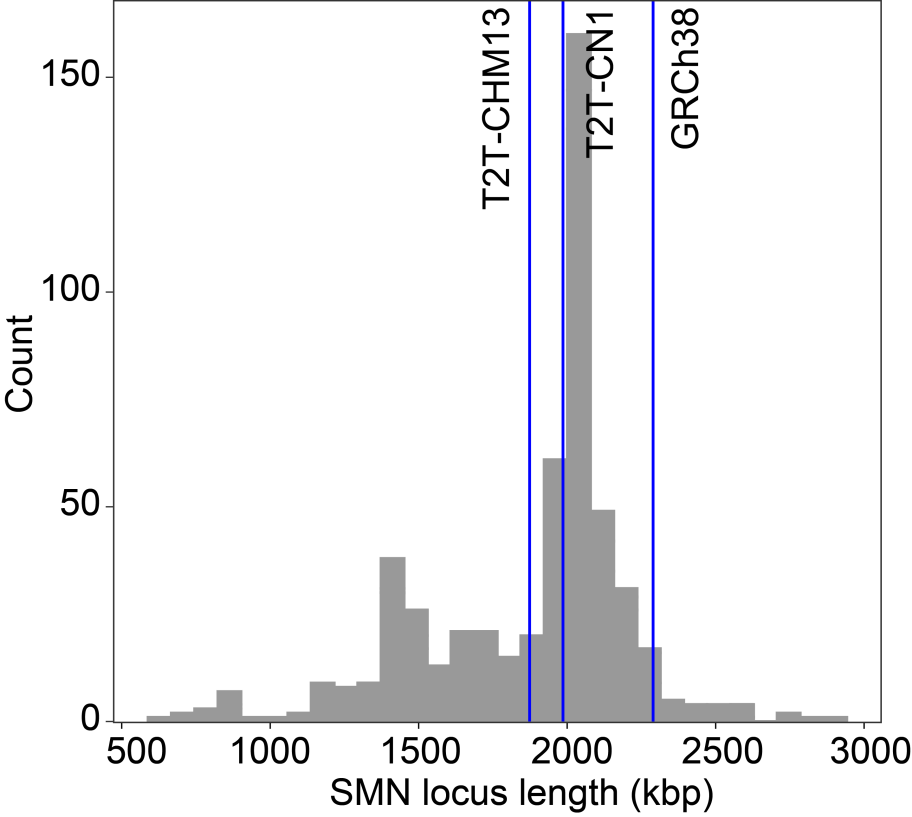

**Supplementary Fig. 54 | Genomic lengths of SMN sequences (from *RAD17* to *MCCC2*) across global human genome assemblies.** The three vertical blue lines denote the SMN locus lengths in T2T-CHM13 (left), T2T-CN1 (middle) and GRCh38 (right).

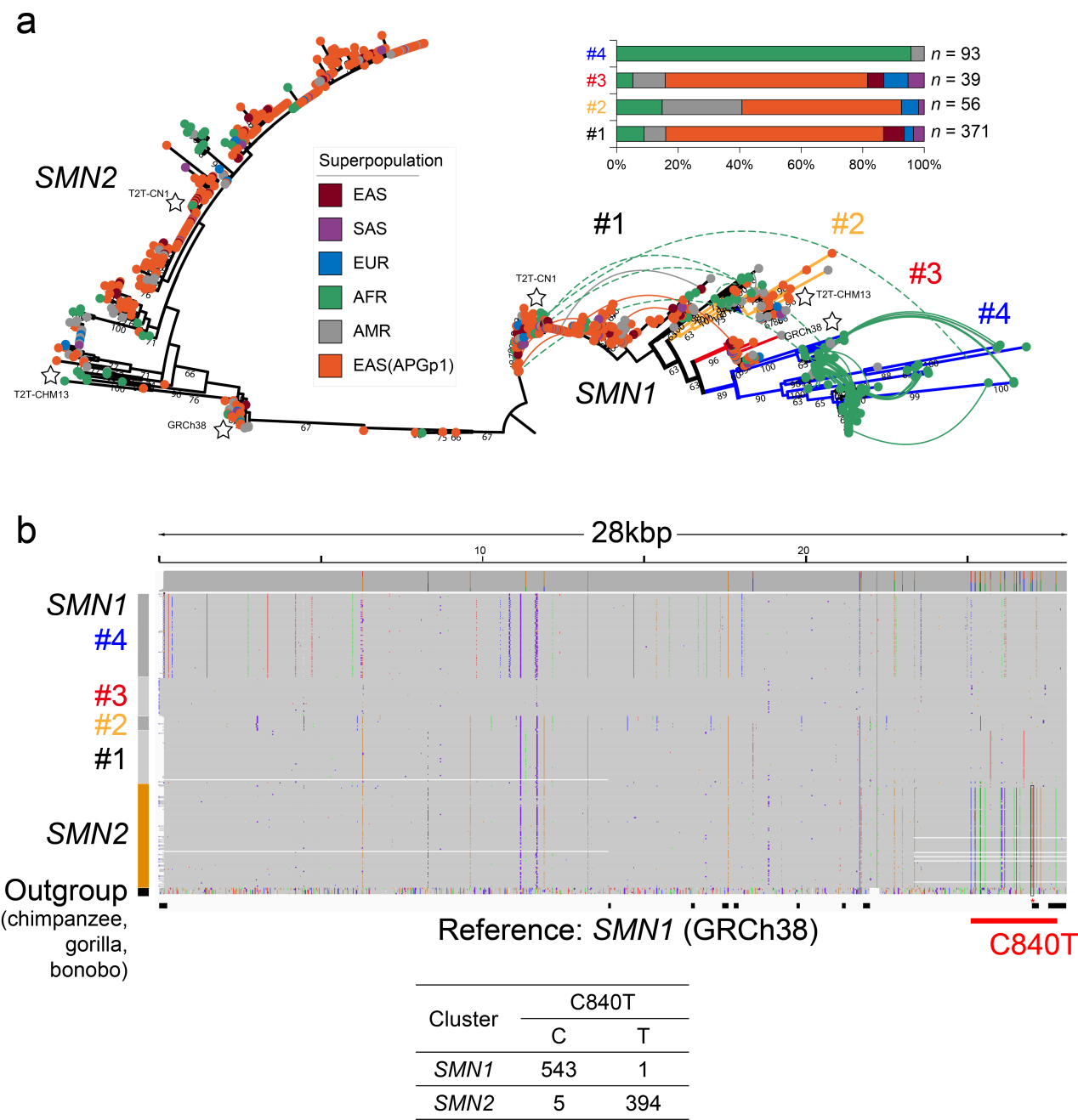

**Supplementary Fig. 55 | Genomic differentiation between *SMN1* and *SMN2* genes. a**, A maximum-likelihood phylogenetic tree encompassing all annotated SMN genes. Stars denote the locations of SMN genes from three reference genomes (T2T-CHM13, T2T-CN1 and GRCh38) in the tree. Edges connecting *SMN1* leaf nodes indicate cases where two copies of *SMN1* genes are present per haploid assembly. Four clusters of *SMN1* genes are defined based on the phylogeny. Bootstrap values less than 60 are not shown. **b**, Alignments of SMN gene sequences against the *SMN1* sequence in GRCh38. Non-redundant unique sequences are used. The divergent region between SMN1 and SMN2 is delineated by a bold red line. The diagnostic causative mutation C840T, which serves as the primary marker distinguishing *SMN1* and *SMN2*, is highlighted, alongside its observed allele frequencies.

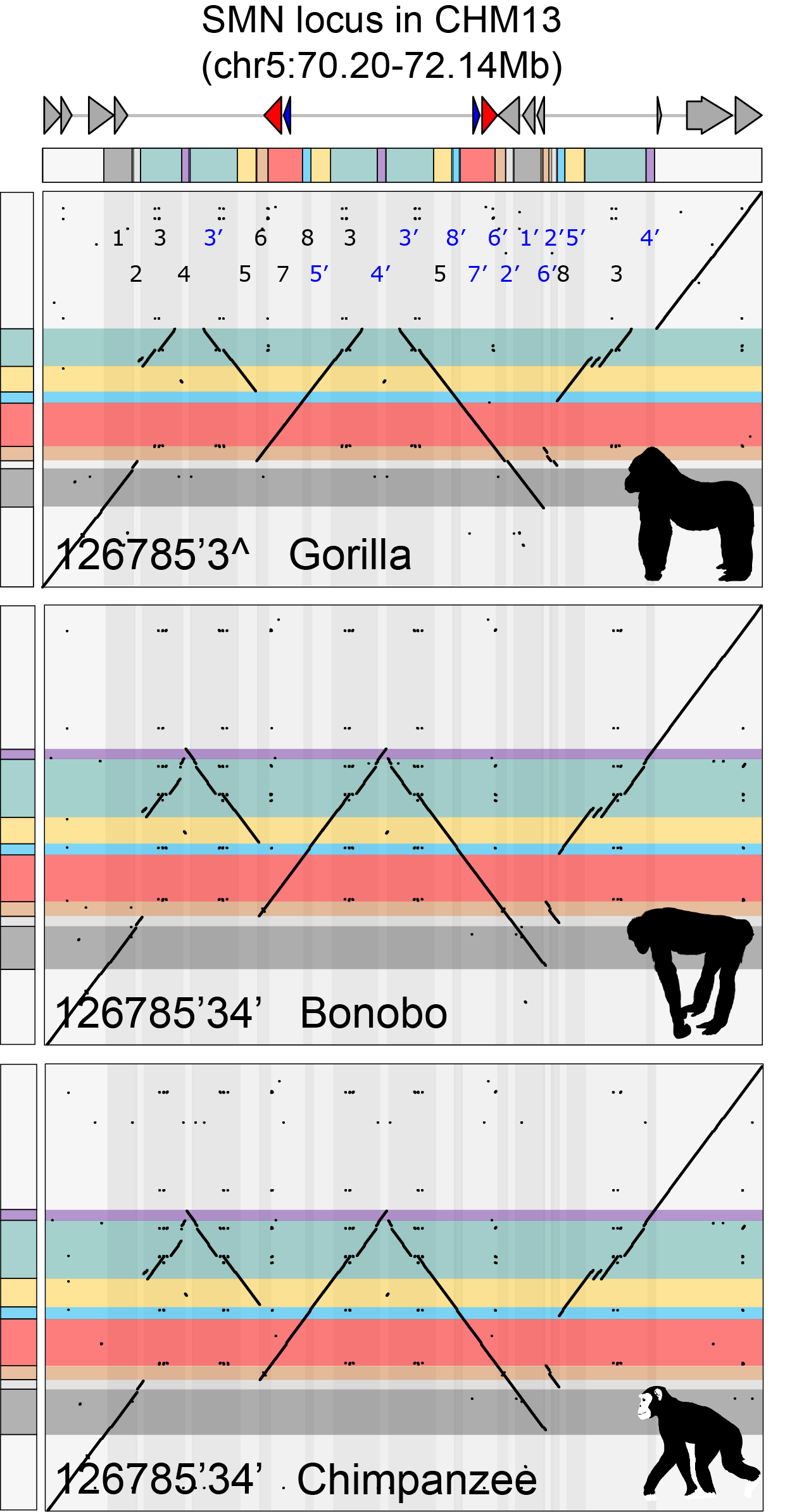

**Supplementary Fig. 56 | Decomposition of SMN loci in non-human primate genomes.** Eight principal blocks are color-coded for distinction. Phased genome assemblies of chimpanzee (*Pan troglodytes*), bonobo (*Pan paniscus*) and gorilla (*Gorilla gorilla*) are from T2T-primates project (v2.0, https://github.com/marbl/Primates), with maternal haplotype assemblies of the three species shown. Numerical strings denote block arrangements across the SMN sequences. The prime symbol (’) represents the reverse strand, and the caret symbol (^) indicates a partial block, relative to the defined reference block.

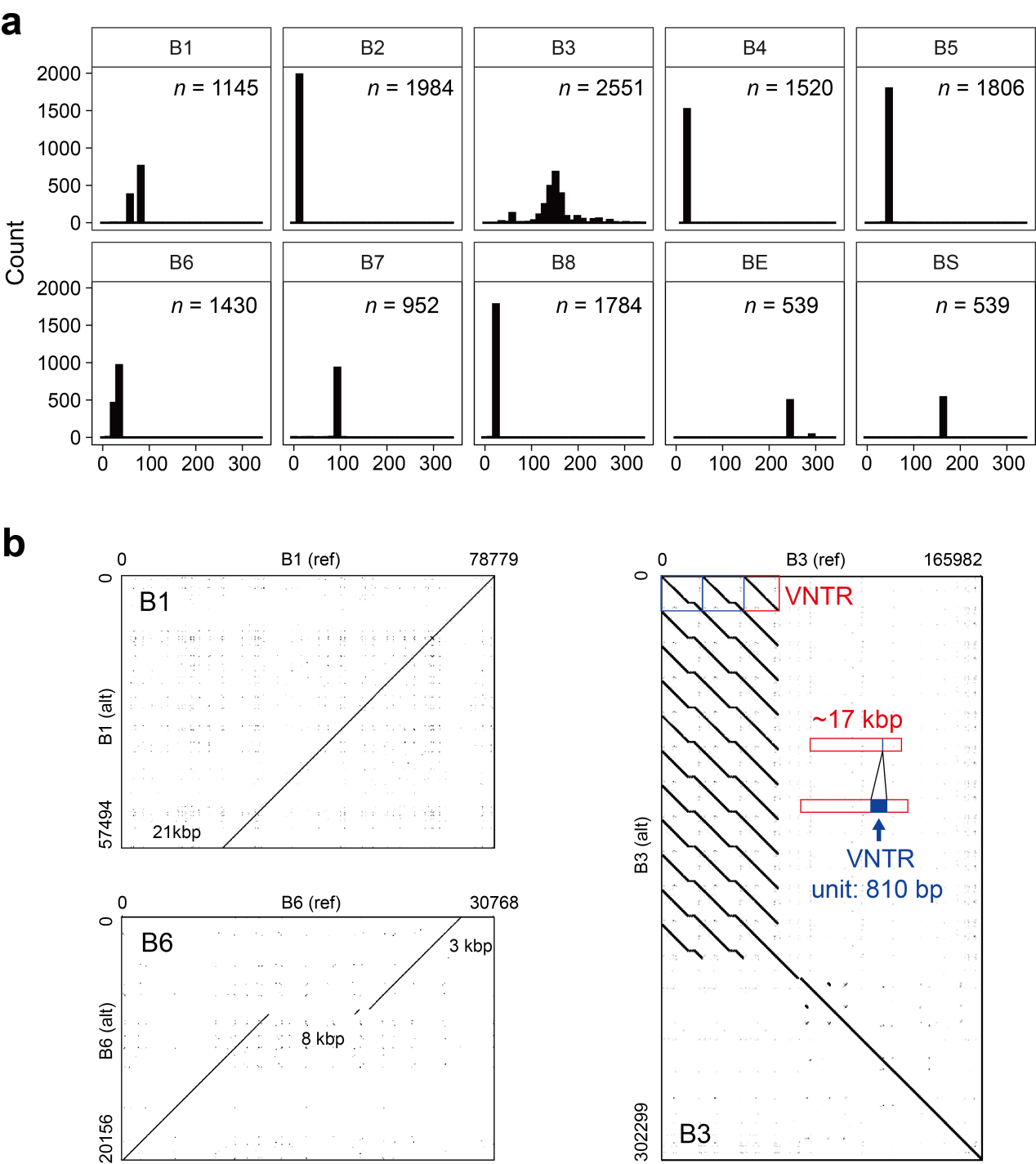

**Supplementary Fig. 57 | Length distribution and variants of SMN blocks. a**, size distribution of all decomposed SMN blocks across all human genome assemblies. BS, left flank of SMN locus; BE, right flank. **b**, Nucleotide alignment plots between reference and alternative sequences for Blocks B1, B3 and B6. B1 and B6 each exhibit two length clusters, corresponding to two subtypes. B1 has an alternative unit that carries an ~21-kbp deletion relative to the T2T-CHM13 reference unit. B6 variant has two deletions of ~3 kbp and ~8 kbp, respectively. B3 shows nested VNTRs of 17-kbp and 810-bp units. The plots are generated by geopard, with a word length of 25 and window size of 0.

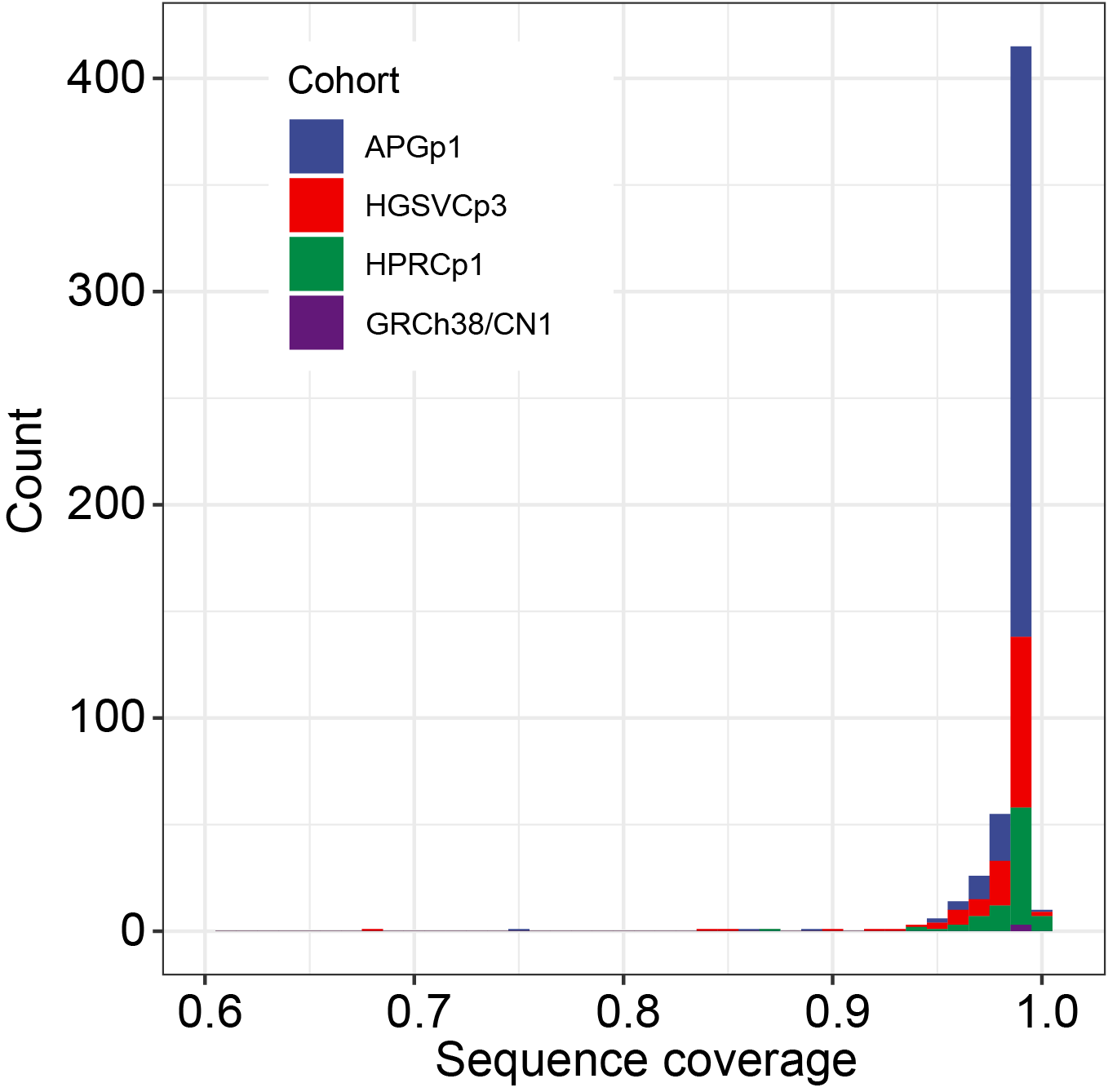

**Supplementary Fig. 58 | Sequence decomposition coverages in the SMN region.** The SMN sequence in each haplotype assembly is decomposed by aligning eight principal minimizer blocks.

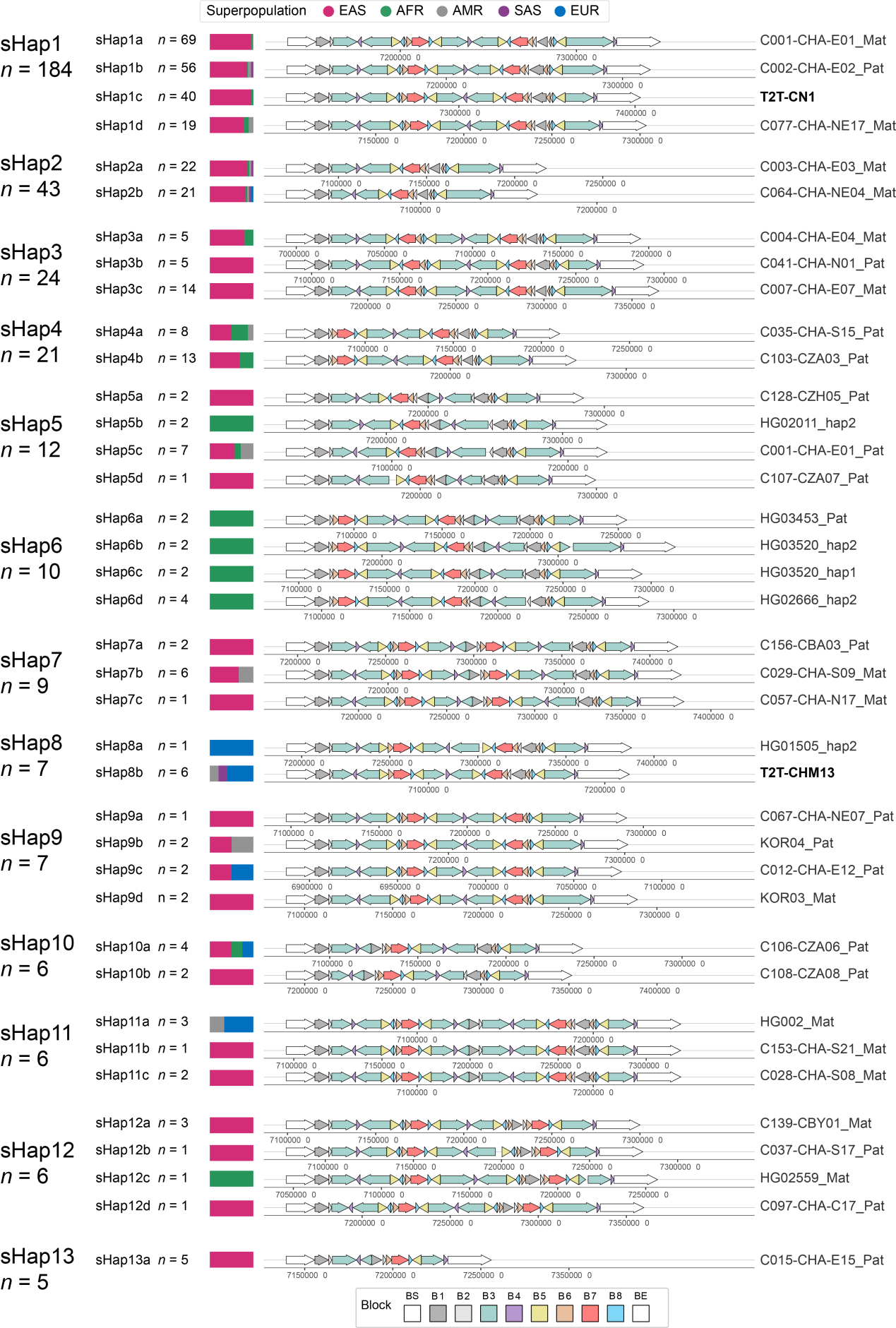

**Supplementary Fig. 59 | Structural haplotypes (sHaps) in the SMN region across human genomes and their frequency in five superpopulations.** Thirteen common sHaps (frequency > 0.01) are shown, including their sub-types with recurrent inversions within Palindrome 3-4-3’. BS, left flank; BE, right flank.

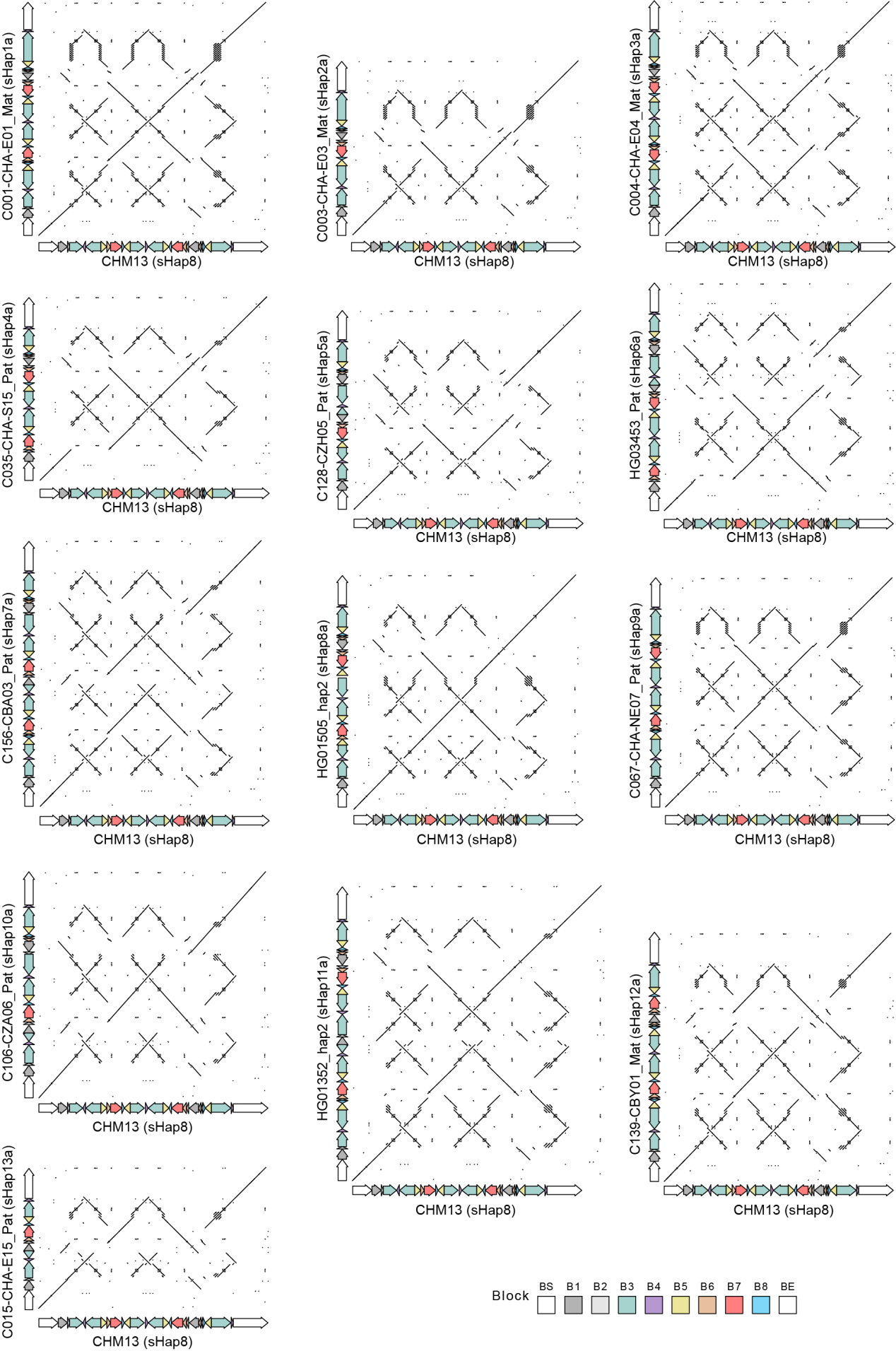

**Supplementary Fig. 60 | Nucleotide sequence alignments among SMN structural haplotypes.** The T2T-CHM13 (sHap8) is used as the reference sequence.
